# Complete functional analysis of type IV pilus components of a reemergent plant pathogen reveals neofunctionalization of paralog genes

**DOI:** 10.1101/2022.10.24.513447

**Authors:** Marcus V Merfa, Xinyu Zhu, Deepak Shantharaj, Laura M. Gomez, Eber Naranjo, Neha Potnis, Paul A. Cobine, Leonardo De La Fuente

**Affiliations:** Department of Entomology and Plant Pathology, Auburn University, Auburn, AL 36849, USA; Department of Biological Sciences, Auburn University, Auburn, AL 36849, USA

## Abstract

Type IV pilus (TFP) is a multifunctional bacterial structure involved in twitching motility, adhesion, biofilm formation, as well as natural competence. Here, by mutagenesis and functional analysis, we dissected the roles of all genes required for TFP biosynthesis and regulation in the reemergent plant pathogenic fastidious prokaryote *Xylella fastidiosa*. This xylem-limited, insect-transmitted pathogen lives constantly under flow conditions and therefore is highly dependent on TFP for host colonization. In addition, TFP-mediated natural transformation is a process that impacts genomic diversity and environmental fitness. Ten out of the thirty-eight genes analyzed were essential for movement and natural competence. Interestingly, seven sets of paralogs exist, and mutations showed opposing phenotypes, indicating evolutionary neofunctionalization of subunits within TFP. The minor pilin FimT3 was the only protein exclusively required for natural competence. We determined that FimT3 (but not the other two FimT paralogs) is a DNA receptor that is conserved among *X. fastidiosa* strains and binds DNA non-specifically via an electropositive surface. Among plant pathogens, this gene was also found in the genome of strains of the plant associated Xanthomonadaceae family. Overall, we highlight here the complex regulation of TFP in *X. fastidiosa*, providing a blueprint to understand TFP in other bacteria living under flow conditions.

## Main

Type IV pili (TFP) are retractable hair-like proteinaceous cell appendages located at one or both poles of a bacterial cell ^1^. In addition to playing fundamental roles in twitching motility and natural transformation, they also influence virulence, adhesion, and biofilm formation in many bacterial species ^2–4^. During twitching motility, TFP extend, attach to surfaces, then retract pulling cells towards the point of attachment ^1,2^. TFP also plays a central part in the DNA-uptake machinery that enables natural transformation ^3,5^. Natural transformation is a horizontal gene transfer mechanism that allows bacterial cells to take up free DNA from the environment and incorporate it into their own genomes via homologous recombination (HR) ^3,6^. The current model of natural transformation in bacteria involves binding of double-stranded DNA to the tip of TFP, which upon retraction brings this molecule to the outer-membrane surface of the cell ^4^. DNA is then transported to the periplasm in a proposed Brownian ratchet mechanism ^4,7^. Subsequently, DNA is translocated to the cytosol as a single strand, where recombination occurs if the incoming DNA shares homology with the genome of the recipient cell ^3^. Alternatively, the incoming DNA is either metabolized or used as template to repair damaged DNA ^8^.

Natural transformation enables rapid evolution by generating genetic diversity and has been involved in spreading of antibiotic resistance, adaptation to new environments and emergence of pathogens ^6,8–10^. In addition to animal pathogens ^6,9^, natural competence has been detected in two plant pathogens, *Ralstonia solanacearum* ^11^ and *Xylella fastidiosa* ^12^, both of which possess a very broad plant host range and are xylem-colonizers^13^. *X. fastidiosa* is a gram-negative fastidious prokaryote that infects and causes reemergent diseases in many economically important crops worldwide, such as grapevine, citrus, almond, peach, blueberry, and olive ^14–16^. This pathogen lives constantly under flow conditions inside the xylem of infected plants and the foregut of xylem-feeding insect vectors ^17^. The main mechanisms of virulence for *X. fastidiosa* are linked to its ability to move systemically within the xylem by TFP-mediated twitching motility, and formation of biofilm that disrupts sap flow ^18^.

*X. fastidiosa* has been classified into subspecies ^19–21^, and genome analyses revealed widespread HR events within and among subspecies ^21–30^. Moreover, experimental evidence showed that *X. fastidiosa* strains are able to recombine in vitro during coculture of either live or live and dead strains ^31^. This process occurs under multiple experimental settings including batch and flow culture conditions with synthetic media or grapevine xylem sap from tolerant and susceptible varieties ^12,32,33^. Finally, many of the genes that are undergoing recombination among strains in vitro or in nature, are annotated as having roles in virulence and bacterial fitness^26^. These observations led to the hypothesis that HR, a driver of genetic diversity, is responsible for environmental adaptation ^26^ and/or host switching/expansion. In fact, intersubspecific HR has been suggested to have a role in plant host expansion by generating strains that infect mulberry ^20^, citrus, coffee ^22^, blueberry and blackberry ^29^.

The assembly, function, and regulation of TFP require multiple molecular components. Minor pilins, which are present in much lower quantities than the major pilin, prime the assembly of TFP ^34^, and some species-specific minor pilins may promote additional functions of TFP, such as aggregation and adherence ^35,36^. The functional diversity in pilins is mediated by differences in the C-terminal region of these proteins, since all pilins (including major and minor pilins) share a highly conserved N-terminal region ^36^. TFP is key to *X. fastidiosa* in both natural competence, which promotes HR and thus generates large genetic diversity ^26,27,31^, and twitching motility^18^, which is its only mean of movement and fundamental for a xylem-limited bacterium living under flow conditions. Therefore, in this study we individually assessed the functional role of a comprehensive set of 38 TFP-related genes by site-directed mutagenesis. We identified ten core genes that were essential for both natural competence and movement of this plant pathogen. However, some components, mostly minor pilins, were only essential to one feature or the other, and interestingly most paralogous genes had opposite functions, suggesting neofunctionalization. Additionally, our study identified the FimT3 minor pilin as the DNA receptor of the *X. fastidiosa* pilus. This minor pilin is found among plant pathogens mainly in bacterial species of the Xanthomonadaceae family. Together, our data provide insight into how TFP paralogs evolve to perform specific functions within bacterial cells, and how different bacterial species use unique minor pilins to acquire exogenous DNA through natural competence.

## Results

### Deletion of TFP-related genes differentially affects natural competence and movement of *X. fastidiosa*

Individual deletion mutant strains were generated by site-directed mutagenesis via natural transformation to determine the functional roles of 38 TFP-related genes (Table S1) ^37^. Genes were chosen according to genome annotation and searches based on homology to counterparts encoded by *Pseudomonas aeruginosa* ^38,39^. To the best of our knowledge, these genes represent the entire set of TFP genes encoded by *X. fastidiosa* strain TemeculaL. Assessment of natural competence was conducted using the pAX1-Cm plasmid, which recombines into the neutral site 1 of the *X. fastidiosa* genome and inserts a chloramphenicol (Cm) resistance cassette ^31,40^, whereas twitching motility was determined by measuring colony fringe widths on solid agar plates ^41^. Although the knockout of genes studied here did not always modulate natural competence and twitching motility in the same manner, the correlation between these two traits among mutant strains was positive (R^2^ = 0.56, *P*<0.001; Table S3), indicating that they were usually increased or reduced in a similar manner upon deletion of TFP genes.

A set of ten genes were considered core TFP components as deletion of such genes abrogated both natural competence and movement of *X. fastidiosa* (Fig. 1 and Fig. S1-3). They include the ATPases *pilB* and *pilT*, the prepilin peptidase *pilD*, the TFP assembly platform *pilC*, the whole alignment subcomplex *pilMNOP*, the secretin *pilQ* and the regulatory gene *pilZ*. Deletion of the sensor gene *pilS* of the PilRS two-component regulatory system (Fig. 1 and Fig. S1-3) significantly increased both natural competence and twitching motility, while deletion of the regulator *pilR* was essential for twitching motility but not for transformation. The minor pilin *fimT3* was the only gene found to be essential exclusively for natural competence (Fig. 1 and Fig. S1). Curiously, this *Δfimt3* strain showed significantly increased twitching motility than wild-type (WT) cells (Fig. 1 and Fig. S2-3).

**Fig. 1.**
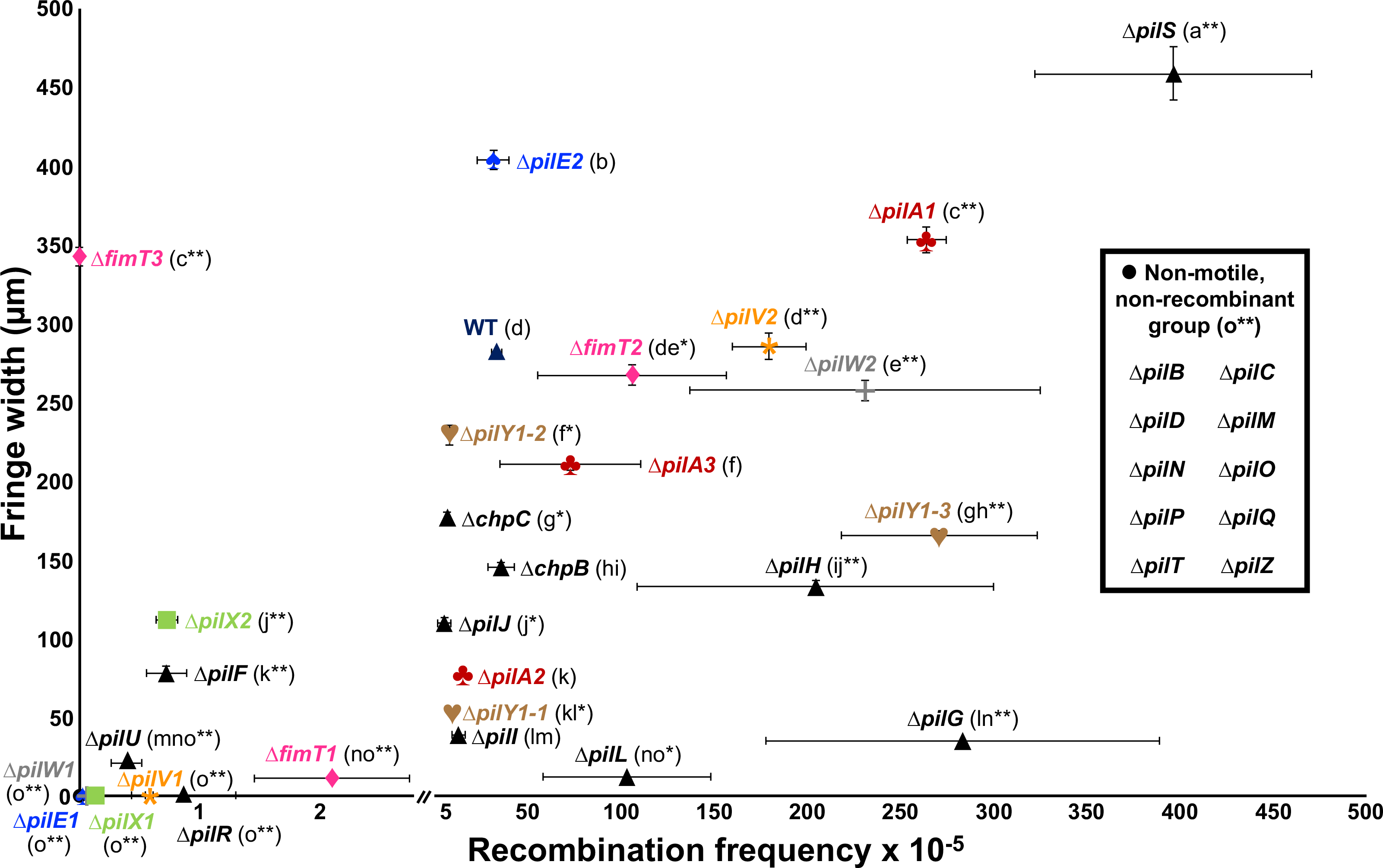
Natural transformation and twitching motility phenotypes of *X. fastidiosa* mutant strains used in this study. Quantification of natural transformation was performed by enumerating total viable cells and recombinants and results are expressed as the ratio of recipient cells transformed (recombination frequency; values shown in the x-axis of the chart). Twitching motility was determined by spotting cells on agar plates and measuring the movement fringe width after 4 days of growth at 28 °C (values shown in the y-axis of the chart). Results of mutant strains for paralogous genes are shown using the same color for text and symbol. The non-motile and non-recombinant mutants are shown as a black dot. The WT is represented as a dark blue triangle and all other mutants are represented as black triangles. Data represent means and standard errors. Different letters in parenthesis indicate significant difference in fringe width as analyzed by ANOVA followed by Tukey’s HSD multiple comparisons of means (*P*<0.05; n = three to 14 independent replicates with eight to 48 internal replicates each). * and ** indicate significant difference (*P*<0.05 and *P*<0.005, respectively) of recombination frequency through natural competence in comparison to the WT as determined using Student’s *t*-test (n = three to 21 independent replicates with two internal replicates each). The detection limit for recombination frequency was 10^−7^ and for twitching motility was 10 μm. Mutants below the detection limit are shown in the non-motile and non-recombinant group.

Seven sets of paralogs exist in the *X. fastidiosa* TFP cluster (PilA1-3, PilE/V/W/X1-2, PilY1-3 and FimT1-3). Intriguingly, the knockout of paralogs presented varied phenotypes suggesting duplication and neofunctionalization of the genes. The minor pilins *pilE1V1W1X1*, which are encoded in the same operon (Table S2), were essential for twitching motility but not for transformation (Fig. 1 and Fig. S2-3), although these mutants did show significantly lower recombination frequencies than WT (Fig. 1 and Fig. S1). In comparison to the WT, Δ*pilE2* presented significant higher twitching motility and no changes in natural transformation, Δ*pilV2* had significant higher natural transformation with no altered movement, Δ*pilW2* showed significant higher natural transformation and significant lower movement, and Δ*pilX2* possessed both significant lower natural transformation and twitching motility (Fig. 1 and Fig. S1-3). Knockout of *pilA1* significantly increased both natural competence and twitching motility, whereas deletion of *pilA2* and *pilA3* significantly reduced twitching motility and did not alter natural transformation (Fig. 1 and Fig. S1-3). A strain with deletion of both *pilA1* and *pilA2* abolished twitching motility (Fig. 1 and Fig. S2-3), as described elsewhere ^37^. Additionally, deletion of the minor pilin *fimT1* significantly decreased both natural competence and movement, while knockout of *fimT2* significantly increased natural competence and did not change twitching motility, and Δ*fimT3* lost natural competence and presented significantly higher movement than the WT (Fig. 1 and Fig. S1-3). As for *pilY1-1*, *pilY1-2* and *pilY1-3*, which are tip adhesins, their deletion significantly reduced twitching motility. However, Δ*pilY1-1* and Δ*pilY1-2* presented significant lower natural competence whilst Δ*pilY1-3* had significant higher natural competence than the WT (Fig. 1 and Fig. S1-3).

Lastly, deletion of the pilotin *pilF* and of the retraction ATPase *pilU* significantly reduced natural competence and movement. Knockout of the chemotaxis genes *pilGHIJL* and *chpBC* significantly reduced twitching. However, Δ*pilG*, Δ*pilH* and Δ*pilL* showed increased natural competence, while Δ*pilJ* and Δ*chpC* had lower natural competence, and Δ*pilI* and Δ*chpB* did not change this phenotype in comparison to the WT (Fig. 1 and Fig. S1-3).

#### TFP-related genes differentially modulate other virulence traits of *X. fastidiosa*

We evaluated phenotypes such as growth, biofilm formation, cell aggregation (measured by settling rate), and virulence in planta of the individual TFP mutants. Growth rates were measured by change in OD_600nm_ over time (Fig. S4-S5). Since this growth analysis considered only turbidity that may be affected by cell attachment (biofilm) to surfaces and cell-to-cell aggregation, the number of viable *X. fastidiosa* cells (measured as CFU/ml) was also evaluated. Only the non-recombinant and non-motile Δ*pilC* and Δ*pilP* mutants showed significant changes in the number of cells in comparison to WT, with both having significantly higher populations (Fig. S6). Biofilm formation is a critical aspect of virulence in *X. fastidiosa*. Individual deletion of *pilB*, *pilD*, *pilH*, *pilI*, *pilM*, *pilN*, *pilO*, *pilQ*, *pilR*, *pilT*, *pilX1* and *fimT1* significantly increased biofilm formation, whereas Δ*pilA3* and Δ*fimT3* formed significant less biofilm than WT (Fig. S7). Planktonic growth was significantly increased for strains with individual deletion of *pilA1*, *pilC*, *pilE1*, *pilF*, *pilM*, *pilN*, *pilO*, *pilP*, *pilQ*, *pilR*, *pilV1*, *pilW1*, *pilW2*, *pilX1*, *pilA1pilA2*, *pilY1-1*, *fimT3* and *chpC*, while individual knockout of *pilD*, *pilT* and *pilY1-3* significantly decreased planktonic growth (Fig. S8). Moreover, only Δ*pilD*, Δ*pilT* and Δ*pilX2* showed significant higher settling rates, with other mutants not differing significantly from the WT (Fig. S9). Biofilm formation and settling rate, as well as planktonic growth and growth rate were positively correlated (Table S3). Meanwhile, biofilm formation and twitching motility/growth rate, settling rate and planktonic growth/growth rate, and growth rate and total viable *X. fastidiosa* had significant negative correlations (Table S3). No other phenotype presented a significant correlation to recombination frequency besides twitching motility.

Virulence of a subset of the *X. fastidiosa* mutant strains was assessed by inoculating *Nicotiana tabacum* L. cv. Petite SR1 plants and scoring disease incidence and severity. Mutants chosen for analysis had different twitching motility phenotypes, such as higher movement (Δ*pilA1* and Δ*pilS*), lower movement (Δ*pilA2*) and non-motility (Δ*pilA1pilA2*, Δ*pilQ* and Δ*pilR*). Additionally, mutants in the DNA-binding *fimT3* (see below) and the paralogs *fimT1* and *fimT2* were tested and no significant change in virulence was found compared to WT (data not shown). WT-inoculated plants presented the highest disease severity over the time course of assays (Fig. S10a), with all plants inoculated with the mutant strains having significantly lower AUDPC (Area Under the Disease Progress Curve) values, except plants inoculated with Δ*pilA2* (Fig. S10b). All inoculated plants reached 100% disease incidence at the end of assays, except plants inoculated with Δ*pilA1*, which had 89% disease incidence (±11%, standard error). However, only WT-inoculated plants consistently presented severe symptoms of leaf scorch, while plants inoculated with the mutant strains mostly presented mild symptoms (Fig. S10d). In addition, when evaluating the population of *X. fastidiosa* in planta by qPCR, only the Δ*pilA1pilA2* double mutant showed a defect in colonizing the top leaf of the plant host, and the Δ*pilQ* and Δ*pilR* mutants presented significant higher population in the basal leaf of plant hosts (Fig. S10c). Together, results demonstrate that the deletion of analyzed genes significantly reduced symptoms development in infected tobacco plants (except for deletion of *pilA2*), which was not explained by bacterial distribution, since most strains could effectively colonize the entirety of plants (except Δ*pilA1pilA2*).

#### The minor pilin FimT3 is the DNA receptor of the *X. fastidiosa* type IV pilus

The observation of the minor pilin FimT3 being the only TFP component that is essential exclusively for natural transformation (Fig.1 and Fig. S1-3) led us to hypothesize that this component is the DNA receptor of the TFP of *X. fastidiosa*. We first analyzed the piliation phenotype of Δ*fimT3* in comparison to WT by transmission electron microscopy (TEM). Both presented long TFP as well as short type I pili, with no apparent changes in the piliation of *X. fastidiosa* cells upon deletion of *fimT3* (Fig. 2a). Next, the ability of these cells to take up DNA from the extracellular environment was evaluated by exposing them to Cy-3-labeled pAX1-Cm plasmid and observing under a fluorescent microscope. After exposure to the fluorescently labeled DNA fragment, WT cells imported DNA allowing for protection from exogenous DNaseI, but no fluorescent DNA foci were present in Δ*fimT3* cells (Fig. 2b). DNA foci were mostly observed at one pole of the cells, which coincides to the described phenotype of *X. fastidiosa* that forms TFP at only one of its poles ^42^. The DNA-uptake ability of the mutant strains of other minor pilins that decreased the natural transformation ability of *X. fastidiosa* upon deletion was also evaluated for comparison. These included Δ*pilE1*, Δ*pilV1*, Δ*pilW1*, Δ*pilX1*, Δ*pilX2*, Δ*fimT1* and Δ*fimT2* (Fig.1 and Fig. S1). The Δ*fimT2* mutant strain was included because this gene is a paralog of *fimT3*, even though its deletion increased the natural competence of *X. fastidiosa* (Fig.1 and Fig. S1). All analyzed mutant strains maintained the DNA-uptake (competence) phenotype (Fig. S11), but at different levels (Fig. S12). WT cells presented the highest percentage of cells with DNA foci, with Δ*pilE1* having the lowest proportion of cells with DNA acquisition (Fig. S12).

**Fig. 2.**
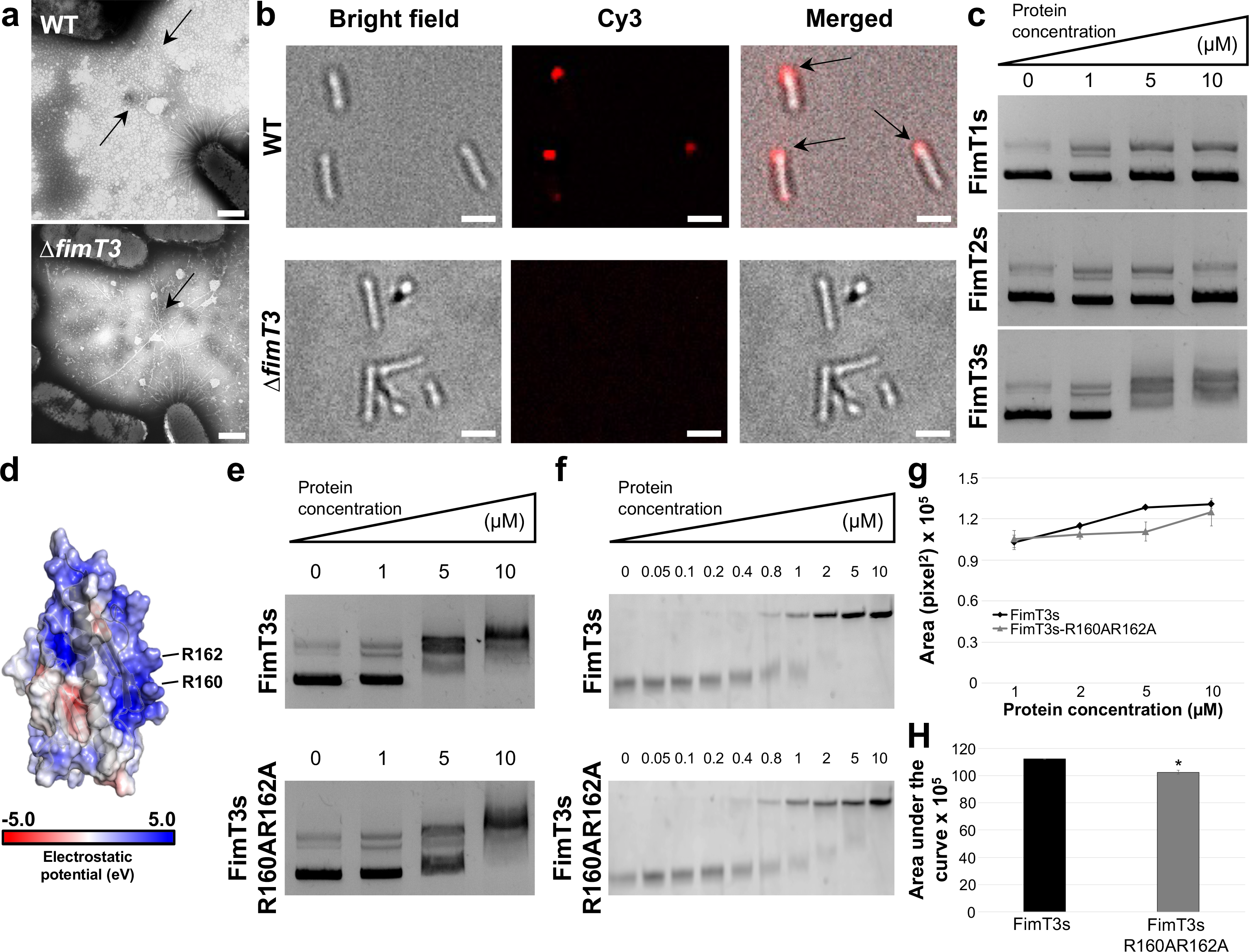
FimT3 DNA binding activity. **a** Transmission electron microscopy micrographs of pilus formation by *X. fastidiosa* WT and Δ*fimT3* cells. Arrows are pointing to TFP. Similar events were captured in two independent experiments. Images were captured at 31,500× magnification. Scale bars, 500 nm. **b** Uptake of Cy-3-labeled pAX1-Cm plasmid into a DNaseI resistant state by *X. fastidiosa* cells observed using a fluorescent microscope. The images correspond to the bright field (left), Cy-3 channel (center) and merged images (right). In merged images, arrows are pointing to fluorescent DNA foci at the cell poles. Similar events were captured in three independent experiments. Images were captured at 100× magnification. Scale bars, 1.5 μm. **c** DNA-binding ability of purified FimT1s, FimT2s and FimT3s assessed by agarose EMSA. Similar events were captured in three independent experiments. **d** Surface electrostatics representation of FimT3 merged with its ribbon representation. The arginine amino acid residues (R160 and R162) that were mutated for functional studies are highlighted in the figure. The electrostatic potential (eV) color scale is shown in the figure. **e** DNA-binding ability of purified FimT3s and FimT3s-R160AR162A assessed by agarose EMSA. Similar events were captured in three independent experiments. **f** Titration of the DNA binding activity of FimT3s and FimT3s-R160AR162A by native acrylamide EMSA. Similar events were captured in two independent experiments. **g** Densitometry analysis of the DNA binding activity of FimT3s and FimT3s-R160AR162A in acrylamide EMSA. The fluorescent intensity of the shifted DNA bands of Cy-3-labeled Km resistance cassette presented in **f** was measured using ImageJ. The fluorescent intensity of shifted bands when treated only with 1, 2, 5, and 10 μM of each protein was measured, since they presented higher shifts in electrophoretic mobility within this protein concentration range. **h** Area under the fluorescent curve. The area under the fluorescent curves in **g** was calculated to quantify the DNA-binding affinity of FimT3s and FimT3s-R160AR162A. Statistical significance was determined using Student’s *t*-test (* indicates *P*<0.05 in comparison to the wild-type protein; n = two independent replicates).

To determine the DNA-binding ability of FimT3, and compare to its paralogs FimT1 and FimT2, the soluble portion of these proteins (named FimT3s, FimT1s and FimT2s, respectively), which lacked the conserved N-terminal hydrophobic α-helices of pilins ^36^, were amplified and cloned for expression. These proteins were then purified and assayed for DNA binding by agarose electrophoretic mobility shift assay (EMSA). Results showed FimT3s was the only protein capable of binding DNA, since no shift was observed with FimT1s and FimT2s when using as much as 10 μM of each protein (Fig. 2c). Moreover, FimT3s was able to bind to different DNA sequences, independent of homology to the *X. fastidiosa* TemeculaL genome, indicating that binding is not sequence-specific (Fig. S13). Collectively, *fimT3* being the only gene essential exclusively for natural competence, the ability of FimT3 to bind to DNA and the loss of DNA acquisition from the extracellular environment by cells upon its deletion, demonstrate that this minor pilin is the DNA receptor of the *X. fastidiosa* TFP.

To further characterize FimT3, we searched for specific amino acid residues involved in DNA-binding. The binding of DNA to TFP via a minor pilin was demonstrated in ComP of *Neisseria*, ComZ of *Thermus thermophilus*, and VC0858 of *Vibrio cholerae* ^4,43,44^. Amino acids alignments revealed that FimT3 and ComP share only 28% identity (64% similarity) over 20% of query sequence, while FimT3 and ComZ have no significant identity. Alignment of FimT3 with VC0858 of *Vibrio cholerae* showed 35% identity and 62% similarity over 18% of query sequence. In spite of the low overall homology the sequence alignments showed that an arginine residue at position 162 of FimT3 aligned with a lysine of ComP (K108) that was previously demonstrated to be essential for its DNA-binding ability ^43^ (Fig. S14a). Additionally, an arginine residue at position 160 of FimT3 aligned with an arginine of VC0858 (R168) that was also important for the DNA-binding activity of the *V. cholerae* pilus ^4^ (Fig. S14b). These two arginine residues of FimT3 (R160 and R162) were predicted in silico to have the highest probabilities of binding DNA among the residues of this protein (Table S4). Moreover, alignment of FimT3 with a FimT ortholog that is essential for natural competence of *Acinetobacter baylyi ^45^* (28% identity and 46% similarity over 91% of query sequence) showed alignment of R162 of FimT3 to an arginine of FimT from *A. baylyi*. Likewise, FimT3 shares 27% identity (48% similarity) over 80% of query sequence with a recently described DNA-binding FimT ortholog of *Legionella pneumophila*, in which the aligned R160 and R162 have been demonstrated as important to bind DNA ^46^ (Fig. S14c).

Analysis of a structural model produced by the Phyre2 algorithm, showed that R160 and R162 of FimT3 are located within an electropositive surface and could form a pocket to bind DNA (Fig. 2d). Thus, to assess the role of these arginine residues in the DNA-binding ability of FimT3, we generated a FimT3s variant by site-directed mutagenesis in which both arginine residues were replaced by alanine residues (FimT3s-R160AR162A). Agarose EMSAs revealed that WT-FimT3s had a higher affinity for DNA than FimT3s-R160AR162A based on the amount of protein needed to shift 50% of the input DNA (Fig. 2e). Similar results were obtained when using the more sensitive acrylamide EMSA and the kanamycin (Km) resistance cassette sequence as target DNA (previously determined to be bound by FimT3s in Fig. S13). An abrupt shift was detected when using concentrations of FimT3s ranging between 1 and 5 μM, whereas a much more subtle and gradual shift was observed for samples treated with FimT3s-R160AR162A within the same concentration range (Fig. 2f). We then performed a densitometry analysis of the shifted bands treated with concentrations ranging between 1 and 10 μM of each protein (Fig. 2g) and calculated the area under the curve to quantify the DNA-binding affinity of FimT3s and FimT3s-R160AR162A. While the DNA-binding affinity of FimT3s-R160AR162A was lower than that of FimT3s (Fig. 2h), the mutations did not completely abolish DNA-binding ability suggesting other residues are contributing to this activity. Nonetheless, these results show that R160 and R162 are important for DNA-binding activity of FimT3.

#### FimT3 is widely conserved within *X. fastidiosa* strains and, among plant pathogens, is encoded by other members of the Xanthomonadaceae family

Previously, DNA-binding homologues of FimT3 have been found in many representative species of γ-Proteobacteria, including the Xanthomonadaceae family to which *X. fastidiosa* belongs ^46^. We performed searches using the Conditional Reciprocal Best BLAST (CRB-BLAST) ^47^, as well as phylogenetic analyses, to further explore the distribution of the *fimT* paralogs within this family. All three paralogs of *fimT* are encoded by *X. fastidiosa*, *X. taiwanensis*, many clade II *Xanthomonas* spp.^48^ and few *Pseudoxanthomonas* spp. (Fig. 3). Curiously, early branching clade I *Xanthomonas* spp. only encode *fimT1* and *fimT3* (Fig. 3). Specifically, *fimT3* was present in 1,416 out of 3,001 genomes of the Xanthomonadaceae (47%), majority of them belonging to species within the genus *Xanthomonas*. In the *fimT3* clade, *X. fastidiosa* strains grouped together, including *X. taiwanensis*, with a similar trend being observed for *Xanthomonas* spp. strains (Fig. 3). However, clade I and clade II *Xanthomonas* spp. and other Xanthomonadaceae species clustered mostly separated (Fig. 3). The alignment of the different FimT3 sequences encoded by *X. fastidiosa* strains showed a maximum of six nonconservative substitutions within strains belonging to the subspecies *pauca*, which encode FimT3 that are 13 amino acids shorter than other strains (Fig. S15a). Alignment of representative FimT3 sequences among strains belonging to the Xanthomonadaceae family revealed that the arginine residues at positions 160 and 162 of FimT3 from *X. fastidiosa* strain TemeculaL were conserved in most orthologous sequences of this protein (Fig. S15b). Additionally, R160 was always preceded by a glycine residue, constituting a highly conserved GRxR motif (Fig. S15b), as seen elsewhere ^46^. Although amino acid identities ranged from 95.60% to 100% within *X. fastidiosa*, and from 15.5% to 80% compared to other species, only 41 out of 1,416 FimT3 sequences encoded by the Xanthomonadaceae do not carry the GRxR motif (Table S5). Overall, results indicate that FimT3 is encoded by representative plant pathogens of the Xanthomonadaceae and that R160 and R162 are highly conserved within FimT3 sequences.

**Fig. 3.**
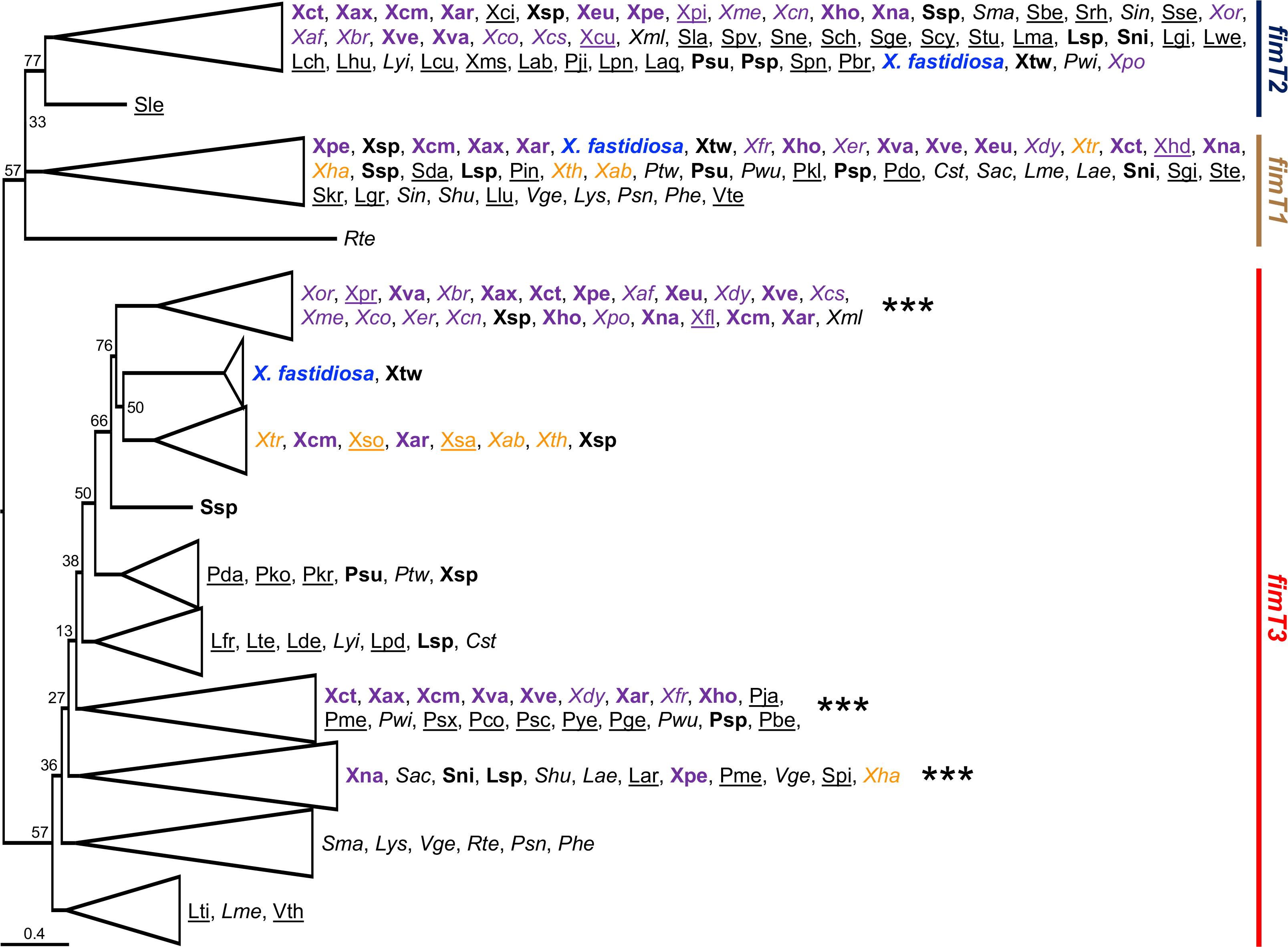
Phylogenetic tree based on nucleotide sequences of the three *fimT* paralogs from Xanthomonadaceae bacterial strains with whole-genome sequences available. The phylogenetic tree was built using the Maximum-likelihood method and visualized using FigTree. Branches with bootstrap values below 70% were collapsed, except where indicated. This was performed to keep conciseness of the figure while indicating the different clusters for *fimT1*, *fimT2* and *fimT3*. *X. fastidiosa* is highlighted in blue within the tree. When known, clade I *Xanthomonas* spp. are highlighted in orange, and clade II *Xanthomonas* spp. are highlighted in purple. Bacterial species encoding all three *fimT* paralogs are indicated in bold, while those encoding two paralogs are shown in italics and those encoding only one paralog are underlined within the tree. The order of appearance of bacterial species in collapsed branches follows the original order of appearance within the tree. For representation purposes, branches are highlighted by color according to each *fimT* paralog. Green: *fimT1*; Dark blue: *fimT2*; Red: *fimT3*. *** Indicates clusters within the *fimT3* clade which contain few FimT3 sequences with no GRxR motif (see Table S5 and Fig. S15). Abbreviations are the following. ***Coralloluteibacterium*:** Cst – *C. stylophorae*. ***Luteimonas*:** Lab – *L. abyssi*; Lae – *L. aestuarii*; Laq – *L. aquatica*; Lar – *L. arsenica*; Lch – *L. chenhongjianii*; Lcu – *L. cucumeris*; Lde – *L. deserti*; Lfr – *L. fraxinea*; Lgi – *L. gilva*; Lgr – *L. granuli*; Lhu – *L. huabeiensis*; Llu – *L. lumbrici*; Lma – *L. marina*; Lme – *L. mephitis*; Lpd – *L. padinae*; Lpn – *L. panaciterrae*; Lsp – *L*. sp.; Lte – *L. terrae*; Lti – *L. terricola*; Lwe – *L. wenzhouensis*; Lyi – *L. yindakuii*. ***Lysobacter*:** Lys *– L*. sp. ***Pseudoxanthomonas*:** Pbe – *P. beigongshangi*; Pbr – *P. broegbernensis*; Pco – *P. composti*; Pda – *P. daejeonensis*; Pdo – *P. dokdonensis*; Pge – *P. gei*; Phe – *P. helianthi*; Pin – *P. indica*; Pja – *P. japonensis*; Pji – *P. jiangjuensis*; Pkl – *P. kalamensis*; Pko – *P. kaohsiungensis*; Pkr – *P. koreensis*; Pme – *P. mexicana*; Psc – *P. sacheonensis*; Psn – *P. sangjuensis*; Psp – *P*. sp.; Psx – *P. spadix*; Psu – *P. suwonensis*; Ptw – *P. taiwanensis*; Pwi – *P. winnipegensis*; Pwu – *P. wuyuanensis*; Pye – *P. yeongjuensis*. *Rehaibacterium*: Rte – *R. terrae*. *Silanimonas*: Sle – *S. lenta*. *Stenotrophomonas*: Sac – *S. acidaminiphila*; Sbe – *S. bentonitica*; Sch – *S. chelatiphaga*; Scy – *S. cyclobalanopsidis*; Sda – *S. daejeonensis*; Sge – *S. geniculata*; Sgi – *S. ginsengisoli*; Shu – *S. humi*; Sin – *S. indicatrix*; Skr – *S. koreensis*; Sla – *S. lactitubi*; Sma – *S. maltophilia*; Sne – *S. nematodicola*; Sni – *S. nitritireducens*; Spn – *S. panacihumi*; Spv – *S. pavanii*; Spi – *S. pictorum*; Srh – *S. rhizophila*; Sse – *S. sepilia*; Ssp – *S*. sp; Ste – *S. terrae*; Stu – *S. tumulicola*. ***Vulcaniibacterium*:** Vge – *V. gelatinicum*; Vte – *V. tengchongense*; Vth – *V. thermophilum*. ***Xanthomonas*:** Xab – *X. albilineans*; Xaf – *X. alfalfae*; Xar – *X. arboricola*; Xax – *X. axonopodis*; Xbr – *X. bromi*; Xcm – *X. campestris*; Xcn – *X. cannabis*; Xcs – *X. cassavae*; Xci – *X. cissicola*; Xct – *X. citri*; Xco – *X. codiaei*; Xcu – *X. curcubitae*; Xdy – *X. dyei*; Xer – *X. euroxanthea*; Xeu – *X. euvesicatoria*; Xfl – *X. floridensis*; Xfr – *X. fragariae*; Xho – *X. hortorum*; Xha – *X. hyacinthi*; Xhd – *X. hydrangea*; Xml – *X. maliensis*; Xms – *X. massiliensis*; Xme – *X. melonis*; Xna – *X. nasturtii*; Xor – *X. oryzae*; Xpe – *X. perforans*; Xpi – *X. pisi*; Xpo – *X. populi*; Xpr – *X. prunicola*; Xsa – *X. sacchari*; Xso – *X. sontii*; Xsp – *X*. sp.; Xth – *X. theicola*; Xtr – *X. translucens*; Xva – *X. vasicola*; Xve – *X. vesicatoria*. ***Xylella*:** Xtw – *X. taiwanensis*.

## Discussion

In this study we defined the roles of individual subunits involved in regulation, assembly, and functioning of the *X. fastidiosa* TFP on natural competence, twitching motility, adhesion, and biofilm (Fig. 4). To our knowledge, our study comprises the largest simultaneous functional description of a comprehensive set of TFP-related genes. By performing site-directed mutagenesis of all TFP genes followed by functional studies, we defined a set of ten core genes (*pilBCDMNOPQTZ*) that were essential for natural competence and movement. Most of these proteins are inner- or outer-membrane associated, confirming the important role of membrane anchorage for functionality of TFP.

**Fig. 4.**
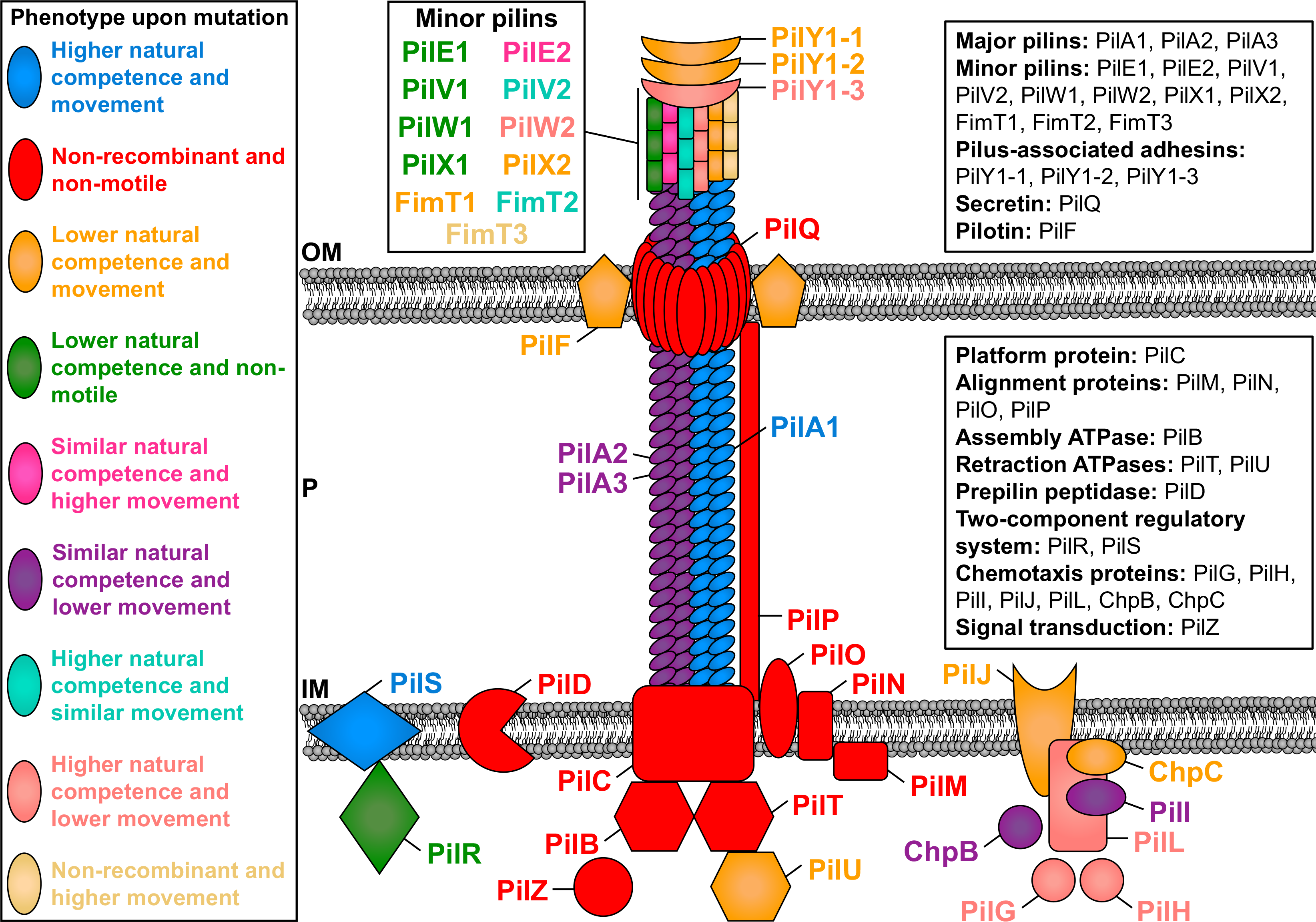
Schematic representation of the functional role of TFP molecular components in natural competence and twitching motility of *X. fastidiosa*. The figure shows the structure of a type IV pilus (PilA1, PilA2 and PilA3), the alignment subcomplex (PilM, PilN, PilO and PilP), the inner membrane motor subcomplex (PilB, PilC, PilD, PilT and PilU), the pore subcomplex (PilF and PilQ), minor pilins (PilE1, PilE2, PilV1, PilV2, PilW1, PilW2, PilX1, PilX2, FimT1, FimT2 and FimT3), TFP tip adhesins (PilY1-1, PilY1-2 and PilY1-3), as well as regulatory proteins (PilG, PilH, PilI, PilJ, PilL, PilR, PilS, PilZ, ChpB and ChpC). Proteins in the figure are color-coded according to their phenotype upon knockout mutation and thus functional role. More information about the role of each protein is described in Table S1, throughout the manuscript and in SI Discussion. OM – outer membrane; P – periplasm; IM – inner membrane.

Collectively, our data highlight the pervasive neofunctionalization among TFP components since seven paralogous gene sets exerted mainly opposing regulatory functions on movement and natural competence. For instance, paralogs of the minor pilins *pilEVWX* and *fimT* and the TFP tip adhesin *pilY1* are mostly organized as operons in two different regions of the *X. fastidiosa* TemeculaL genome (Table S2). Here, we observed that individual mutations in one set of these paralogs abrogated twitching motility and greatly reduced natural competence, while deletions in the other set of paralogous genes produced variable phenotypes that were mostly opposite to their counterparts. It has been demonstrated that minor pilins prime the assembly of TFP ^34^. This interaction is specific, in which major pilins require a specific cognate subset of minor pilins for TFP assembly ^49^. Therefore, the presence of paralogs for major and minor pilins displaying opposite functions within *X. fastidiosa* possibly suggest assembly of multiple TFP fibers with specific functions. However, this needs to be further assessed (see SI discussion). Most remarkably among paralogs, is the discovery of the minor pilin FimT3 as the TFP DNA receptor. This is an outstanding neofunctionalization example since FimT1 and FimT2 did not show DNA adherence abilities. This finding was also supported by our phylogenetic analysis, which showed divergent evolution of FimT3 from its paralogs. FimT3 plays an essential role in natural competence by mediating extracellular DNA uptake that may be integrated into the genome, thereby conferring new phenotypic traits that may allow an increase in bacterial fitness and/or host expansion of *X. fastidiosa*. The interaction between DNA molecules and an extracellular specific receptor is a key first step in the DNA uptake and thus natural competence of bacterial species. Initially, DNA binding by TFP has been observed in *P. aeruginosa* ^50^, *Streptococcus pneumoniae* ^51^, and *V. cholerae* ^4^, in which pilus retraction drives DNA uptake by transporting it to the cell surface, which is then imported to the periplasm by the competence protein ComEA ^4,7^. Although these early studies provide evidence that the specialized DNA receptor resides within type IV competence pilus fibers, no direct binding of DNA by individual purified major and/or minor pilins was reported. In *V. cholerae*, targeted mutations of positively charged amino acid residues within the minor pilins VC0858 and VC0859 reduced the DNA-binding affinity of its TFP ^4^, but the ability of these minor pilins to directly bind DNA was not assayed. Direct DNA binding has been first observed for the pilin protein ComP from *N. meningitidis*, with binding occurring at a positively charged surface of this protein ^43^. ComP orthologs are found mostly within the Neisseriaceae family, where they mediate DNA uptake by binding to specific DNA uptake sequences, as *Neisseria* spp. preferentially take up homotypic DNA ^43,52^. ComZ, encoded by *T. thermophilus*, was the second minor pilin described as a TFP DNA receptor ^44^. However, ComZ is an unusual minor pilin that possesses an additional large β-solenoid domain inserted into the common β-sheet structure of pilins. Furthermore, ComZ binds to another minor pilin, PilA2 (PilV in *X. fastidiosa*), and its DNA binding ability does not appear to require positive charges in its surface ^44^. Nonetheless, the binding of ComZ to another minor pilin suggests that specific interactions within the pilus structure may occur to assemble the competence pilus fiber.

Here, the deletion of individual paralogs of the major pilin PilA abrogated neither natural competence nor twitching motility, suggesting functional overlapping among them (see SI Discussion). In addition to the novel functions described above, our study confirmed the role of core components that were previously analyzed in other bacterial species including *Acinetobacter baumannii* ^53^, *A. baylyi* ^45^, *T. thermophilus* ^54^, *Haemophilus influenzae* ^55^, *P. aeruginosa* ^56–59^, *L. pneumophila^46^* and *X. fastidiosa* ^32,42,60^. The exceptions were *pilP* and *pilZ* in which previous deletion of both greatly reduced natural competence ^61,62^, but did not abolish it here. It is of note that very few of these genes have been simultaneously assayed for their role in both natural transformation and movement mediated by TFP.

Deletion of FimT3 in *X. fastidiosa*, but not of the paralogs FimT1 and FimT2, exclusively disrupted the natural competence of this bacterium. FimT3 binds to DNA via a surface in which positively charged arginine residues contribute to the DNA binding ability of this protein. However, the binding appears to be non-specific, which is in contrast to ComP ^43^. FimT from the human pathogen *L. pneumophila* ^46^, which is a homologue of the FimT3 studied here, was recently identified as a pilin with DNA-binding ability. Another FimT homolog was also demonstrated to be specialized in natural competence in *A. baylyi* ^45^, although no DNA binding was assessed. In our study, FimT3 seems to be particularly enriched in plant pathogens of the Xanthomonadaceae family, being encoded by all *X. fastidiosa* strains as well as many other bacterial species belonging to this family. Thus, our results, together with previous studies, demonstrate that DNA binding by TFP occurs among bacterial species through a similar component such as minor pilins, which themselves are not widely distributed or conserved, but are rather specific to different bacterial groups. For instance, the ability of strain-specific subsets of minor pilins to modulate TFP dynamics has already been described in *P. aeruginosa* (see SI Discussion) ^49^. Understanding how this minor pilin interacts with different minor and major pilins of *X. fastidiosa* in the competence pilus is an important area to explore. It is not known whether the other Xanthomonadaceae bacterial species encoding FimT3 that were found in our analysis are also naturally competent. For instance, *X. fastidiosa* strains belonging to the *pauca* subspecies have a shorter FimT3 amino acid sequence, missing part of the C-terminal, and have a few aminoacidic changes (although maintain the arginine residues studied here). The only reported attempt to transform a *X. fastidiosa* strain belonging to the subspecies *pauca* by natural competence in vitro has failed ^63^. Studies with the *X. fastidiosa* subsp. *pauca* FimT3 variant should investigate both individual DNA-binding activity and potential roles of the C-terminus in interaction with the rest of the TFP structure.

In summary, our study highlights the fundamental functions of TFP in virulence and evolution evidenced by complex regulatory features. Due to the worldwide threat that *X. fastidiosa* presents to many economically important crops and to the ever-increasing list of plant hosts, it is imperative to investigate how this plant pathogen acquires foreign DNA and expands its genetic diversity to help researchers understand its eco-evolutionary history and re-emergence of this pathogen in many geographic areas. In addition, our results provide a robust blueprint to assess TFP functions in bacteria living under flow conditions. An extensive discussion including additional genes analyzed in this study and their functional roles in natural transformation and twitching motility of *X. fastidiosa*, in addition to unique behaviors and other phenotypes such as biofilm formation, virulence in planta, and diversity in paralogous genes functions, is included in SI Discussion. All of this indicates that a complex and tightly regulated machinery is involved in these bacterial functions.

## Methods

Methods that are only cited here are described in detail in SI Materials and Methods.

### Bacterial strains, plasmids, and culture conditions

All strains and plasmids used in this study are listed in Table S6. All *X. fastidiosa* mutant strains used here are derivatives of the *X. fastidiosa* subsp. *fastidiosa* strain TemeculaL ^26^, and were originated by site-directed mutagenesis of each gene of interest using natural competence as described elsewhere ^37^. Primers used in this process are listed in Table S7. *X. fastidiosa* was recovered from −80 °C glycerol stocks and routinely cultured for seven days at 28 °C on periwinkle wilt (PW) agar plates ^64^, modified by removing phenol red and using bovine serum albumin (1.8 g/l) (BSA; Gibco Life Sciences Technology), and sub-cultured onto fresh PW agar plates for another seven days at 28 °C before use. *X. fastidiosa* mutant strains were grown similarly but using PW plates amended with Km. All assays were performed using the subcultured *X. fastidiosa* strains. Cells were suspended and cultured in PD3 broth ^64^ to perform phenotypic assays, while phosphate-buffered saline (PBS) was used to suspend cells in liquid for in planta assays. Whenever needed, the antibiotics Km and Cm were used at concentrations of 50 and 10 μg/ml, respectively. When used together with Cm, the Km concentration was reduced to 30 μg/ml. Luria-Bertani (LB) medium (BD Difco) was used to culture *E. coli* cells. When needed, Km, Cm and ampicillin (Amp) were added to LB at concentrations of 50, 35 and 100 μg/ml, respectively.

### Phenotypic analyses of *X. fastidiosa* WT and mutant strains

Analyses of natural competence among strains ^31^, twitching motility ^33^, growth curve and growth rate ^31^, biofilm formation and planktonic growth ^31,65^, settling rate ^31^, piliation observation by TEM ^37^, and virulence in planta using tobacco as model system ^66^ were performed as described elsewhere. All experiments had at least three biological replicates, unless otherwise stated.

### DNA uptake assays

DNA uptake assays were performed as similarly described for the analysis of natural competence among *X. fastidiosa* strains (see SI Materials and Methods). Briefly, recipient cells were suspended in PD3 broth to OD_600nm_ = 0.6, spotted onto PD3 agar plates and grown for three days at 28 °C. Then, 1 μg of fluorescently labeled pAX1-Cm plasmid (10-μl volume), labeled using the *Label* IT Nucleic Acid Labeling kit, Cy3 (Mirus Bio LLC), was added on top of cells, air-dried, and incubated at 28 °C for another 24 h. After, cells were harvested in 150 μl of PD3 broth, and a 50 μl aliquot was treated with 10 units of DNase I (New England Biolabs) for 10 minutes at 37 °C to degrade the remaining extracellular DNA, and cells were observed using a 100× oil immersion objective in a Nikon Eclipse Ti inverted microscope (Nikon). Image acquisition was performed using a Nikon DS-Q1 digital camera (Nikon) controlled by the NIS-Elements software version 3.0 (Nikon), which was also used to create merged fluorescent images. To detect Cy3, an excitation wavelength of 590 nm was used (tetramethyl rhodamine isothiocyanate; TRITC filter). The proportion of cells that acquired extracellular DNA was calculated as the percentage of cells with fluorescent DNA foci to cells without fluorescent DNA foci. Experiments were performed at least three times independently.

### Protein cloning, expression, and purification

Cloning, expression, and purification of FimT1s, FimT2s and FimT3s were performed similarly as described elsewhere using the pHIS-Parallel1 plasmid (Table S6) ^67,68^. Amino acid exchanges in FimT3s were performed by designing a pair of primers (Table S7) that exchanged the arginine residues at positions 160 and 162 by alanine residues. This pair of primers was designed using the QuickChange Primer Design tool (Agilent Technologies, Inc.; https://www.agilent.com/store/primerDesignProgram.jsp). For amino acid exchanges, the whole pHIS-Parallel1-*fimT3s* construct was amplified via *Pfu* DNA polymerase (G-Biosciences) using a standard protocol from the manufacturer and this pair of primers in a S1000 thermal cycler (Bio-Rad). Expression and purification of FimT3s-R160AR162A were performed as described above (see SI Materials and Methods) ^67^. PCR products were purified using the Gel/PCR DNA Fragments Extraction kit (IBI Scientific). Correct cloning and amino acid exchanges in all constructs were confirmed by Sanger sequencing (Sequetech Corporation) using the T7 promoter forward primer (Table S7).

### DNA binding assays

Agarose EMSAs were mainly performed to assess the DNA-binding ability of purified FimT1s, FimT2s and FimT3s. Briefly, 200 ng of DNA (usually pAX1-Cm plasmid) were incubated for 30 minutes at 28 °C with increasing concentrations of purified proteins in 20 μL EMSA reaction buffer (50 mM Tris-HCl, pH 7.5; 50 mM NaCl; 200 mM KCl; 5 mM MgCl_2_; 5 mM EDTA, pH 8.0; 5 mM DTT; 0.25 mg/ml BSA). Lysis buffer containing 250 μM imidazole, in which proteins were suspended, was included as blank control. After the incubation period, DNA was separated by electrophoresis on a 0.8% agarose gel containing GelRed nucleic acid gel stain (Biotium) in Tris-acetate-EDTA buffer (120 V for 40 minutes). Experiments were performed at least three times independently. On the other hand, native acrylamide EMSAs were used to perform titration of the DNA binding activity of FimT3s and FimT3s-R160AR162A. In summary, 60 ng of Cy-3 labeled Km resistance cassette were incubated for 30 minutes at 28 °C with increasing concentrations of purified proteins in 20 μL EMSA reaction buffer and separated by electrophoresis on a 3.5% native acrylamide gel in Tris-borate-EDTA buffer (40 V for 4 hours). Native acrylamide gels were pre-run at 40 V for 30 minutes before being used. DNA samples were directly visualized using the ImageQuant LAS 4010 Imaging System (GE Healthcare), since the used DNA was fluorescently labeled with Cy3. Experiments were performed two times independently. Densitometry analysis of the shifted bands of Cy-3 labeled DNA in native acrylamide gels was performed by measuring the fluorescent intensity of these bands when treated with 1, 2, 5, and 10 μM of each protein using ImageJ ^69^. Then, these values were used to calculate the area under the fluorescent curves, as performed to obtain the AUDPC (see SI Materials and Methods) ^70^, to quantify the DNA-binding affinity of FimT3s and FimT3s-R160AR162A by determining the progress of shifted bands in relation to each protein concentration.

### Bioinformatic analyses

FimT3 modeling was performed using the Phyre2 web portal ^71^ and visualized via PyMOL version 2.4.0 (Schrödinger, LLC). The surface electrostatics of FimT3 was predicted using the APBS Electrostatics plugin from PyMOL. Sequence alignments of FimT3 with ComP, VC0858 and FimT from *A. baylyi* and *L. pneumophila* were performed by retrieving the respective sequences from NCBI, aligning through the T-Coffee Multiple Sequence Alignment Server ^72^ and visualizing using the BoxShade webserver (https://embnet.vital-it.ch/software/BOX_form.html). For FimT3 analyses within the Xanthomonadaceae family, DNA sequences of different FimT homologs from *Xylella*, *Xanthomonas*, *Lysobacter* and *Stenotrophomonas* were used as query for screening genomes of Xanthomonadaceae using CRB-BLAST ^47^ with default settings. Nucleotide sequences for the CRB-BLAST hits were then retrieved and aligned using MAFFT ^73^, followed by RAxML version 8.0.24 ^74^ to build a phylogenetic tree, which was visualized using FigTree version 1.4.4 (http://tree.bio.ed.ac.uk/). FimT3-encoding sequences were retrieved from the phylogenetic tree using the TREE2FASTA Perl script ^75^, translated into amino acid sequences and aligned using Clustal Omega (Clustal 12.1) to determine percentage of identical amino acids ^76^. The visualization of the alignment of representative FimT3 sequences among *X. fastidiosa* strains and representative bacterial species belonging to the Xanthomonadaceae family was performed using the BoxShade webserver. The FimT3 GRxR motif sequence logo was generated using the WebLogo webserver^77^.

### Data analysis

Data from natural competence assays (recombination frequency) and the area under the fluorescent curve were compared by two-tailed Student’s *t*-test. Data from twitching motility, growth rate, viable *X. fastidiosa* CFU/ml obtained during natural competence assays, biofilm formation, planktonic growth, settling rate, AUDPC, *X. fastidiosa* population in planta and percentage of cells acquiring DNA from the extracellular environment were individually analyzed by one-way analysis of variance (ANOVA) followed by Tukey’s HSD multiple comparisons of means in R 4.0.0 under the package multcomp ^78^. Correlation among analyzed phenotypes was determined by Pearson’s correlation using the SigmaPlot software version 11.0 (Systat Software Inc.).

## Acknowledgments

We thank Dr. Michelle Mendonça Pena for helping with phylogenetic analyses. We thank the Alabama Supercomputing Authority for granting access to their high-performance computing platform. We acknowledge funding from Alabama Agricultural Experiment Station (AAES) Hatch Program (L.D., P.A.C., N.P) and Auburn Internal Grant Program.

## Author contributions

M.V.M., P.A.C. and L.D.F. designed research; M.V.M., X.Z. D.S., L.M.G., E.N. and N.P. performed research; M.V.M., N.P., P.A.C. and L.D. analyzed data; and M.V.M. and L.D.F. wrote the paper.

