## Supplemental figures and tables for "Complete functional analysis of type IV pilus components of a reemergent plant pathogen reveals neofunctionalization of paralog genes"

**Supplementary Information Text**

**SI Discussion**

Our core genes that were essential for both natural competence and twitching motility of *X. fastidiosa* represent a central machinery required for the biogenesis and functioning of type IV pili (TFP) resembling that of *P. aeruginosa*, a well-studied model for TFP and twitching motility [1]. This central machinery involves, in addition to the major pilin PilA, the retraction and extension ATPases PilB and PilT, respectively, the platform protein PilC, the prepilin peptidase PilD and the secretin PilQ [2]. Moreover, the essential function of PilMNOP in *X. fastidiosa* was close to that found in *N. meningitidis*, in which the presence of this alignment subcomplex is essential for the piliation phenotype and functions performed by TFP [3,4], with the opposite being described in *P. aeruginosa* [5]. Surprisingly, *pilZ* was essential not only for twitching motility, but also natural competence in our study. This seems to be the first observation of *pilZ* being essential for this phenotype. Many PilZ orthologs have been demonstrated to interact with the secondary messenger 3',5'-cyclic diguanylic acid (c-di-GMP) by directly binding this molecule [6-8] or by interacting with other c-di-GMP receptor protein [9]. Although the *X. fastidiosa* PilZ does not appear to possess canonical c-di-GMP binding domains, it could interact with this messenger through degenerate domains, as already describe to one of its proteins [10], or by interacting with an unidentified receptor. This would contribute to establish the apparent link between high levels of c-di-GMP and induction of twitching motility in this bacterium [11]. However, this hypothesis needs further elucidation.

In contrast, although required for movement, TFP genes are not essential for natural competence in *P. aeruginosa* [12]. Likewise, deletion of TFP genes in *V. cholerae* does not abrogate its natural competence [13]. This may be related to the lifestyle of each individual species, their specific needs and genome content. *X. fastidiosa* lives exclusively in the xylem of plant hosts and foregut of insect vectors [14] in addition to having a reduced genome in comparison to the closely related *Xanthomonas* spp. [15]. Besides, *X. fastidiosa* moves only through twitching motility [14]. Together, these findings suggest that this bacterium relies heavily on TFP genes to perform essential functions. Curiously, however, is the presence of many paralogs of TFP genes encoded by *X. fastidiosa*. Deletion of the major pilin *pilA* has been shown to abolish natural competence and movement in *Acinetobacter baylyi* [16] and *A. baumannii* [17], and only twitching motility in *P. aeruginosa* [18]. In our study, the individual deletion of *pilA* paralogs did not abrogate either phenotype, suggesting functional overlapping. *V. cholerae*, for example, encodes three different TFP, including the competence pilus [13], toxin-coregulated pili (TCP) [19], and mannose-sensitive hemagglutination (MSHA) pili [20]. Although TCP and MSHA pili are encoded by their own set of genes, rather than being encoded by *pil* and associated genes, the existence of different TFP led us to hypothesize that PilA1, PilA2 and PilA3 could compose different pilus with specific functions. We were particularly interested in defining whether these different paralogs could specialize in natural competence or twitching motility.

As mentioned above, our results instead suggested functional overlapping among PilA paralogs. Previously, TFP hyperpiliation was observed in a *X. fastidiosa* ∆*pilA1* mutant [21], which could explain our observation that deletion of this gene increases both natural competence and twitching motility of this bacterium. In addition, the same study described that deletion of *pilA2* leads to loss of movement [21], opposing our results. However, movement analysis of ∆*pilA2* was previously performed using PD3 agar plates, while in this study we used PW without BSA agar plates, which proved to be more conducive for twitching motility and thus yield more accurate results. Therefore, we determined that *pilA2* is not essential for the movement of *X. fastidiosa*, which was previously described [21]. Our ∆*pilA3* mutant produced similar phenotypic results as ∆*pilA2*, further supporting the overlapping of functions. However, we cannot discard that these *pilA* paralogs may play different roles according to different host conditions and/or environmental signals. For instance, different *pilA* alleles among strains of other bacterial species have been implicated in kin recognition [22] and susceptibility to a bacteriophage [23]. Nonetheless, a study performed with a non-pathogenic strain of *X. fastidiosa* encoding five *pilA* paralogs demonstrated that deletion of the global $\sigma^{54}$ regulator RpoN downregulated the expression of only one of these paralogs [24], suggesting some level of specificity.

In *P. aeruginosa*, the PilR regulatory protein of the PilRS two-component regulatory system is the transcription factor that binds to a *cis*-acting region upstream to the *pilA* promoter and enables its transcription by the RNA polymerase containing the alternative σ factor RpoN [25,26]. On the other hand, the PilS sensor protein interacts directly with PilA for pilin autoregulation [27]. When high levels of PilA are present in the inner membrane, PilS deactivates PilR by adopting a phosphatase-active conformation, whereas absence of PilA leads to phosphorylation of PilR by PilS that remains in a kinase state, which in turn increases transcription of *pilA* and of other coregulated genes presumably important for virulence and surface-associated behaviors of *P. aeruginosa* [27,28]. Deletion of *pilR* abrogates *pilA* transcription [29], with individual deletion of both *pilR* and *pilS* leading to significant reductions of twitching motility (or even abolition) in *P. aeruginosa* [28], *A. baumannii* [17], *A. baylyi* [16], *Dichelobacter nodosus* [30], *Xanthomonas oryzae* pv. *oryzae* [31], besides *X. fastidiosa* [32]. The twitching motility of the latter three was only evaluated in a Δ*pilR* background. Moreover, natural competence has been negatively impacted by the deletion of this two-component system in *A. baumannii* [17], but not in *A. baylyi* [16]. Our phenotypic results upon deletion of *pilR* follow what has been mostly described, with abrogation of movement and significant reduction in natural competence. Conversely, deletion of *pilS* in *X. fastidiosa* significantly increased both phenotypes, generating the highest recombination frequencies and longest fringe widths of twitching motility recorded in this study. This is the first report of a Δ*pilS* mutant having increased TFP-related phenotypes and suggests novel regulatory mechanisms in *X. fastidiosa*. Notably, the well-studied PilS of *P. aeruginosa* has six transmembrane segments and thus faces the cytoplasm of cells whilst interacting with PilA stored in their inner membrane [27,33]. The PilS of *X. fastidiosa*, however, has only five transmembrane domains (predicted using the TMHMM Server, v. 2.0) [34], which likely allows it to still interact with PilA in the inner membrane, but also suggests that it can interact with different or even additional signals for pilin regulation. This is supported by the fact that the vast majority of histidine kinase sensor proteins from two-component systems are membrane-bound with a C-terminal cytoplasmic kinase domain [35], and not periplasmic as predicted for *X. fastidiosa*. However, this hypothesis needs to be further explored.

The motility of bacterial species can also be controlled by chemotaxis, a mechanism in which cells sense environmental stimuli and move either toward or away from the chemical signal. In *X. fastidiosa*, twitching motility is regulated by the Pil-Chp operon composed of *pilGIJL-chpBC* [36]. In addition, we found that this plant pathogen encodes the PilH chemosensory response regulator, previously thought to be absent in this bacterium [36], in another area of the genome far from the Pil-Chp operon (Table S2). Similarly, the Pil-Chp chemosensory system that regulates TFP assembly and movement in *P. aeruginosa* is composed of *pilGHIJK-chpABC* [37-40]. *X. fastidiosa* lacks a *pilK* homolog, while PilL is likely this bacterium's counterpart to ChpA from *P. aeruginosa* [36]. Based on the *P. aeruginosa* and *Escherichia* *coli* (Gram-negative model system for chemotaxis involving flagellum) chemosensory systems, these are composed mainly by a transmembrane chemoreceptor (MCP/PilJ; located in the inner membrane) that interacts with histidine kinases (CheA/PilL) via adaptor proteins (CheW/PilI-ChpC) [38,41]. In *E. coli*, MCPs alter the kinase activity of CheA in the presence of ligand binding (chemical stimuli), which transfers a high-energy phosphoryl group to the response regulator CheY (PilG) that will then interact with the flagellar motor proteins to modulate the direction of rotation [41]. The adaptation to ligand concentrations is performed through reversible methylation changes to the MCP adjusted by a methyltransferase (CheR) and a methylesterase (CheB/ChpB) [41]. Likewise, the Pil-Chp system in *P. aeruginosa* controls TFP biogenesis by modulating the intracellular levels of the secondary messenger 3',5'-cyclic monophosphate (cAMP), whereas the response regulators PilG and PilH presumably control twitching motility in a cAMP-independent manner by modulating pilus extension and retraction mediated by the ATPases PilB and PilT, respectively, in a hitherto unidentified mechanism [42,43]. Individual deletions of *pilGIJ* in *P. aeruginosa* leads to impaired twitching motility [38,39,43], whereas Δ*pilH* mutants either displayed reduced movement or an altered pattern of twitching motility depending on the strain background [38,43]. Additionally, Δ*chpA* (PilL) mutants present significantly reduced twitching motility [37,43], with individual deletions of *chpBC* not altering this phenotype [37]. In *A. baylyi*, *pilG* knockout significantly reduced twitching motility with no apparent effects on natural competence [16]. On the contrary, deletion of *pilG* and *pilH* in *A. baumannii* significantly reduced twitching motility, but with deletion of *pilG* leading to abrogation of natural competence and deletion of *pilH* not affecting this phenotype [17]. Lastly, deletion of both *pilG* and *pilL* in *X. fastidiosa* leads to a deficient phenotype of twitching motility [36,44].

Our results showed that deletion of any gene belonging to the Pil-Chp chemosensory system of *X. fastidiosa* strain TemeculaL significantly reduced its twitching motility, although no abrogation of this phenotype was observed among deletion mutants. The observed differences with previous studies of *X. fastidiosa* may be due to different strain genetic backgrounds, even though the genes analyzed in this study from strain TemeculaL share 100% homology with corresponding genes in strain Temecula1, and/or gene knockout methods. Here, we have deleted genes of interest by entirely replacing them by a kanamycin-resistance cassette in *X. fastidiosa* strain TemeculaL, while in the other mentioned studies, knockout of *pilG* and *pilL* was performed by transposon insertion in strain Temecula1. Regarding natural competence phenotypes, we obtained variable results upon deletion of Pil-Chp genes. Regardless of reduced twitching motility, Δ*pilG*, Δ*pilH* and Δ*pilL* mutants presented significant higher natural competence phenotypes (contrarily to what was observed in *A. baylyi* and *A. baumannii*, described above), while deletion of *pilJ* and *chpC* reduced this trait and Δ*pilI* and Δ*chpB* did not present altered natural competence. Conclusively, we have completely assayed the Pil-Chp operon function on both natural competence and twitching motility of *X. fastidiosa* and determined that the individual components of this system were not essential for either of these traits in our study. Together, our findings with the PilRS two-component regulatory system and with the Pil-Chp chemosensory system suggest that TFP regulation in *X. fastidiosa* has its own specific characteristics, which may include alternate signaling pathways and/or environmental stimuli.

Returning to the analysis of paralogous genes in *X. fastidiosa*, next we have the minor pilins *pilEVWX* and *fimT*, as well as the TFP tip adhesin *pilY1*. The minor pilins *pilE1V1W1X1* and *fimT1*, as well as the tip adhesin *pilY1-1* are encoded in the same operon (Table S2). A similar genomic arrangement has also been observed in *P. aeruginosa* [45]. On the other hand, paralogs of these genes are encoded in two sequential operons located in a different region of the *X. fastidiosa* genome (Table S2). One operon is composed of *pilE2* and *pilY1-2* and the other of *pilV2W2X2* and *fimT2* (Table S2). Additionally, a third tip adhesin, *pilY1-3*, is present but is not encoded in an operon, while we determined that FimT3, which is also not encoded in an operon, is the DNA receptor of the *X. fastidiosa* TFP (discussed in the main manuscript). In *P. aeruginosa*, genetic analyses revealed the presence of the minor pilins *pilEVWX* and *fimTU* that are required for TFP assembly (except FimT, which is a substitute of FimU), twitching motility and infection by pilus-specific phage [2,45-48]. Moreover, it has been demonstrated that these minor pilins prime the assembly of TFP whilst promoting the display of the PilY1 adhesin at the cell surface and thus at the pilus tip [49]. Briefly, PilVWX interact with PilY1 and form a complex that is bound by PilE, whereas FimU (substituted by FimT in *X. fastidiosa*) contacts PilA directly and promotes its connection to PilVWXY1E by interacting with PilV and PilE. Minor pilins are essential for TFP assembly since their deletion abrogates pilus assembly and twitching motility [45-50]. However, *P. aeruginosa* mutant cells lacking the minor pilin operon still produce reduced amounts of TFP that are twitching motility-defective, which is mediated by the expression of minor pseudopilins from this bacterium's type II secretion system (TIISS) [49]. The TFP assembly system and the TIISS are evolutionary related and share many homologs [51,52]. Ultimately, in a strain background lacking minor pseudopilins encoded by the TIISS, the minimal components required for TFP assembly included PilVWXY1 and either FimU or PilE [49]. In *A. baylyi*, deletion of *pilW* abrogated both natural competence and twitching motility, while individual deletions of *pilV*, *pilY1* and *fimT* abolished natural competence and reduced twitching motility [16].

In *X. fastidiosa*, we observed that individual deletion of the genes belonging to the *pilE1V1W1X1*, *fimT1* and *pilY1-1* operon mostly abrogated twitching motility. The exceptions were Δ*fimT1* and Δ*pilY1-1*, which still presented movement, but significantly reduced. However, deletion mutants of all these genes were still naturally competent but displayed significantly reduced recombination frequencies. In *X. fastidiosa* strain Temecula1, deletion of *fimT1* abrogated twitching motility [32], contrary to our results. This may be due to different strain backgrounds and/or gene knockout methods, as commented above. Our results demonstrated that *pilE1V1W1X1* are essential for twitching motility of *X. fastidiosa* strain TemeculaL, while no gene encoded within this operon is essential for natural competence. Curiously, the presence of mutants defective in movement indicates that paralogs of these minor pilins and tip adhesin did not compensate for the loss of their counterparts. In fact, deletion mutants within the operons *pilE2Y1-2* and *pilV2W2X2*/*fimT2* produced variable results. Deletion of *pilE2* significantly increased movement and did not alter natural competence; Δ*pilV2* and Δ*fimT2* presented significant increased natural competence and no change in twitching motility; Δ*pilW2* displayed significant higher natural competence and significant lower movement; and deletion of *pilX2* and *pilY1-2* significantly reduced both phenotypes. On the other hand, the Δ*pilY1-3* mutant presented higher natural competence and lower twitching motility. Thus, no gene belonging to this operon was essential for either natural competence or twitching motility, suggesting a key role for the *pilE1V1W1X1*, *fimT1* and *pilY1-1* operon in comparison to their paralogs. Intriguingly, it has been demonstrated that *P. aeruginosa* strains encode major pilins associated with a specific set of minor pilins. This means that major pilins require their specific subset of minor pilins for both TFP assembly and twitching motility, since strains expressing the major pilin from another strain that has heterologous minor pilins do not assemble TFP nor perform twitching motility [53]. If this correlation also happens in *X. fastidiosa*, it may explain the lack of compensation by their paralogs when the *pilE1V1W1X1*, *fimT1* and *pilY1-1* operon was deleted. Moreover, the ability of minor pseudopilins from the TIISS in *P. aeruginosa* to assemble small amounts of TFP when minor pilins are deleted is suggested to occur at a lower affinity, leading to inefficient pilus extension that cannot compete with retraction [49]. This is because of the specific interaction of major pilins to their minor pilin subsets described above [53]. Nevertheless, we suggest that the ability of the minor pilin mutants Δ*pilE1V1W1X1* and Δ*fimT1* of *X. fastidiosa* to remain naturally competent at low frequencies regardless of losing their motility may be due to TFP assemble by TIISS minor pseudopilins encoded by this bacterium [54]. This would allow *X. fastidiosa* to assemble TFP that can still perform DNA uptake but are not able to carry out twitching motility. However, all our hypotheses raised here need to be further investigated.

It would be interesting to determine whether the different PilA paralogs encoded by *X. fastidiosa* require specific subsets of minor pilins. Strikingly, the strain we have analyzed in this study encodes three *pilA* paralogs, but only two subsets of minor pilins. Additionally, this strain encodes a third *pilY1* paralog not encoded within the two subsets of minor pilins. PilY1 was initially described in *Neisseria* spp. (in which it is named PilC) as important for adherence to epithelial cells and thus acting as a pilus tip adhesin [55]. In *P. aeruginosa*, PilY1 was shown to bind calcium ($Ca^{2+}$) and oppose pilus retraction in a calcium-dependent manner [56]. In *X. fastidiosa*, the paralog PilY1-2 is the only TFP tip adhesin that has a Ca-binding motif and likely contributes for this bacterium to cope within the xylem sap of plant hosts, which is an environment with high Ca concentrations [57]. Deletion of *pilY1* in *X. fastidiosa*, however, has generated antagonistic results. Deletion of *pilY1-1* in some studies has been shown to significantly reduce twitching motility [32,58]. Conversely, analysis of the same mutant strain and of Δ*pilY1-2* in another study demonstrated that deletion of these genes does not alter the movement phenotype of *X. fastidiosa* [57]. As mentioned above, our study, however, demonstrated that deletion of all three *pilY1* paralogs significantly reduced twitching motility. Nonetheless, phenotypic results of natural competence and twitching motility from our study and others demonstrate that the presence of different *pilY1* paralogs with variable domains in *X. fastidiosa* may dictate function specificity.

At last, deletion of both *pilF* and *pilU* in *X. fastidiosa* led to significantly reduced natural competence and twitching motility phenotypes. PilF has been described in *P. aeruginosa* as a pilotin (outer membrane lipoprotein) required for the localization and assembly of the multimeric PilQ secretin, being essential for both TFP and twitching motility of this bacterium [59,60]. In *A. baylyi*, both *pilF* and *pilU* are essential for natural competence and twitching motility. On the other hand, PilU is a secondary retraction ATPase homologous to PilT essential for twitching motility of *P. aeruginosa* [61]. Our ∆*pilU* results, however, are close to those described for *A. baumannii* and *V. cholerae* in which PilU is not essential for natural competence [13,17]. Thus, PilU does not work as an independent retraction ATPase, but it likely functions in conjunction with PilT to increase the retraction force of TFP [17,62,63]. Curiously, this mechanism was also suggested in *P. aeruginosa* despite previous phenotypic results upon deletion of *pilU* [64].

TFP have also been demonstrated to affect other bacterial phenotypes besides natural competence and twitching motility. These include aggregation and adherence, biofilm formation, conjugation, electron transfer, manipulation of host cells, virulence, and secretion of proteins [65]. Therefore, we analyzed whether deletion of TFP-related genes could also affect additional key phenotypes of *X. fastidiosa*, including growth, biofilm formation, cell aggregation and virulence in planta. Deletion of genes analyzed in this study greatly changed the growth curve and growth rate of *X. fastidiosa*. However, since these analyses are traditionally performed by only measuring the turbidity of bacterial cultures, we also investigated the ability of TFP genes to alter the growth of this bacterium by counting the number of viable cells (CFU/ml) grown during natural competence assays. This was performed because *X. fastidiosa* presents higher recombination frequency via natural competence when growing exponentially [66], thus changes in growth could indirectly reflect in its natural competence. However, only two non-recombinant mutant strains, ∆*pilC* and ∆*pilP*, had altered growth by presenting significant higher number of viable cells than the WT, thus indicating that our natural competence results were not affected by changes in the growth of *X. fastidiosa*.

Previously, deletion of *pilG* and *pilL* has been shown to significantly decrease biofilm formation by *X. fastidiosa* [36,67], while knockout of *pilB*, *pilQ* and *fimT1* significantly increases this phenotype [32,68], and deletion of *pilY1-1* and *pilY1-2* does not affect it [57]. Our results mostly followed what has been described for *X. fastidiosa*, except for ∆*pilG* and ∆*pilL*, which presented similar biofilm formation to the WT. This may be due to different strain genetic backgrounds and/or knockout methods as discussed above. Nonetheless, we have established here that 14 TFP-related genes modulate biofilm formation, whereas 19 TFP genes modulate planktonic growth, an opposite phenotype to biofilm formation. In addition, only three genes have been found to modulate cell aggregation despite its known positive correlation to biofilm formation [14]. We have indeed found a positive correlation between biofilm formation and settling rate (measurement of cell aggregation), while these two phenotypes were negatively correlated with planktonic growth (Table S3). Therefore, our results demonstrate that TFP may modulate other important phenotypes of *X. fastidiosa* besides natural competence and twitching motility. However, it is not known whether TFP genes directly regulate biofilm formation, planktonic growth and cell aggregation in *X. fastidiosa* by hitherto unknown mechanisms or if that is an indirect reflect of altered twitching motility phenotypes, which in turn modulate biofilm formation [14].

The functional role of some TFP genes in the virulence in planta of *X. fastidiosa* has also been previously investigated. Deletion of *pilG* and *pilL*, which abrogated twitching motility in previous studies [36,44], resulted in avirulence and delayed, less severe symptoms in infected grapevines, respectively [36,67]. Moreover, the non-motile Δ*pilB* and Δ*pilQ* mutants have significantly reduced basipetal translocation in planta, which occurs in a reverse direction away from the leaves and thus against the flow of xylem sap [68]. Similarly, Δ*pilA*, Δ*pilQ* and Δ*pilT* mutants in *R. solanacearum*, which are all impaired in twitching motility, also presented reduced virulence in tomato plants [69,70]. However, all these studies were performed with twitching motility-deficient mutant strains. Here, we have assayed the virulence of *X. fastidiosa* using mutant strains with variable phenotypes, including higher movement, lower movement, and non-motility. Regardless, all presented reduced virulence in planta, except for the Δ*pilA2* mutant, which promoted similar disease severity to WT-inoculated plants. Although the results obtained for movement-deficient strains were in accordance with their twitching motility phenotypes, we expected that mutant strains with higher twitching motility would be hypervirulent. Likewise, *X. fastidiosa* mutant strains lacking the *rpfF* gene, which encodes the diffusible signal factor (DSF; quorum sensing molecule of this bacterium) [71] synthase, have increased motility and are hypervirulent in planta [57,71-73]. DSF negatively regulates twitching motility in *X. fastidiosa* and positively modulates biofilm formation [71]. Thus, the higher virulence of Δ*rpfF* is linked to increased movement of *X. fastidiosa* within infected plants and higher colonization of xylem vessels [72,73]. In our study, however, we have mutated genes directly linked to TFP assembly and functioning, and not genes upstream of these processes. Also, we did not observe significant impairment in the colonization of infected plants by the analyzed mutant strains. This may be due to experimental bias, since we have inoculated basal leaves of young tobacco plants, which possibly allowed *X. fastidiosa* cells to colonize the plants by moving with the flow of xylem sap. Therefore, it would be interesting to complement our study in the future by inoculating the canopy of full-grown plants and observing whether colonization will be impaired or enhanced due to the observed changes in twitching motility in vitro upon deletion of TFP-related genes, which may be reflected in their basipetal movement. Together, the fact that twitching motility-deficient strains, as well as mutant strains with higher movement, had reduced virulence in this study indicate that TFP genes are required for the proper functioning and regulation of this machinery, which is needed for full symptom development; whilst deletion of its individual components suppresses virulence.

Overall, our data demonstrate that *X. fastidiosa* has a central TFP machinery that is composed and functions similarly to other well-studied TFP systems, such as the ones from *P. aeruginosa* and *N. meningitidis*. However, *X. fastidiosa* also has many unique behaviors involving mainly its regulatory proteins and minor pilins, which are worth being investigated to better understand the TFP dynamics of this bacterium.

**SI Methods**

**Site-directed mutagenesis of genes of interest in *X. fastidiosa* strain TemeculaL.** The deletion of each gene of interest (GOI) (Table S1) in *X. fastidiosa* strain TemeculaL [74] was performed using a protocol developed by our research group [21]. Briefly, the upstream and downstream regions immediately flanking each GOI were amplified from the *X. fastidiosa* TemeculaL genome using pairs of primers (Table S7) containing overlapping nucleotides with the Km resistance cassette encoded by the pUC4K plasmid (Table S6). When deleting genes in operons that share overlapping nucleotides with flanking genes, the corresponding nucleotides were maintained in the primer design to mitigate frameshift mutations. To obtain the targeting construct, the amplified upstream and downstream regions of each GOI were fused to the Km resistance cassette via overlap-extension PCR, as detailed elsewhere [21]. The purified PCR product was then used to transform WT *X. fastidiosa* strain TemeculaL cells through natural competence directly. In summary, WT cells were suspended in PD3 broth to $OD_{600nm}$ of 0.25 (~$10^8$ cells/ml), and 10 µL of this suspension was spotted on a PD3 agar plate, with 10 µL of the targeting construct being spotted on top of it. The resulting mix of cells and the targeting construct was air-dried and incubated at 28 ºC for five days. Then, cells were suspended into 1ml of PD3 broth and plated into PW agar plates amended with Km for the selection of mutant strains obtained through homologous recombination. Deletion of each GOI was confirmed through PCR (Table S7), in which non-amplification of an internal sequence of each GOI, and amplification of an internal sequence of the upstream region of each GOI and the Km resistance cassette, confirmed deletion of each of these genes in *X. fastidiosa* strain TemeculaL (WT was included as control in each PCR reaction). The obtained mutant strains were stored as 25% glycerol stocks in PD3 broth at -80 ºC until use. PCR reactions were carried out using a standard protocol with the iProof High-Fidelity PCR kit (Bio-Rad) in a S1000 thermal cycler (Bio-Rad). PCR products and agarose gel fragments were purified using the Gel/PCR DNA Fragments Extraction kit (IBI Scientific). The pUC4K plasmid was prepared from an overnight culture of *E. coli* Dh5$\alpha$ using the extraction kit GeneJET Plasmid Miniprep kit (Thermo Scientific).

**Analysis of natural competence among *X. fastidiosa* strains.** The plasmid pAX1-Cm [75], prepared from an overnight culture of *E. coli* EAM1 (Table S6) [76] using the extraction kit GeneJET Plasmid Miniprep kit (Thermo Scientific), was used in this assay. The plasmid concentration was adjusted to 100 ng/µl before using (Cytation 3 Image Reader spectrophotometer; BioTek Instruments Inc.) and aliquots were stored at -20 ºC until use. For natural competence assays, recipient cells were suspended in PD3 broth to $OD_{600nm}$ of 0.25 and spotted onto PD3 agar plates similarly as described above for site-directed mutagenesis. Briefly, 10 µl of cells were spotted together with 1 µg of pAX1-Cm (10-µl volume), air-dried, and incubated at 28 ºC for five days. Then, cells were suspended in 1 ml of PD3 broth, diluted by 10-fold serial dilutions and plated in selective (PW+Cm agar plates for WT cells and PW+Km+Cm agar plates for mutant strains) and non-selective (PW agar plates for WT cells and PW+Km agar plates for mutant strains) media for growth of recombinants and total viable cells (both counted as CFU/ml), respectively. After 21 days of incubation at 28 ºC, CFUs were enumerated for recombinants and total viable cells and the recombination frequency was calculated as the ratio of the number of recombinants to total viable cells. At least three independent biological replicates with two internal replicates were performed for each *X. fastidiosa* strain. To confirm homologous recombination, five recombinant CFUs from each strain per biological replicate were randomly selected and plated onto new selective medium agar plates to observe growth, and colony PCR was performed using a pair of primers designed to amplify the Cm resistance cassette [21] (Table S7). PCR reactions were carried out using a standard protocol with the *Taq* 5X Master Mix (New England Biolabs) in a S1000 thermal cycler (Bio-Rad).

**Twitching motility assays.** The twitching motility of *X. fastidiosa* strains was evaluated using PW agar plates without BSA, as previously described [77]. In short, 15 to 20 spots of each strain (six strains per plate) were made onto PW without BSA plates using the needle side of a sterile inoculation loop (Globe Scientific). Plates were incubated at 28 ºC for 4 days before analysis. Then, the colony peripheral fringes were visualized under ×10 magnification using a Nikon Eclipse Ti inverted microscope (Nikon). Image acquisition was performed using a Nikon DS-Q1 digital camera (Nikon) controlled by the NIS-Elements software version 3.0 (Nikon). Fringe widths were measured for six colonies per strain per plate, with four measurements per colony, using the ImageJ software [78]. Twitching experiments were performed at least three times independently, with 48 internal replicates each.

**Growth curve and growth rate, biofilm formation and planktonic growth, and settling rate measurements of *X. fastidiosa* strains.** *X. fastidiosa* cells were suspended in PD3 broth to $OD_{600nm}$ of 0.25 and inoculated into polystyrene 96-well plates (Corning Inc.) to generate growth curves [79]. Eight wells were used per strain, which were inoculated as 10 µl aliquots into 190 µl of PD3 broth. In addition, eight wells per plate were inoculated with 200 µl of PD3 broth to serve as controls. Plates were incubated at 28 ºC and 150 rpm for eight days and the $OD_{600nm}$ value for each well was measured daily using the Cytation 3 Image Reader spectrophotometer (BioTek Instruments Inc.). $OD_{600nm}$ values were adjusted by subtracting values from control wells. Growth rates were calculated as the slope of the line obtained by performing natural log-transformation of the growth values at the exponential growth phase (two to six days post inoculation) using the formula: rate = [ln ($OD_{600nm}$ day 6) - ln ($OD_{600nm}$ day 2)]/time (days) [79]. On the other hand, planktonic growth was evaluated by transferring a 150 µl aliquot of the supernatant of each suspension at the final day of evaluation (day 8) to a new 96-well plate and measuring the $OD_{600nm}$, while biofilm formation was quantified by staining cells that remained attached to each well using a 0.1%

crystal violet solution, as previously described [80]. Briefly, wells were gently rinsed three times with Milli-Q water and stained with 230 µl of 0.1% crystal violet solution for 20 minutes at room temperature. Then, the crystal violet solution was removed, the wells were gently rinsed once again three times with Milli-Q water, and the crystal violet was solubilized by adding 230 µl of 95% ethanol and incubating under agitation (150 rpm) for 5 minutes. The $OD_{600nm}$ values of wells were then measured using the Cytation 3 Image Reader spectrophotometer (BioTek Instruments Inc.). At last, settling rate (used as a measure of cell aggregation), was determined by suspending cells in PD3 broth to $OD_{600nm}$ of 1.0 and measuring the $OD_{600nm}$ values in a cuvette when cells settled exponentially (0 to 2 hours post inoculation; hpi) [79]. As used for growth rate, the settling rate was calculated using the formula: rate = [ln ($OD_{600nm}$ 0 hpi) - ln ($OD_{600nm}$ 2 hpi)]/time (hours). Experiments were performed at least three times independently.

**Transmission electron microscopy.** The piliation phenotype of cells was visualized under a transmission electron microscope, as described elsewhere [21]. Briefly, two-day old *X. fastidiosa* cultures were harvested from PW without BSA agar plates and suspended in 200 µl of sterile Milli-Q water. Then, 10 µl of each cell suspension were pipetted onto a Formvar-coated TEM grid (Electron Microscopy Sciences) and cells were allowed to settle for 10 minutes. After, the leftover liquid was blotted out using a filter paper and the grid was negatively stained with 10 µl of 2% phosphotungstic acid (PTA) for 10 seconds. The excess PTA was then removed using filter paper, and grids were air-dried and observed under a Zeiss EM10 transmission electron microscope (Carl Zeiss), with images being captured at 31,500× magnification using the MaxIm DL software (Diffraction LTD). Alternatively, grids were placed directly on top of *X. fastidiosa* cells growing in PW without BSA agar plates for few seconds and negatively stained with PTA, as described above, for observation in the transmission electron microscope. Two independent experiments were performed.

***X. fastidiosa* in planta virulence assays using the model plant *Nicotiana tabacum*.** For in planta assays, *Nicotiana tabacum* L. cv. Petite Havana SR1 plants were propagated at the Plant Science Research Center at Auburn University (AL, USA) and inoculated with *X. fastidiosa* TemeculaL WT, Δ*pilA1*, Δ*pilA2*, Δ*pilA1pilA2*, Δ*pilQ*, Δ*pilR* and Δ*pilS* strains using a modified protocol from elsewhere [81].

The experiment was conducted in a completely randomized design. Briefly, transplanted three-week-old tobacco plants were inoculated by pin pricking using 23-gauge needles with 20 μL of either PBS (mock control) or individual suspensions of each *X. fastidiosa* strain at an $OD_{600nm}$ of 1.0 in PBS. Inoculation at the base of the second and third (counting from the bottom) leaf petioles was done twice with one week apart. In total, plants were individually inoculated with PBS (n=10) and *X. fastidiosa* strains (n=10). Disease incidence (percentage of symptomatic plants per total inoculated plants) and severity were recorded weekly for nine-time points after first symptom appearance (about 8 weeks post-inoculation). Disease severity was calculated for each plant by counting symptomatic leaves and total number of leaves [(symptomatic leaves/total leaves) × 100], whereas the Area Under the Disease Progress Curve (AUDPC) was calculated by the midpoint rule method [82] using disease severity data. Two independent experiments were performed.

On the other hand, the *X. fastidiosa* population in planta was determined by qPCR using leaf samples from the end time point of disease evaluation. To analyze the ability of *X. fastidiosa* strains to colonize infected plants, basal and top leaves were used in this analysis. DNA was extracted from 100 mg of the petiole of each leaf using a modified CTAB protocol [83]. qPCR reactions were performed using the HL5/HL6/HLp pair of primers and TaqMan probe labeled with FAM (Table S7) [84]. DNA amplifications were performed using a standard protocol with the PerfeCTa Multiplex qPCR ToughMix Low ROX kit (Quantabio) in a C1000 thermal cycler base with a CFX96 real-time system (Bio-Rad), using standard cycling parameters. DNA amplification of each sample was repeated in triplicates. Reaction efficiencies of 95 to 105% were confirmed for each qPCR run. The number of *X. fastidiosa* cells in each sample was calculated by comparing the obtained values with a standard curve made from 10-fold serial dilutions of the genomic DNA of *X. fastidiosa*. Two independent experiments were performed.

**Cloning, expression, and purification of FimT1s, FimT2s and FimT3s.** FimT1s, FimT2s and FimT3s from *X. fastidiosa* strain TemeculaL were cloned into pHIS-Parallel1 for *E. coli* expression (primers used in this process are listed in Table S7). PCR reactions were carried out using a standard protocol with the iProof High-Fidelity PCR kit (Bio-Rad) in a S1000 thermal cycler (Bio-Rad). PCR products and agarose gel fragments were purified using the Gel/PCR DNA Fragments Extraction kit (IBI

Scientific). Restriction enzymes and T4 DNA ligase used in the cloning process were obtained from New England Biolabs and Promega, respectively. *E. coli* BL21(DE3) cells carrying each vector were grown in LB broth to $OD_{600nm}$ 0.6-0.8, and protein expression was induced with 400 μM of isopropyl β-D-1-thiogalactopyranoside (*IPTG*) for 3 hours and 30 minutes. Then, cells were collected by centrifugation (4,000 rpm for 15 minutes at 4 ºC), suspended in lysis buffer (500 mM NaCl, 20 mM Tris, pH 8.0) and disrupted by sonication. The soluble fractions were collected by centrifugation (12,000 rpm for 30 minutes at 4 ºC) and the proteins were individually purified by affinity chromatography using Ni-NTA agarose resin columns (Thermo Scientific). Protein concentration in each sample was determined by Bradford assay (VWR) and, when needed, samples were concentrated using the Spin-X UF Concentrator (Corning) and centrifugation (12,000 rpm for 1 hour at 4 ºC). Protein purifications were visualized by submitting samples to 10% SDS-PAGE (150 V for 1 hour) and staining with Coomassie Brilliant Blue.

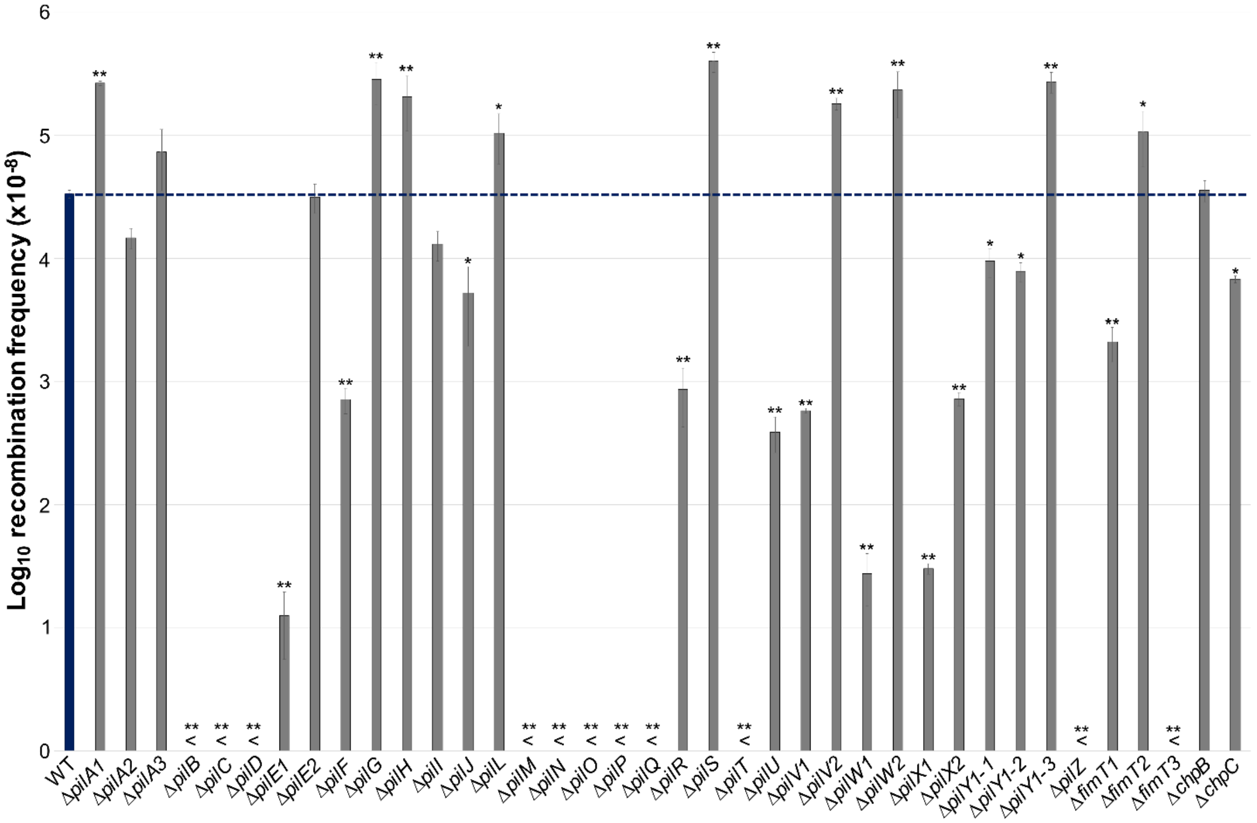


**Fig. S1.** Recombination frequencies obtained through natural transformation of *X. fastidiosa* mutant strains used in this study. Quantification of natural transformation was performed in PD3 plates by applying 1 µg of the pAX1-Cm plasmid to equivalent numbers of recipient cells of each *X. fastidiosa* strain to generate chloramphenicol-resistant ($Cm^R$) mutations. Total viable cells and transformants were counted and results are expressed as the ratio of recipient cells transformed. WT is highlighted in blue, and the dashed blue line indicates the mean value of recombination frequency for the WT. Data represent means and standard errors. * and ** indicate significant difference ($P$<0.05 and $P$<0.005, respectively) of recombination frequency through natural competence in comparison to the WT as determined using Student's *t*-test (n = three to 21 independent replicates with two internal replicates each). "<" indicates below detection limit, which was $10^{-7}$. Mutant strains below the detection limit were considered non-recombinant.

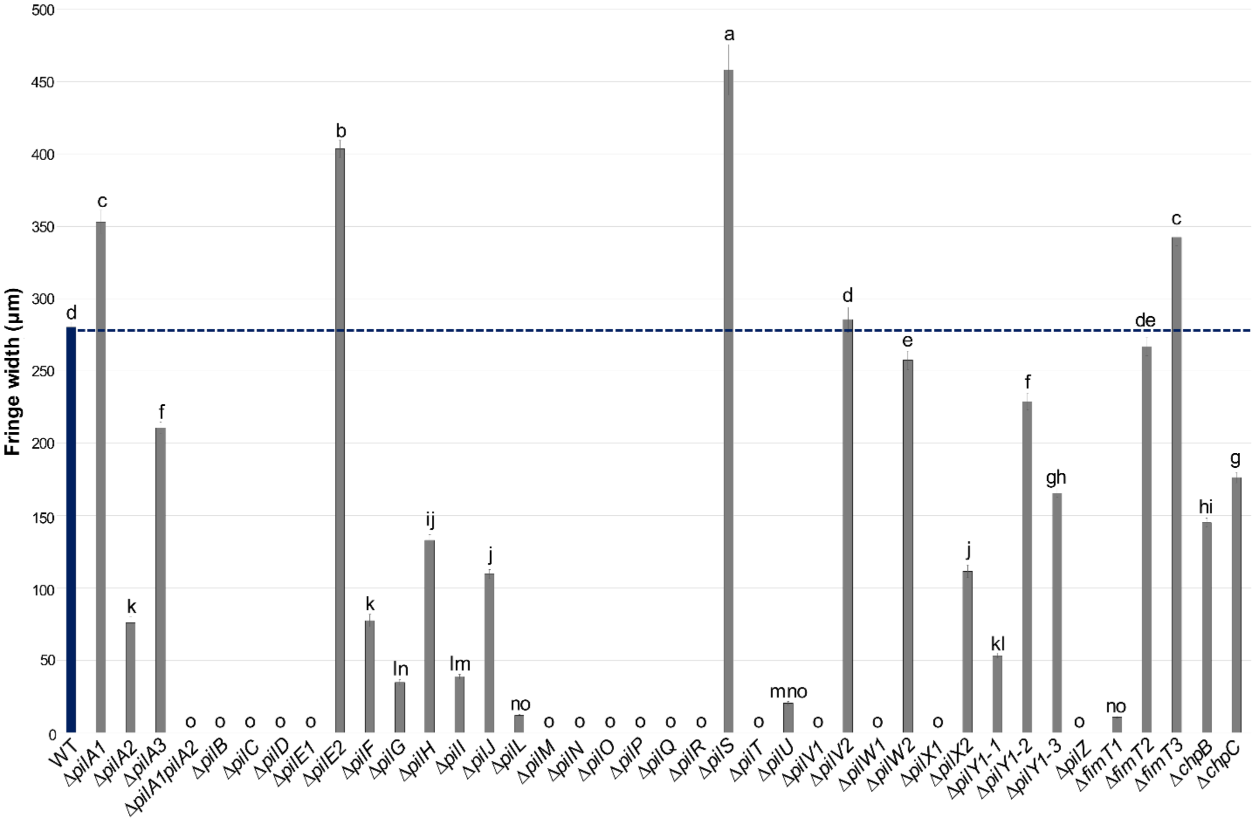


**Fig. S2.** Fringe width measurements of the twitching motility phenotypes of *X. fastidiosa* mutant strains used in this study. Twitching motility was determined by spotting cells of each strain in PW without BSA plates and measuring the movement fringe width after 4 days of growth at 28 ºC. WT is highlighted in blue, and the dashed blue line indicates the mean value of fringe width for the WT. Data represent means and standard errors. Different letters on top of bars indicate significant difference as analyzed by ANOVA followed by Tukey's HSD multiple comparisons of means ($P$<0.05; n = three to 14 independent replicates with eight to 48 internal replicates each). The detection limit was 10 µm. Mutant strains below the detection limit were considered non-motile.

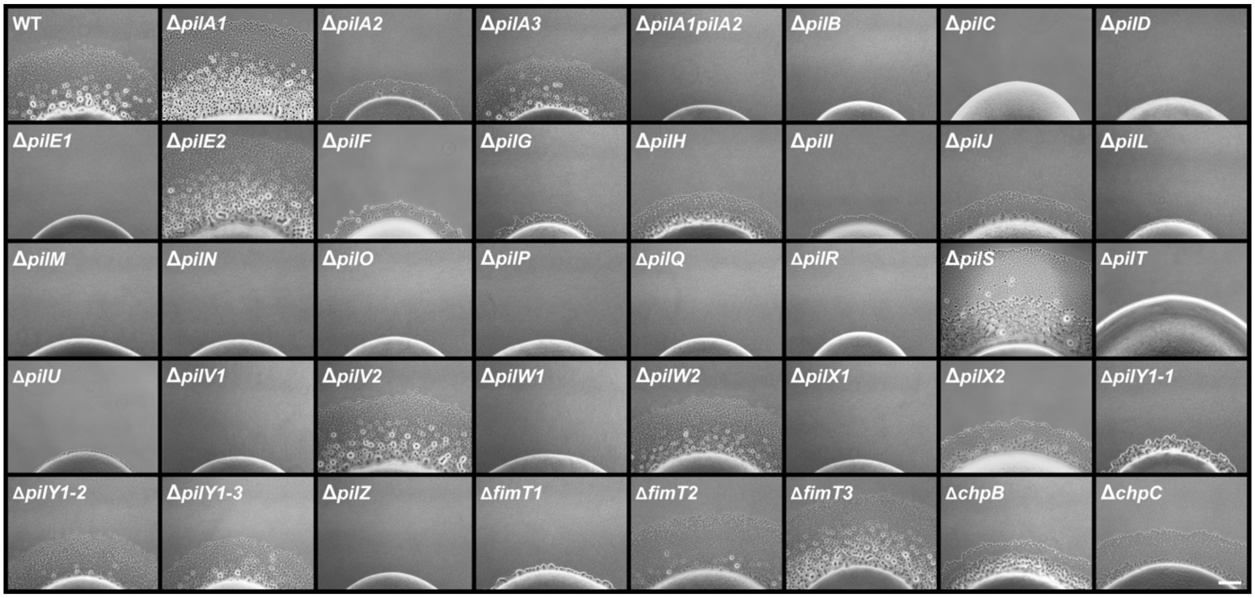


**Fig. S3.** Figure panel with representative pictures of the twitching motility phenotypes of *X. fastidiosa* mutant strains used in this study. The assay was performed as described in figure S2. Similar events were captured in three to 14 independent experiments. Images were captured at 10× magnification. Scale bar (right lower panel), 100 μm.

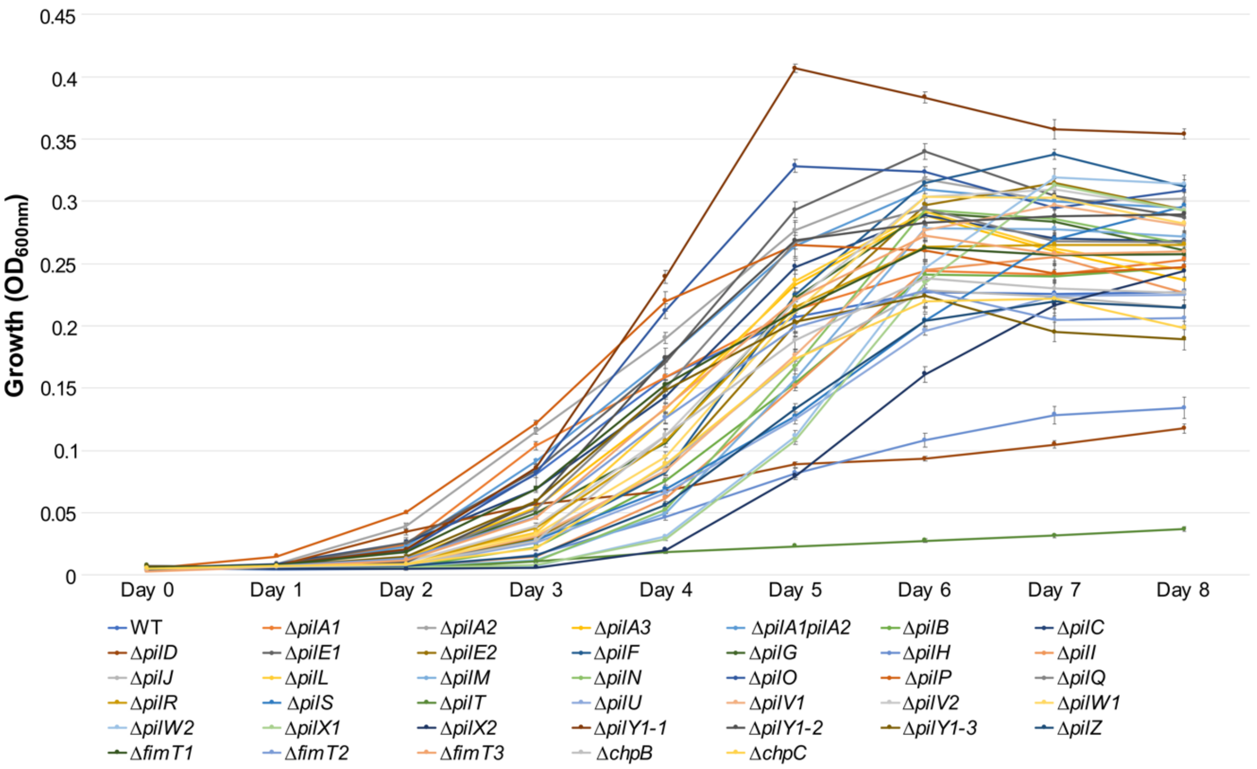


**Fig. S4.** Growth curves of *X. fastidiosa* mutant strains used in this study. Growth curves were generated by culturing bacteria in PD3 broth within 96-well plates and measuring the optical density at 600 nm ($OD_{600nm}$) values each day for 8 days. Data represent means and standard errors (n = three to 15 independent replicates, with eight internal replicates each).

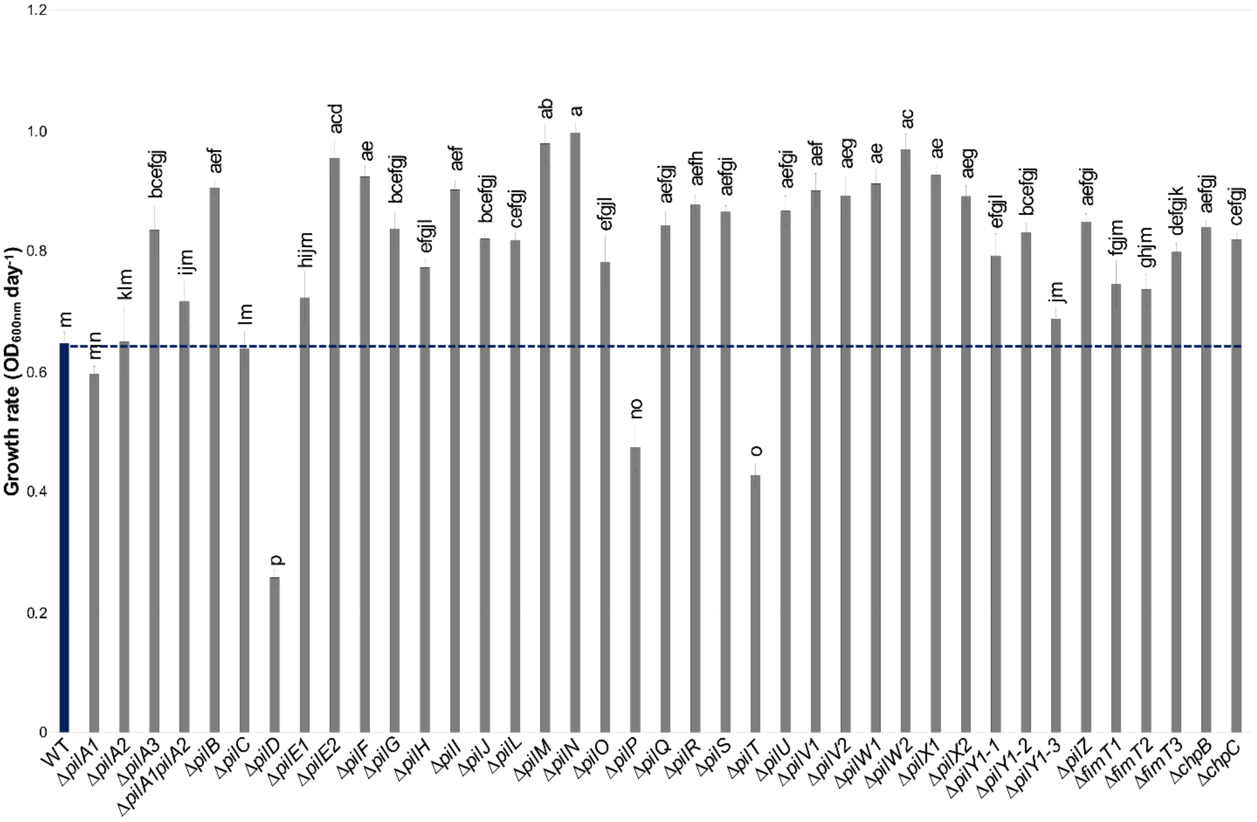


**Fig. S5.** Growth rates of *X. fastidiosa* mutant strains used in this study. Growth rate was calculated from the growth curve at the exponential growth phase (2 to 6 days post inoculation). WT is highlighted in blue, and the dashed blue line indicates the mean value of growth rate for the WT. Data represent means and standard errors. Different letters on top of bars indicate significant difference as analyzed by ANOVA followed by Tukey's HSD multiple comparisons of means ($P$<0.05; n = three to 15 independent replicates with eight internal replicates each).

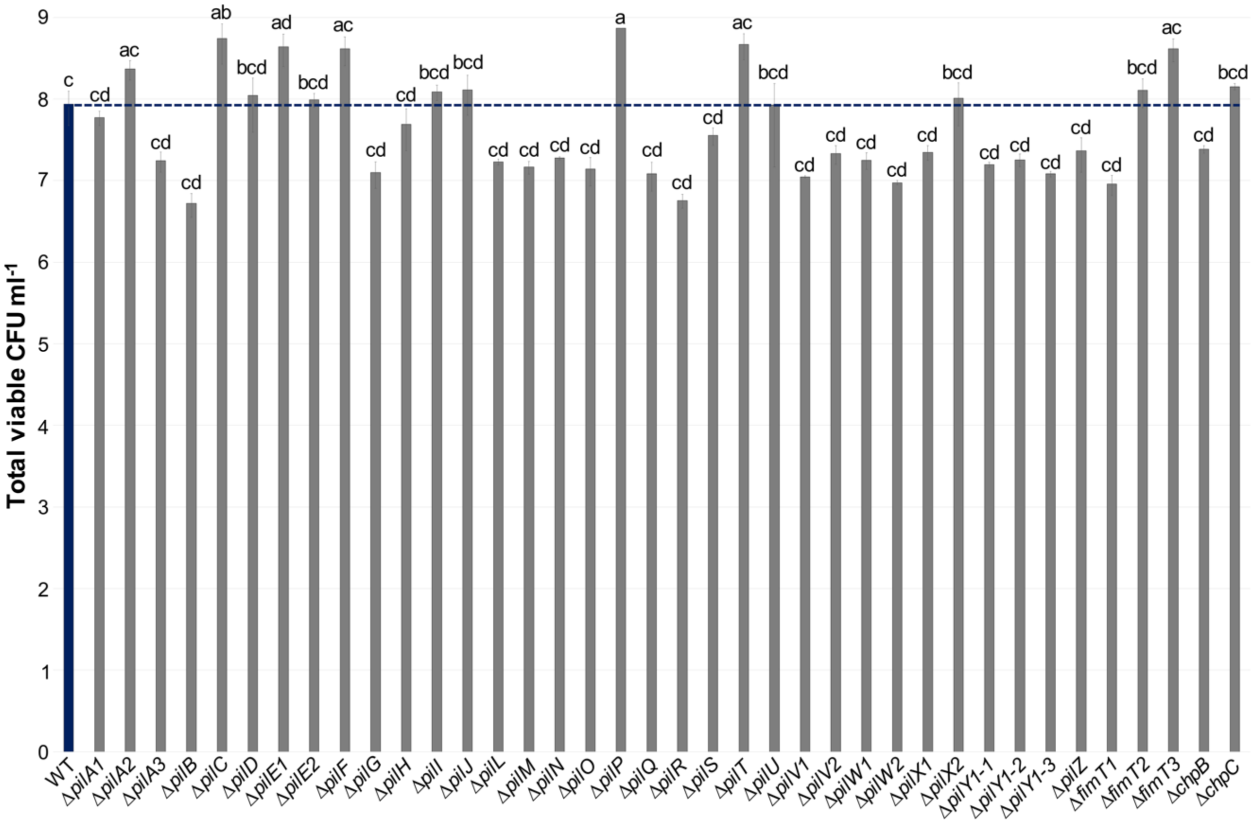


**Fig. S6.** Total number of viable CFU/ml of *X. fastidiosa* mutant strains used in this study after growth during natural competence assays. Total viable CFUs of each *X. fastidiosa* strain obtained during natural competence assays described in figure S1 are shown here. WT is highlighted in blue, and the dashed blue line indicates the mean value of the total number of viable CFU/ml obtained during natural competence assays for the WT. Data represent means and standard errors. Different letters on top of bars indicate significant difference as analyzed by ANOVA followed by Tukey's HSD multiple comparisons of means ($P<0.05$; n = three to 21 independent replicates with two internal replicates each).

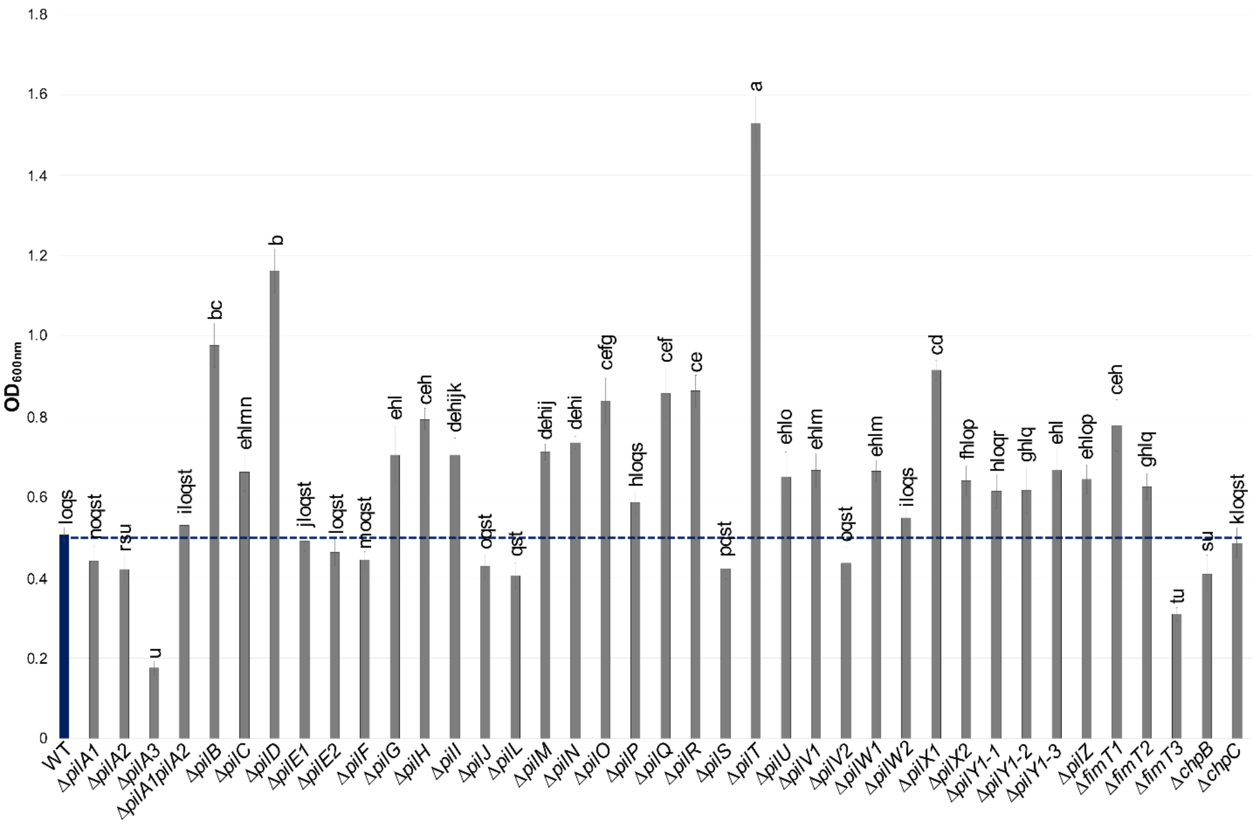


**Fig. S7.** Biofilm formation of *X. fastidiosa* mutant strains used in this study. Biofilm was measured by staining *X. fastidiosa* cells attached to the 96-well plates at the end of the growth curve experiment with crystal violet. WT is highlighted in blue, and the dashed blue line indicates the mean value of biofilm formation for the WT. Data represent means and standard errors. Different letters on top of bars indicate significant difference as analyzed by ANOVA followed by Tukey's HSD multiple comparisons of means ($P$<0.05; n = three to 14 independent replicates with six to eight internal replicates each).

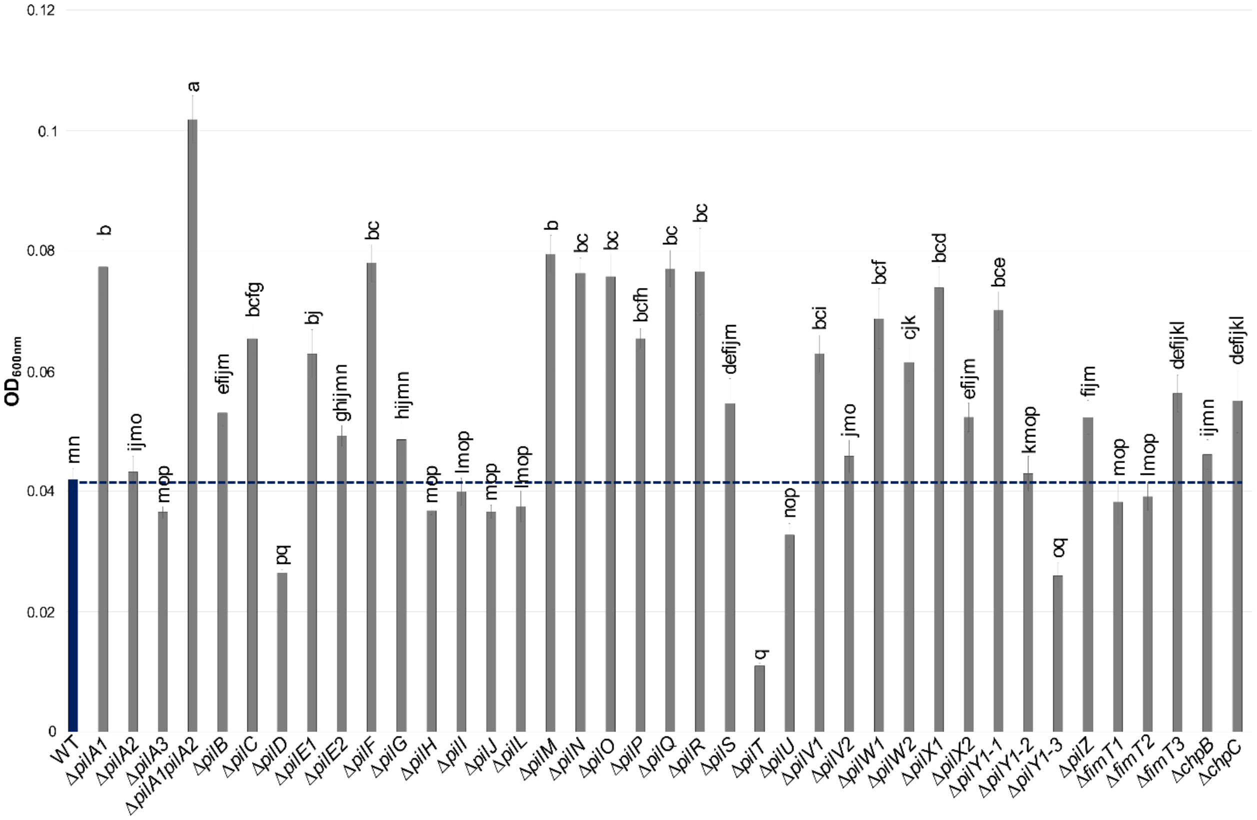


**Fig. S8.** Planktonic growth of *X. fastidiosa* mutant strains used in this study. Planktonic growth was quantified by measuring the optical density at 600 nm ($OD_{600nm}$) values of the supernatant of each *X. fastidiosa* strain at the end of the growth curve experiment. WT is highlighted in blue, and the dashed blue line indicates the mean value of planktonic growth for the WT. Data represent means and standard errors. Different letters on top of bars indicate significant difference as analyzed by ANOVA followed by Tukey's HSD multiple comparisons of means ($P$<0.05; n = three to 14 independent replicates with six to eight internal replicates each).

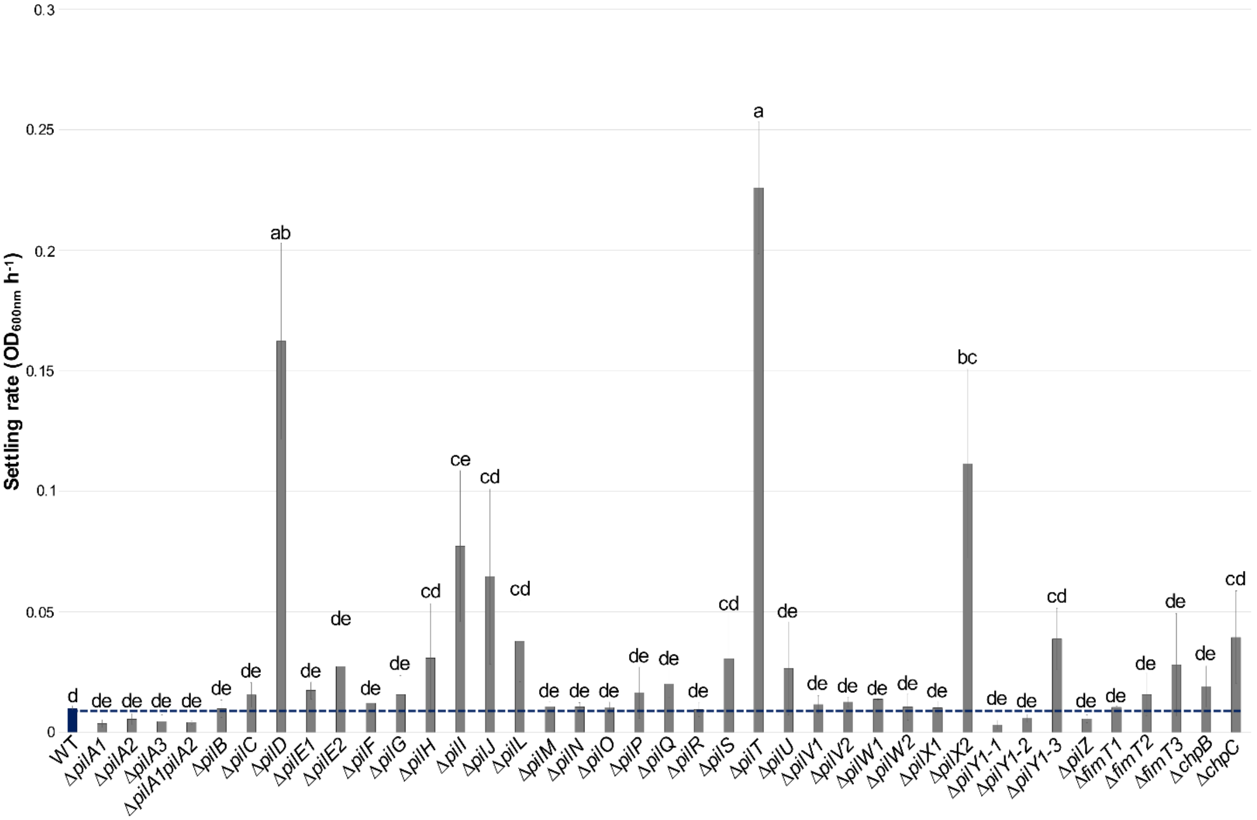


**Fig. S9.** Settling rates of *X. fastidiosa* mutant strains used in this study. Settling rate was measured by suspending ($OD_{600nm}$ = 1.0) the analyzed *X. fastidiosa* strains in a cuvette in 1 ml of PD3 broth and measuring $OD_{600nm}$ values at the initial time point and after 2 hours. WT is highlighted in blue, and the dashed blue line indicates the mean value of settling rate for the WT. Data represent means and standard errors. Different letters on top of bars indicate significant difference as analyzed by ANOVA followed by Tukey's HSD multiple comparisons of means ($P$<0.05; n = three to 15 independent replicates).

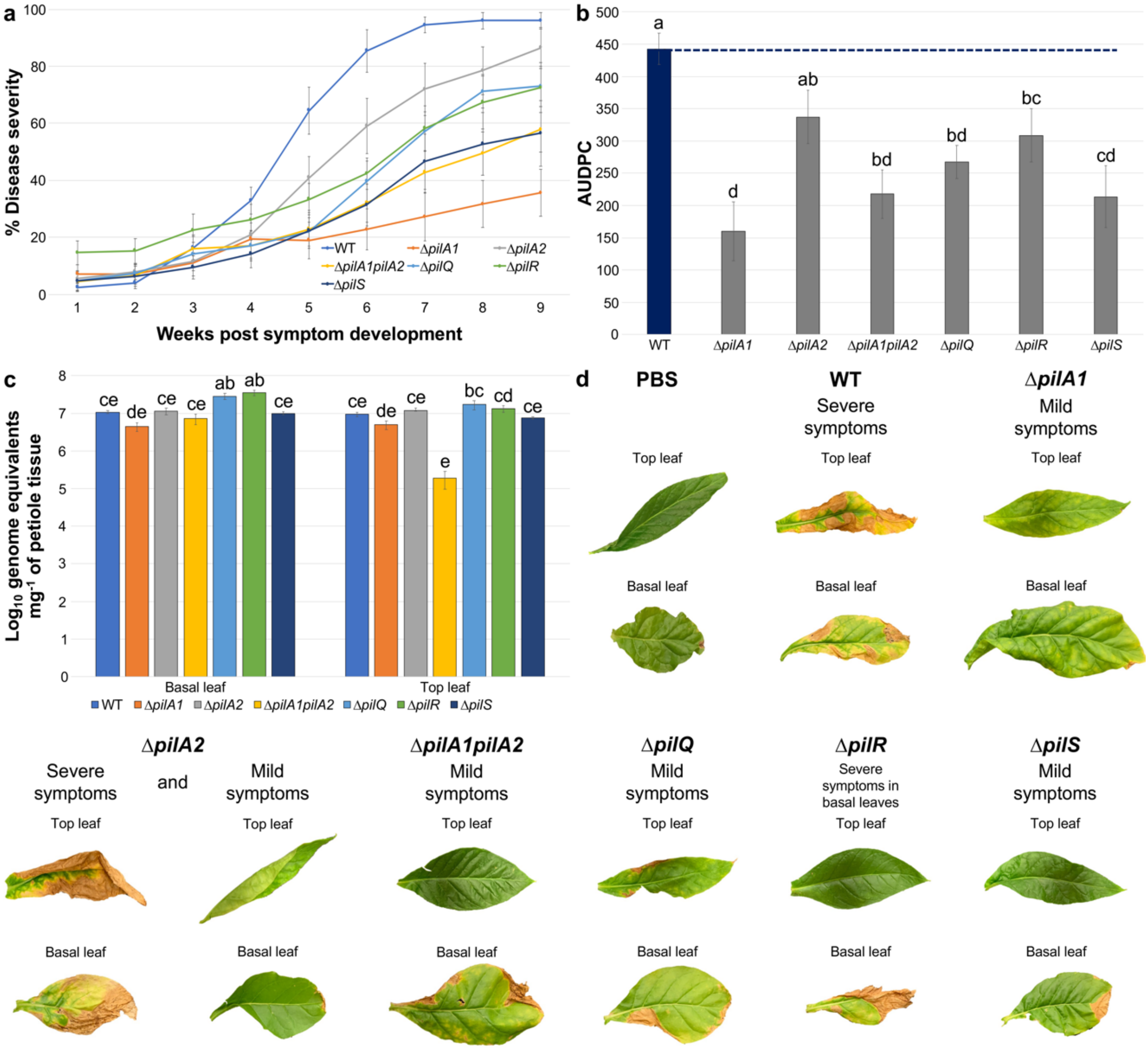


**Fig. S10.** Deletion of *pil* genes negatively affects virulence of *X. fastidiosa*. **a** Disease severity progression over time in inoculated tobacco plants. *X. fastidiosa* WT and mutant strains were inoculated into *Nicotiana tabacum* L. cv. Petite Havana SR1 plants (PBS mock inoculation used as control). Mutant strains chosen for inoculation presented different twitching motility phenotypes, including higher twitching motility (Δ*pilA1* and Δ*pilS*), lower twitching motility (Δ*pilA2*) and non-motility (Δ*pilA1pilA2*, Δ*pilQ* and Δ*pilR*). Leaf scorch symptoms were recorded for measurements of disease incidence and severity once a week during nine weeks after appearance of the first disease symptoms. At the final time point of evaluation, disease incidence in all inoculated plants reached 100%, except for plants inoculated with Δ*pilA1*, which reached 88.88% disease incidence (±11.11%, standard error). On the other hand, disease severity reached 96% in the WT, 35% in Δ*pilA1*, 86% in Δ*pilA2*, 57% in Δ*pilA1pilA2*, 73% in Δ*pilQ*, 72% in Δ*pilR* and 56% in Δ*pilS*. Data represent means and standard errors from two independent experiments (n = seven to ten plants in each independent experiment). **b** Mean AUDPC per treatment group. AUDPC was calculated using data from disease severity over nine weeks after first disease symptom appearance. WT is highlighted in blue, and the dashed blue line indicates the mean value of AUDPC for WT-inoculated plants. AUDPC was significantly lower for plants inoculated with all mutant strains in comparison to WT-inoculated plants, except for Δ*pilA2*. Data represent means and standard errors. Different letters on top of bars indicate significant difference as analyzed by ANOVA followed by Tukey's HSD multiple comparisons of means ($P$<0.05; n = two independent experiments with seven to ten plants each). **c** *X. fastidiosa* in planta population determined by qPCR at the last time point of evaluation. The population of

WT and mutant strains throughout inoculated plants was calculated using petioles of basal and top leaves. ∆*pilA1pilA2* cells showed a defect in colonizing the top leaf of plant hosts, while ∆*pilQ* and ∆*pilR* showed a significant higher population in the basal leaf of infected plants. This shows that mutant strains still move inside the xylem of plant hosts, but absence of deleted genes impairs full symptom development. Data represent means and standard errors. Different letters on top of bars indicate significant difference as analyzed by ANOVA followed by Tukey's HSD multiple comparisons of means ($P$<0.05; n = two independent experiments with three leaf replicates each). **d** Figure panel with representative pictures of leaf scorch symptoms in WT- and mutant strains-inoculated plants (top and basal leaves), as well as control plants (PBS mock inoculation). Only WT-inoculated plants consistently presented severe leaf scorch symptoms. Similar events were captured in two independent experiments.

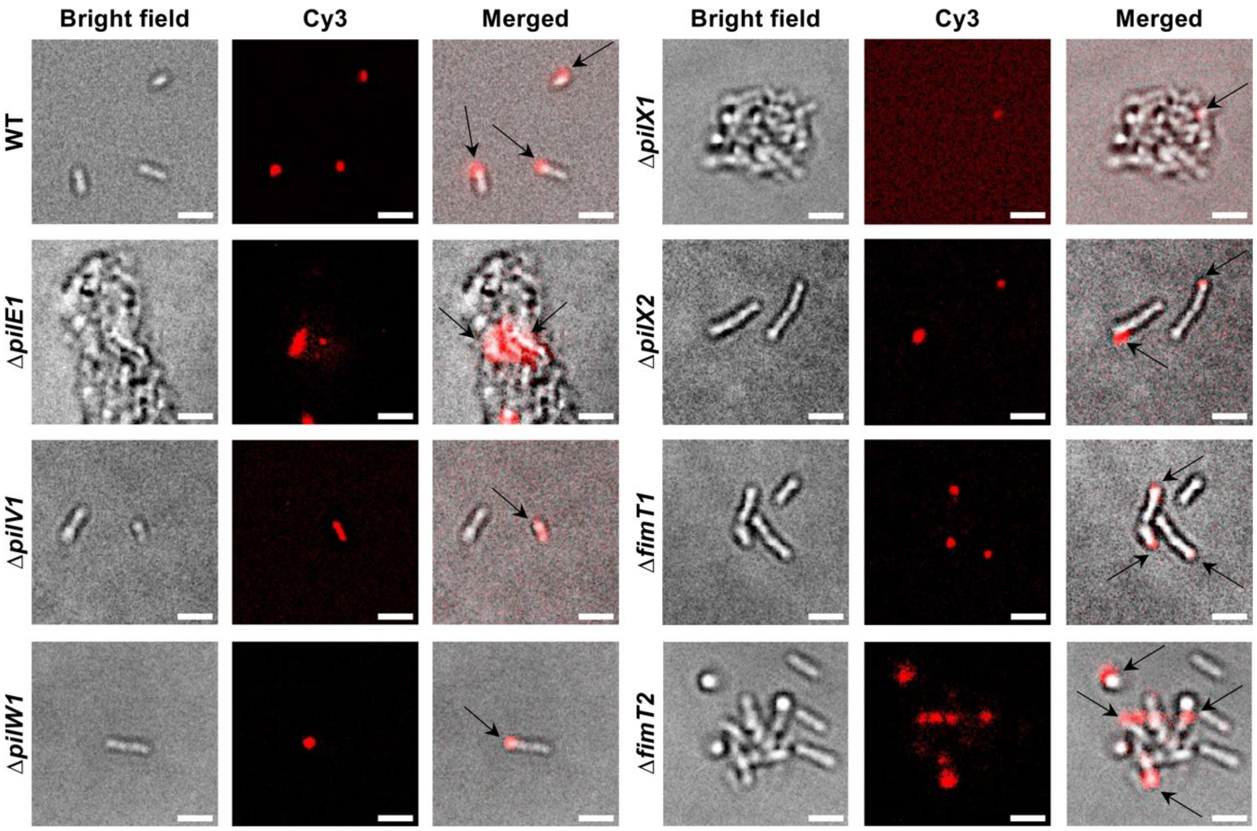


**Fig. S11.** *X. fastidiosa* cells take up Cy-3 labeled DNA into a DNase I resistant state. WT and mutant cells were exposed to Cy-3-labeled pAX1-Cm plasmid (1 µg) for 24 hours, treated with DNase I and DNA foci were observed using a fluorescent microscope. The images shown correspond to the bright field (left), Cy-3 channel (center) and merged fluorescent images (right). In merged fluorescent images, arrows are pointing to fluorescent DNA foci at the poles of cells. Similar events were captured in two to three independent experiments. All evaluated cells presented uptake of Cy-3 labeled DNA. Mutant strains analyzed here correspond to knockout of minor pilins that presented lower recombination through natural competence in comparison to the WT. Δ*fimT1* and Δ*fimT2* were included for comparison with Δ*fimT3*. Images were captured at 100× magnification. Scale bars, 1.5 µm.

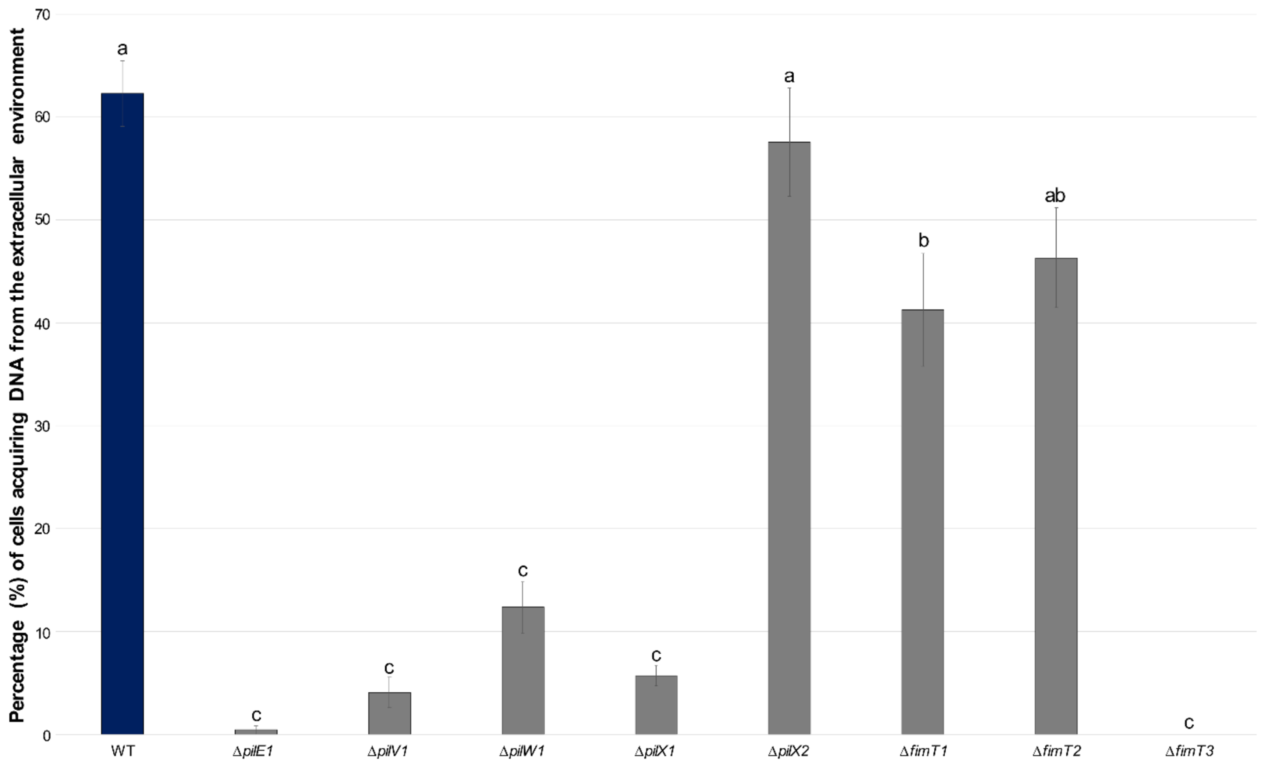


**Fig. S12.** Percentage of *X. fastidiosa* cells that acquired DNA from the extracellular environment during DNA uptake assays in the different analyzed strains. Total cells and cells with DNA foci (Cy-3 labeled pAX1-Cm plasmid) were counted and results are expressed as the percentage of cells with DNA foci. All mutants of minor pilins apart Δ*pilX2* and Δ*fimT2* presented significant lower DNA uptake than WT cells. Data represent means and standard errors. Different letters on top of bars indicate significant difference as analyzed by ANOVA followed by Tukey's HSD multiple comparisons of means ($P$<0.05; n = two to three independent replicates with three to seven technical replicates each).

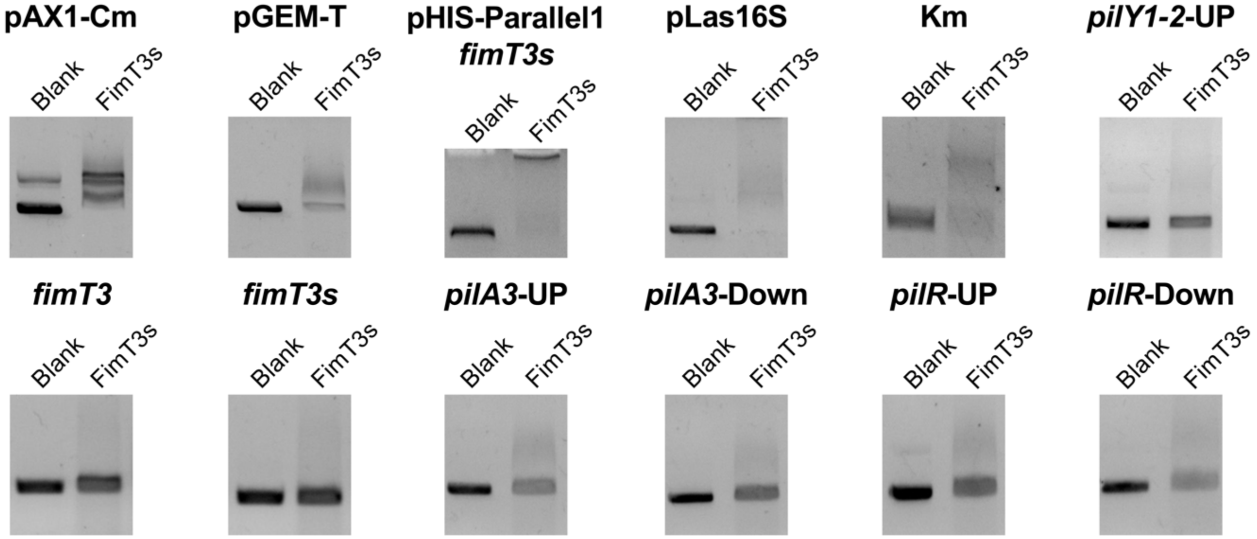


**Fig. S13.** FimT3 DNA binding activity in different DNA sequences. DNA-binding activity of purified FimT3s (5 µM) to different DNA sequences was assessed by agarose EMSA using standard amounts (200 ng) of each DNA sequence. FimT3s was incubated with each DNA sequence for 30 min at 28 ºC and resolved by electrophoresis on a 0.8% agarose gel. Lysis buffer containing 250 µM imidazole, in which proteins were suspended, was included as blank control. DNA binding activity of FimT3s in large sequences (larger than 1,200 bp) is observed as clear shifts in the electrophoretic mobility of bands, while DNA binding activity of FimT3s in small sequences (smaller than 900 bp) is observed as smearing of bands. Amplicon sizes: pAX1-Cm – 4,361 bp; pGEM-T – 3,000 bp; pHIS-Parallel1-*fimT3s* – 5,977 bp; pLas16S – 5,206 bp; Km – 1,202 bp; *pilY1-2*-UP – 988 bp; *fimT3* – 609 bp; *fimT3s* – 519 bp; *pilA3*-UP – 852 bp; *pilA3*-Down – 887 bp; *pilR*-UP – 900 bp; *pilR*-Down – 937 bp. The experiment included DNA sequences with homology to the genome of *X. fastidiosa* TemeculaL (pAX1-Cm, pHIS-Parallel1-*fimT3s*, *pilY1-2*-UP, *fimT3*, *fimT3s*, *pilA3*-UP, *pilA3*-Down, *pilR*-UP and *pilR*-Down), as well as sequences with no apparent homology (pGEM-T, pLas16S and Km resistance cassette). FimT3s did not appear to preferably bind certain DNA sequences, indicating that its DNA-binding ability is non-sequence-specific. *fimT3* – full-length *fimT3* amplified from *X. fastidiosa* TemeculaL; *fimT3s* – soluble portion of *fimT3* amplified from *X. fastidiosa* TemeculaL; sequences labeled with “UP” and “Down” correspond to upstream and downstream sequences, respectively, used to construct the targeting sequences for site-directed mutagenesis of each gene of interest. Similar events were captured in two independent experiments.

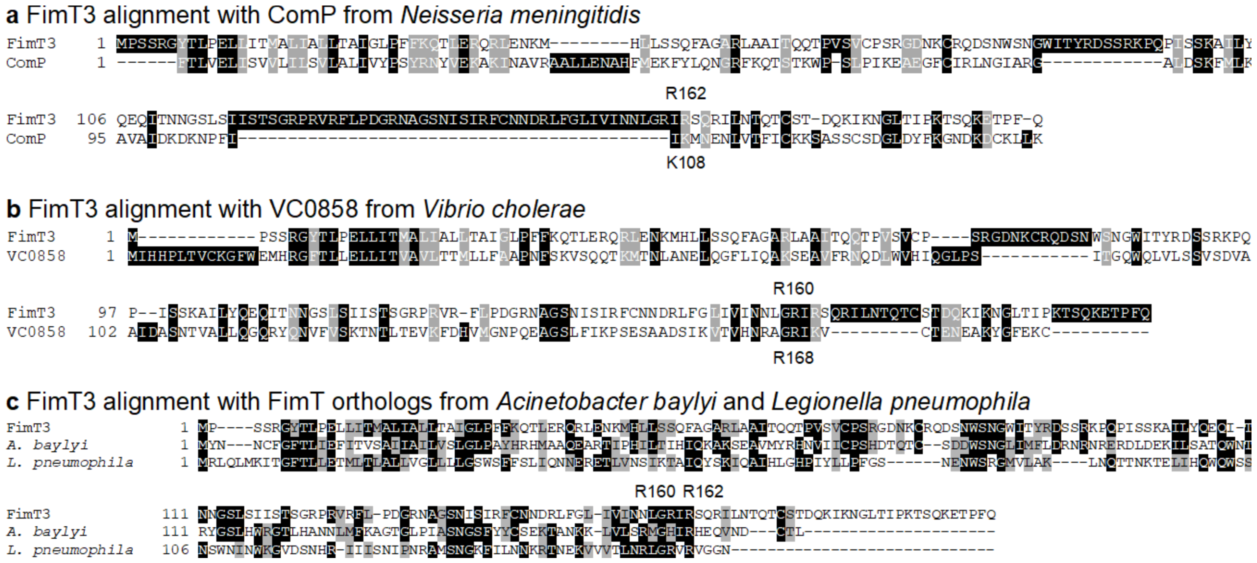


**Fig. S14.** Alignment of FimT3 from *X. fastidiosa* to DNA-binding minor pilins from *Neisseria meningitidis* (ComP), *Vibrio cholerae* (VC0858), *Acinetobacter baylyi* (FimT) and *Legionella pneumophila* (FimT). The sequence of each protein was obtained from NCBI, aligned through T-Coffee, and visualized using BoxShade. Black shading indicates conserved residues; grey shading indicates conservative mutations; and white color indicates divergence among sequences. **a**, **b** and **c** show alignment of FimT3 with ComP, VC0858, and FimT from *A. baylyi* and *L. pneumophila*, respectively. The arginine residue at position 162 (R162) from FimT3 aligned with a lysine of ComP (K108) demonstrated to be essential for the DNA-binding ability of the latter protein. On the other hand, the arginine residue at position 160 (R160) from FimT3 aligned with an arginine of VC0858 (R168) demonstrated to be important for the DNA-binding ability of the *V. cholerae* pilus. In addition, R160 aligned with an arginine residue of FimT from *A. baylyi*, and both R160 and R162 aligned with arginine residues of FimT from *L. pneumophila* that are important to its DNA-binding ability. Alignment of these specific amino acid residues are highlighted by a yellow shading in **a**, **b** and **c**.

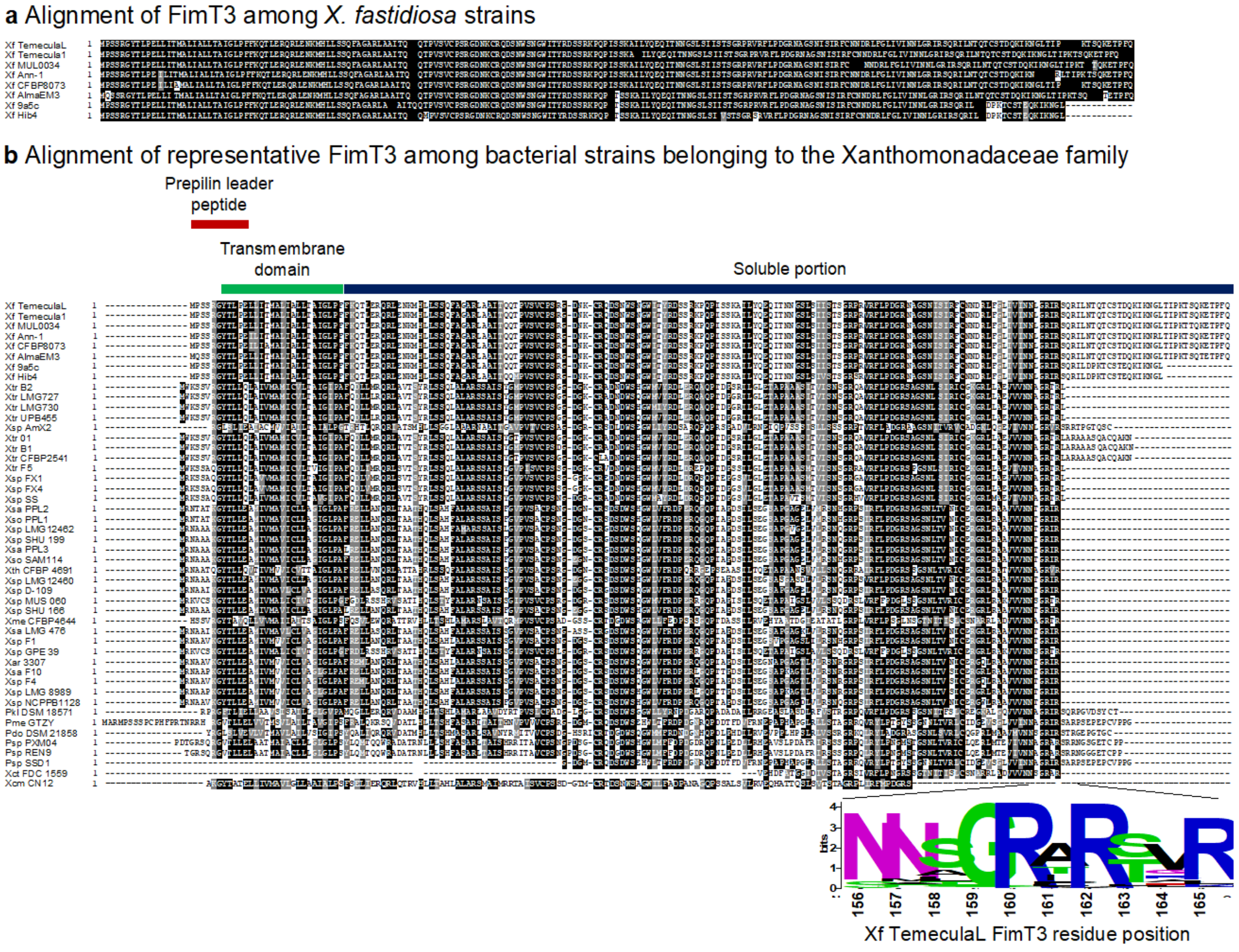


**Fig. S15.** The DNA-binding residues of FimT3 is highly conserved within bacterial members of the Xanthomonadaceae family encoding this protein. The sequence of each protein was downloaded from NCBI, screened for the presence of FimT3 by tblastn, aligned through MAFFT, and visualized using BoxShade. Black shading indicates conserved residues; grey shading indicates conservative mutations; and white color indicates divergence among sequences. **a** Alignment of different FimT3 sequences among *X. fastidiosa* strains. FimT3 is nearly identical in all *X. fastidiosa* strains, with strains from subspecies *pauca* (strains 9a5c and Hib4) presenting the highest divergence (97.25% and 95.6% of identical amino acids, respectively, in comparison to FimT3 from strain TemeculaL). **b** Alignment of representative FimT3 sequences from bacterial members of the Xanthomonadaceae family encoding this protein. Although the percentage of identical amino acids ranged from 15.5% to 100%, most sequences aligned with the arginine amino acid residues at positions 160 and 162 of FimT3 from *X. fastidiosa* strain TemeculaL (highlighted with a yellow shading in the figure). This indicates that these arginine residues are highly conserved within FimT3 sequences. For conciseness of the figure, only representative FimT3 sequences are shown in the alignment. The different portions of FimT3 (prepilin leader peptide, transmembrane domain, and soluble portion) are indicated in the figure. A sequence logo generated using the full alignment of 1,416 FimT3 sequences to highlight the conserved GRxR motif is shown in the bottom of the figure. Abbreviations are the following. ***Pseudoxanthomonas*:** Pdo – *P. dokdonensis*; Pkl – *P. kalamensis*; Pme – *P. mexicana*; Psp – *P.* sp. ***Xanthomonas*:** Xar – *X. arboricola*; Xcm – *X. campestris*; Xct – *X. citri*; Xme – *X. melonis*; Xsa – *X. sacchari*; Xso – *X. sontii*; Xsp – *X.* sp.; Xth – *X. theicola*; Xtr – *X. translucens*. ***Xylella*:** Xf – *X. fastidiosa*.

**Table S1.** List of *pil* and associated genes deleted in *X. fastidiosa* strain TemeculaL.

| Locus ID[a] | Gene | Genome annotation[a] | General description | Reference |
|---|---|---|---|---|
| **Major pilin[b]** | | | | |
| PD1924 | *pilA1* | Type IV fimbrial precursor | Major pilin subunit that composes the main structural component of TFP | 2 |
| PD1926 | *pilA2* | Fimbrial protein | | |
| PD1077 | *pilA3* | Fimbrillin | | |
| **Minor pilin** | | | | |
| PD0024 | *pilE1* | Type IV pilin | Minor pilins responsible for priming the assembly of TFP and promoting the display of the PilY1 adhesin at the tip of the pilus | 49 |
| PD1610 | *pilE2* | Type IV pilin | | |
| PD0020 | *pilV1* | Prepilin leader sequence | | |
| PD1614 | *pilV2* | Type IV pilus modification protein | | |
| PD0021 | *pilW1* | Prepilin type N-terminal cleavage//methylation domain | | |
| PD1613 | *pilW2* | Pilus assembly protein | | |
| PD0022 | *pilX1* | Type IV fimbrial biogenesis protein | | |
| PD1612 | *pilX2* | Type IV fimbrial biogenesis protein | | |
| PD0019 | *fimT1* | Prepilin like leader sequence | Minor pilin that interacts directly with PilA and mediates its connection to the other minor pilins subcomplex (PilVWXY1E) | |
| PD1615 | *fimT2* | Prepilin type N-terminal cleavage/methylation domain | | |
| PD1735 | *fimT3* | Type IV fimbrial biogenesis protein | | |
| **Pilus-associated adhesin** | | | | |
| PD0023 | *pilY1-1* | Type IV fimbrial biogenesis protein | TFP tip adhesin | 45 |
| PD1611 | *pilY1-2* | Type IV fimbrial biogenesis protein | | |
| PD0502 | *pilY1-3* | Type IV fimbrial biogenesis protein | | |
| **Secretin** | | | | |
| PD1691 | *pilQ* | Fimbrial assembly protein | Multimeric outer membrane secretin that forms gated pores from which TFP are extruded | 2 |
| **Pilotin** | | | | |
| PD1623 | *pilF* | Type IV fimbrial biogenesis protein/stability protein | Pilotin protein required for the outer membrane localization and assembly of the multimeric PilQ secretin | 59 |
| **Platform protein** | | | | |
| PD1923 | *pilC* | Fimbrial assembly protein | Platform protein that mediates TFP assembly through the polymerization and depolymerization ATPases | 5 |

| | | | | |
|---|---|---|---|---|
| **Alignment protein** | | | | |
| PD1695 | *pilM* | Fimbrial assembly membrane protein | Alignment subcomplex that is involved in TFP assembly and stabilization by linking the outer membrane secretin pore subcomplex (PilF and PilQ) to the inner membrane motor subcomplex (PilB, PilC, PilD, PilT and PilU) | 85 |
| PD1694 | *pilN* | Fimbrial assembly membrane protein | | |
| PD1693 | *pilO* | Fimbrial assembly membrane protein | | |
| PD1692 | *pilP* | Fimbrial assembly protein | | |
| **Assembly ATPase** | | | | |
| PD1927 | *pilB* | Pilus biogenesis protein | TFP polymerization (assembly) ATPase; promotes TFP extension | 2 |
| **Retraction ATPase** | | | | |
| PD1147 | *pilT* | Type IV fimbrial biogenesis protein | TFP depolymerization (disassembly) ATPases; promote TFP retraction | 2 |
| PD1148 | *pilU* | Type IV fimbrial biogenesis protein | | |
| **Prepilin peptidase** | | | | |
| PD1922 | *pilD* | Type IV prepilin leader peptidase | Prepilin peptidase that cleaves the leader peptide of all pilins (major and minor) at the cytoplasmic milieu of the inner membrane and methylates the mature pilin | 50,86 |
| **Two-component regulatory system** | | | | |
| PD1928 | *pilR* | Two-component system, regulatory protein | Cytoplasmic response regulator that binds to the promoter of *pilA* (together with $\sigma^{54}$) to activate transcription of the major pilin | 25,26 |
| PD1929 | *pilS* | Two-component system, sensor protein | Sensor kinase that interacts directly with PilA for pilin transcription autoregulation | 27 |
| **Chemotaxis protein** | | | | |
| PD0845 | *pilG* | Pilus protein | Chemosensory system composed by a transmembrane chemoreceptor (PilJ), a histidine kinase (PilL), a methylesterase (ChpB), response regulators (PilG and PilH) and coupling proteins (PilI and ChpC), which has been shown to affect twitching motility | 36,38 |
| PD1632 | *pilH* | Regulatory protein | | |
| PD0846 | *pilI* | Type IV fimbrial biogenesis protein | | |
| PD0847 | *pilJ* | Type IV fimbrial biogenesis protein | | |
| PD0848 | *pilL* | Chemotaxis-related protein kinase | | |
| PD0849 | *chpB* | Chemotaxis response regulator protein | | |
| PD0850 | *chpC* | Chemotaxis protein | | |

| | | | | |
|---|---|---|---|---|
| **Signal transduction** | | | | |
| PD1497 | *pilZ* | Type IV fimbriae assembly protein | Regulatory protein associated with TFP extension in a possible c-di-GMP (3′,5′-cyclic-di-guanylate) -dependent manner | [8,87] |

[a]Locus IDs and genome annotations are from the genome of *X. fastidiosa* subsp. *fastidiosa* reference strain Temecula1, GenBank Accession number AE009442.1. Genome annotation was visualized using Geneious (Biomatters Ltd.).
[b]Genes are organized by functional categories in the table.

**Table S2.** Genomic organization of *pil* and associated genes in *X. fastidiosa* TemeculaL.

| Gene(s)[a] | Operon[b] | DNA strand |
|---|---|---|
| *fimT1* (PD0019), *pilV1* (PD0020), *pilW1* (PD0021), *pilX1* (PD0022), *pilY1-1* (PD0023), *pilE1* (PD0024) | PD0019→PD0020→PD0021→PD0022→PD0023→PD0024 | Leading |
| *pilY1-3* (PD0502) | Not in operon | Leading |
| *pilG* (PD0845), *pilI* (PD0846), *pilJ* (PD0847), *cheA/pilL* (PD0848), *cheB/chpB* (PD0849), *cheW/chpC* (PD0850) | PD0845→PD0846→PD0847→PD0848→PD0849→PD0850 | Leading |
| *pilA3* PD1077 | Not in operon | Lagging |
| *pilT* (PD1147) | Not in operon | Leading |
| *pilU* (PD1148) | Not in operon | Leading |
| pilZ (PD1497), **PD1498**, **PD1499** | PD1499→PD1498→PD1497 | Lagging |
| *pilE2* (PD1610), *pilY1-2* (PD1611) | PD1611→PD1610 | Lagging |
| *pilX2* (PD1612), *pilW2* (PD1613), *pilV2* (PD1614), *fimT2* (PD1615) | PD1615→PD1614→PD1613→PD1612 | Lagging |
| *pilF* PD1623, **PD1622** | PD1623→PD1622 | Lagging |
| *pilH* (PD1632), **PD1631** | PD1631→PD1632 | Leading |
| *pilQ* (PD1691), *pilP* (PD1692), *pilO* (PD1693), *pilN* (PD1694), *pilM* (PD1695) | PD1695→PD1694→PD1693→PD1692→ PD1691 | Lagging |
| *fimT3* (PD1735) | Not in operon | Leading |
| **PD1921**, *pilD* (PD1922), *pilC* (PD1923) | PD1923→PD1922→PD1921 | Lagging |
| *pilA1* (PD1924) | Not in operon | Leading |
| *pilA2* (PD1926) | Not in operon | Leading |
| *pilB* (PD1927) | Not in operon | Leading |
| *pilR* (PD1928) | Not in operon | Lagging |
| *pilS* (PD1929) | Not in operon | Lagging |

[a]Genes in bold have no apparent effect on type IV pili and/or natural transformation. They are included in the list because they are located within the same operon of *pil* genes.
[b]Operon prediction was retrieved from elsewhere [88].

**Table S3.** Pearson's correlation results for the analyzed phenotypic traits of *X. fastidiosa*.

| Phenotypic traits[a] | Recombination frequency | Twitching motility | Biofilm formation | Planktonic growth | Growth rate | Settling rate | Total viable CFU/ml[b] |
|---|---|---|---|---|---|---|---|
| **Recombination frequency** | - | **$R^2$ = 0.56 *P*<0.001** | $R^2$ = -0.23 *P* = 0.15 | $R^2$ = -0.14 *P* = 0.38 | $R^2$ = 0.02 *P* = 0.89 | $R^2$ = -0.11 *P* = 0.47 | $R^2$ = -0.27 *P* = 0.09 |
| **Twitching motility** | **$R^2$ = 0.56 *P*<0.001** | - | **$R^2$ = -0.54 *P*<0.001** | $R^2$ = -0.14 *P* = 0.38 | $R^2$ = 0.09 *P* = 0.58 | $R^2$ = -0.14 *P* = 0.38 | $R^2$ = -0.10 *P* = 0.51 |
| **Biofilm formation** | $R^2$ = -0.23 *P* = 0.15 | **$R^2$ = -0.54 *P*<0.001** | - | $R^2$ = -0.16 *P* = 0.30 | **$R^2$ = -0.34 *P*<0.05** | **$R^2$ = 0.59 *P*<0.001** | $R^2$ = 0.01 *P* = 0.94 |
| **Planktonic growth** | $R^2$ = -0.14 *P* = 0.38 | $R^2$ = -0.14 *P* = 0.38 | $R^2$ = -0.16 *P* = 0.30 | - | **$R^2$ = 0.34 *P*<0.05** | **$R^2$ = -0.55 *P*<0.001** | $R^2$ = 0 *P* = 1.0 |
| **Growth rate** | $R^2$ = 0.02 *P* = 0.89 | $R^2$ = 0.09 *P* = 0.58 | **$R^2$ = -0.34 *P*<0.05** | **$R^2$ = 0.34 *P*<0.05** | - | **$R^2$ = -0.51 *P*<0.001** | **$R^2$ = -0.49 *P*<0.005** |
| **Settling rate** | $R^2$ = -0.11 *P* = 0.47 | $R^2$ = -0.14 *P* = 0.38 | **$R^2$ = 0.59 *P*<0.001** | **$R^2$ = -0.55 *P*<0.001** | **$R^2$ = -0.51 *P*<0.001** | - | $R^2$ = 0.24 *P* = 0.12 |
| **Total viable CFU/ml[a]** | $R^2$ = -0.27 *P* = 0.09 | $R^2$ = -0.10 *P* = 0.51 | $R^2$ = 0.01 *P* = 0.94 | $R^2$ = 0 *P* = 1.0 | **$R^2$ = -0.49 *P*<0.005** | $R^2$ = 0.24 *P* = 0.12 | - |

[a]Results are shown as correlation coefficients ($R^2$) and *p*-values (*P*). Results in bold show significant correlations (either positive or negative) among phenotypes.
[b]Total viable CFU/ml obtained in natural competence assays.

**Table S4.** DNA-binding probability of amino acid residues from FimT3.

| Amino acid | Position | DNA-binding probability[a] |
|---|---|---|
| R (arginine) | 160 | 0.8941 |
| R (arginine) | 162 | 0.8199 |
| W (tryptophan) | 81 | 0.8161 |
| K (lysine) | 177 | 0.784 |
| N (asparagine) | 180 | 0.7829 |
| R (arginine) | 93 | 0.7772 |
| G (glycine) | 159 | 0.7689 |
| K (lysine) | 94 | 0.7577 |
| K (lysine) | 179 | 0.7566 |
| G (glycine) | 84 | 0.7465 |
| R (arginine) | 123 | 0.7433 |
| R (arginine) | 54 | 0.7417 |
| S (serine) | 188 | 0.7412 |

[a]The DNA-binding probability of each amino acid residue from FimT3 of *X. fastidiosa* strain TemeculaL was calculated using the DRNApred webserver [89] and ranges from 0 to 1.0. Only amino acid residues with a DNA-binding probability above 0.74 are shown in the table.

**Table S5.** Homologs of FimT3 are found in members of the Xanthomonadaceae family.

| Bacterial species | Number of strains encoding FimT3[a] | Identical amino acids (%)[b] |
|---|---|---|
| ***Xylella*** | | |
| *X. fastidiosa* | 121 | 95.60 – 100 |
| *X. taiwanensis* | 9 | 80 |
| ***Coralloluteibacterium*** | | |
| *C. stylophorae* | 1 | 37.13 |
| ***Luteimonas*** | | |
| *L. aestuarii* | 1 | 36.05 |
| *L. arsenica* | 1 | 34.5 |
| *L. deserti* | 1 | 38.82 |
| *L. fraxinea* | 2 | 39.18 |
| *L. mephitis* | 1 | 38.01 |
| *L. padinae* | 1 | 35.5 |
| *L.* sp. | 9 | 33.73 – 39.18 |
| *L. terrae* | 1 | 36.47 |
| *L. terricola* | 1 | 36.75 |
| *L. yindakuii* | 1 | 36.31 |
| ***Lysobacter*** | | |
| *L.* sp. | 1 | 28.92 |
| ***Pseudoxanthomonas*** | | |
| *P. beigongshangi* | 1 | 41.38 |
| *P. composti* | 1 | 37.39 |
| *P. daejeonensis* | 2 | 40.48 |
| *P. gei* | 1 | 44.31 |
| *P. helianthi* | 1 | 27.88 |
| *P. japonensis* | 2 | 36.05 – 36.63 |
| *P. kaohsiungensis* | 1 | 40.83 |
| *P. koreensis* | 1 | 46.01 |
| *P. mexicana* | 7 (1 with no GRxR motif) | 23.15 – 43.75 |
| *P. sacheonensis* | 1 | 40.23 |
| *P. sangjuensis* | 1 | 29.81 |
| *P.* sp. | 19 | 35.84 – 42.61 |
| *P. spadix* | 3 | 38.6 – 42.44 |
| *P. suwonensis* | 2 | 41.32 |
| *P. taiwanensis* | 1 | 41.92 |
| *P. winnipegensis* | 11 | 35.84 – 37.21 |
| *P. wuyuanensis* | 2 | 42.94 |
| *P. yeongjuensis* | 1 | 41.28 |
| ***Rehaibacterium*** | | |
| *R. terrae* | 1 | 27.71 |
| ***Stenotrophomonas*** | | |
| *S. acidaminiphila* | 7 | 34.52 – 35.71 |
| *S. humi* | 1 | 34.68 |
| *S. maltophilia* | 50 | 30.36 – 39.16 |
| *S. nitritireducens* | 1 | 34.73 |
| *S. pictorum* | 2 | 36.53 |
| *S.* sp. | 7 | 32.56 – 48.84 |
| ***Vulcaniibacterium*** | | |
| *V. gelatinicum* | 2 (1 with no GRxR motif) | 15.5 – 28.9 |
| *V. thermophilum* | 2 | 33.33 |
| ***Xanthomonas*** | | |
| *X. albilineans* | 17 | 44.32 |

| | | |
|---|---|---|
| *X. alfalfae* | 1 | 44.25 |
| *X. arboricola* | 96 (1 with no GRxR motif) | 27.65 – 46.63 |
| *X. axonopodis* | 63 (26 with no GRxR motif) | 27.54 – 44.83 |
| *X. bromi* | 2 | 45.4 |
| *X. campestris* | 128 (2 with no GRxR motif) | 28.4 – 49.43 |
| *X. cannabis* | 3 | 43.1 – 44.77 |
| *X. cassavae* | 2 | 45.71 |
| *X. citri* | 31 (6 with no GRxR motif) | 27.81 – 48.31 |
| *X. codiaei* | 1 | 43.02 |
| *X. dyei* | 3 | 29.34 – 42.44 |
| *X. euroxanthea* | 2 | 41.67 – 41.95 |
| *X. euvesicatoria* | 11 (1 with no GRxR motif) | 28.4 – 45.8 |
| *X. floridensis* | 1 | 43.93 |
| *X. fragariae* | 56 | 34.91 – 35.5 |
| *X. hortorum* | 73 | 32.54 – 43.68 |
| *X. hyacinthi* | 1 | 37.95 |
| *X. maliensis* | 1 | 42.69 |
| *X. melonis* | 3 | 45.93 |
| *X. nasturtii* | 19 | 26.04 – 43.1 |
| *X. oryzae* | 404 | 42.53 – 43.79 |
| *X. perforans* | 19 (3 with no GRxR motif) | 28.57 – 45.86 |
| *X. populi* | 1 | 41.95 |
| *X. prunicola* | 3 | 43.68 |
| *X. sacchari* | 5 | 44.32 – 44.51 |
| *X. sontii* | 5 | 46.02 – 46.82 |
| *X.* sp. | 34 | 27.81 – 48.86 |
| *X. theicola* | 2 | 46.82 |
| *X. translucens* | 69 | 51.7 – 52.27 |
| *X. vasicola* | 67 | 29.34 – 44.83 |
| *X. vesicatoria* | 11 | 28.24 – 44.19 |

[a]A total of 3,0001 genomes of bacterial species belonging to the Xanthomonadaceae family were downloaded from NCBI on May 25$^{th}$, 2022. In the table only the number of genomes from bacterial species that encode homologs of FimT3 are shown.

[b]Sequence alignments to FimT3 of *X. fastidiosa* strain TemeculaL were produced using Clustal Omega (Clustal 12.1).

**Table S6.** Bacterial strains and plasmids used in this study.

| Strain name in manuscript or plasmid | Genotype or description | Source |
| --- | --- | --- |
| ***Xylella fastidiosa*** | | |
| WT | Wild-type *X. fastidiosa* subsp. *fastidiosa* strain TemeculaL | [74] |
| Δ*pilA1* | *X. fastidiosa* strain TemeculaL with chromosomal *pilA1* (PD1924) deletion, $Km^R$ | [21] |
| Δ*pilA2* | *X. fastidiosa* strain TemeculaL with chromosomal *pilA2* (PD1926) deletion, $Km^R$ | This study |
| Δ*pilA3* | *X. fastidiosa* strain TemeculaL with chromosomal *pilA3* (PD1077) deletion, $Km^R$ | This study |
| Δ*pilA1pilA2* | *X. fastidiosa* strain TemeculaL with chromosomal *pilA1* and *pilA2* (PD1924 and PD1926, respectively) deletion, $Km^R$, $Cm^R$ | [21] |
| Δpi*lB* | *X. fastidiosa* strain TemeculaL with chromosomal *pilB* (PD1927) deletion, $Km^R$ | This study |
| Δ*pilC* | *X. fastidiosa* strain TemeculaL with chromosomal *pilC* (PD1923) deletion, $Km^R$ | This study |
| Δ*pilD* | *X. fastidiosa* strain TemeculaL with chromosomal *pilD* (PD1922) deletion, $Km^R$ | This study |
| Δ*pilE1* | *X. fastidiosa* strain TemeculaL with chromosomal *pilE1* (PD0024) deletion, $Km^R$ | This study |
| Δ*pilE2* | *X. fastidiosa* strain TemeculaL with chromosomal *pilE2* (PD1610) deletion, $Km^R$ | This study |
| Δ*pilF* | *X. fastidiosa* strain TemeculaL with chromosomal *pilF* (PD1623) deletion, $Km^R$ | This study |
| Δ*pilG* | *X. fastidiosa* strain TemeculaL with chromosomal *pilG* (PD0845) deletion, $Km^R$ | This study |
| Δ*pilH* | *X. fastidiosa* strain TemeculaL with chromosomal *pilH* (PD1632) deletion, $Km^R$ | This study |
| Δ*pilI* | *X. fastidiosa* strain TemeculaL with chromosomal *pilI* (PD0846) deletion, $Km^R$ | This study |
| Δ*pilJ* | *X. fastidiosa* strain TemeculaL with chromosomal *pilJ* (PD0847) deletion, $Km^R$ | This study |
| Δ*pilL* | *X. fastidiosa* strain TemeculaL with chromosomal *pilL* (PD0848) deletion, $Km^R$ | This study |
| Δ*pilM* | *X. fastidiosa* strain TemeculaL with chromosomal *pilM* (PD1695) deletion, $Km^R$ | This study |
| Δ*pilN* | *X. fastidiosa* strain TemeculaL with chromosomal *pilN* (PD1694) deletion, $Km^R$ | This study |
| Δ*pilO* | *X. fastidiosa* strain TemeculaL with chromosomal *pilO* (PD1693) deletion, $Km^R$ | This study |
| Δ*pilP* | *X. fastidiosa* strain TemeculaL with chromosomal *pilP* (PD1694) deletion, $Km^R$ | This study |
| Δ*pilQ* | *X. fastidiosa* strain TemeculaL with chromosomal *pilQ* (PD1691) deletion, $Km^R$ | This study |
| Δ*pilR* | *X. fastidiosa* strain TemeculaL with chromosomal *pilR* (PD1928) deletion, $Km^R$ | This study |
| Δ*pilS* | *X. fastidiosa* strain TemeculaL with chromosomal *pilS* (PD1929) deletion, $Km^R$ | This study |
| Δ*pilT* | *X. fastidiosa* strain TemeculaL with chromosomal *pilT* (PD1147) deletion, $Km^R$ | This study |
| Δ*pilU* | *X. fastidiosa* strain TemeculaL with chromosomal *pilU* (PD1148) deletion, $Km^R$ | This study |
| Δ*pilV1* | *X. fastidiosa* strain TemeculaL with chromosomal *pilV1* (PD0020) deletion, $Km^R$ | This study |

| | | |
|---|---|---|
| Δ*pilV2* | *X. fastidiosa* strain TemeculaL with chromosomal *pilV2* (PD1614) deletion, Km$^R$ | This study |
| Δ*pilW1* | *X. fastidiosa* strain TemeculaL with chromosomal *pilW1* (PD0021) deletion, Km$^R$ | This study |
| Δ*pilW2* | *X. fastidiosa* strain TemeculaL with chromosomal *pilW2* (PD1613) deletion, Km$^R$ | This study |
| Δ*pilX1* | *X. fastidiosa* strain TemeculaL with chromosomal *pilX1* (PD0022) deletion, Km$^R$ | This study |
| Δ*pilX2* | *X. fastidiosa* strain TemeculaL with chromosomal *pilX2* (PD1612) deletion, Km$^R$ | This study |
| Δ*pilY1-1* | *X. fastidiosa* strain TemeculaL with chromosomal *pilY1-1* (PD0023) deletion, Km$^R$ | This study |
| Δ*pilY1-2* | *X. fastidiosa* strain TemeculaL with chromosomal *pilY1-2* (PD1611) deletion, Km$^R$ | This study |
| Δ*pilY1-3* | *X. fastidiosa* strain TemeculaL with chromosomal *pilY1-3* (PD0502) deletion, Km$^R$ | This study |
| Δ*pilZ* | *X. fastidiosa* strain TemeculaL with chromosomal *pilZ* (PD1497) deletion, Km$^R$ | This study |
| Δ*fimT1* | *X. fastidiosa* strain TemeculaL with chromosomal *fimT1* (PD0019) deletion, Km$^R$ | This study |
| Δ*fimT2* | *X. fastidiosa* strain TemeculaL with chromosomal *fimT2* (PD1615) deletion, Km$^R$ | This study |
| Δ*fimT3* | *X. fastidiosa* strain TemeculaL with chromosomal *fimT3* (PD1735) deletion, Km$^R$ | This study |
| Δ*chpB* | *X. fastidiosa* strain TemeculaL with chromosomal *chpB* (PD0849) deletion, Km$^R$ | This study |
| Δ*chpC* | *X. fastidiosa* strain TemeculaL with chromosomal *chpC* (PD0850) deletion, Km$^R$ | This study |
| ***Escherichia coli*** | | |
| Dh5α | *fhuA2 Δ(argF-lacZ)U169 phoA glnV44 Φ80 Δ(lacZ)M15 gyrA96 recA1 relA1 endA1 thi-1 hsdR17* | New England Biolabs |
| EAM1 | DH5α derivative; Sp$^r$ St$^r$*attP*$_{HK022}$::(P$_{LlacO-1}$-PD1607) Expresses the *X. fastidiosa* subsp. *fastidiosa* strain Temecula1 DNA methylase | [76] |
| BL21(DE3) | *fhuA2 [lon] ompT gal (λ DE3) [dcm] ΔhsdS λ DE3 = λ sBamHIo ΔEcoRI-B int::(lacI::PlacUV5::T7 gene1) i21 Δnin5* | New England Biolabs |
| Dh5α-pUC4K | *E. coli* Dh5α bearing the pUC4K plasmid | [21] |
| EAM1-pAX1-Cm | *E. coli* EAM1 bearing the pAX1-Cm plasmid | [79] |
| Dh5α-pHIS-Parallel1-*fimT1s* | *E. coli* Dh5α bearing the pHIS-Parallel1-*fimT1s* plasmid | This study |
| Dh5α-pHIS-Parallel1-*fimT2s* | *E. coli* Dh5α bearing the pHIS-Parallel1-*fimT2s* plasmid | This study |
| Dh5α-pHIS-Parallel1-*fimT3s* | *E. coli* Dh5α bearing the pHIS-Parallel1-*fimT3s* plasmid | This study |
| Dh5α-pHIS-Parallel1-*fimT3s*-R160AR162A | *E. coli* Dh5α bearing the pHIS-Parallel1-*fimT3s*-R160AR162A plasmid | This study |
| BL21(DE3)-pHIS-Parallel1-*fimT1s* | *E. coli* BL21(DE3) bearing the pHIS-Parallel1-*fimT1s* plasmid | This study |
| BL21(DE3)-pHIS-Parallel1-*fimT2s* | *E. coli* BL21(DE3) bearing the pHIS-Parallel1-*fimT2s* plasmid | This study |

| | | |
|---|---|---|
| BL21(DE3)-pHIS-Parallel1-*fimT3s* | *E. coli* BL21(DE3) bearing the pHIS-Parallel1-*fimT3s* plasmid | This study |
| BL21(DE3)-pHIS-Parallel1-*fimT3s*-R160AR162A | *E. coli* BL21(DE3) bearing the pHIS-Parallel1-*fimT3s*-R160AR162A plasmid | This study |
| Dh5α-pLas16S | *E. coli* Dh5α bearing the pLas16S plasmid | [90] |
| **Plasmids** | | |
| pUC4K | Donor of kanamycin resistance cassette, $Km^R$ $Amp^R$ | [91] |
| pAX1-Cm | pGEM-T derivative; contains multiple cloning site Plasmid that recombines into the neutral site 1 (NS1) of *X. fastidiosa* and inserts a chloramphenicol resistance cassette | [75] |
| pHIS-Parallel1 | Cloning vector, f1 ori *lacI* T7 promoter ($P_{T7}$) $Amp^R$ | [92] |
| pHIS-Parallel1-*fimT1s* | $Amp^R$, $P_{T7}$-*fimT1s* | This study |
| pHIS-Parallel1-*fimT2s* | $Amp^R$, $P_{T7}$-*fimT2s* | This study |
| pHIS-Parallel1-*fimT3s* | $Amp^R$, $P_{T7}$-*fimT3s* | This study |
| pHIS-Parallel1-*fimT3s*-R160AR162A | $Amp^R$, $P_{T7}$-*fimT3s*-R160AR162A | This study |
| pLas16S[a] | Plasmid standard to quantify the population of '*Candidatus* Liberibacter asiaticus' as genome equivalents through qPCR | [90] |
| pGEM-T | Cloning vector, f1 ori *lacZ* $Amp^R$ | Promega |

$Km^R$ – kanamycin-resistant; $Cm^R$ – chloramphenicol-resistant; $Amp^R$ – ampicillin-resistant.
[a]In this study, pLas16S was used for EMSA in agarose gel with FimT3s.

**Table S7.** List of PCR primers and qPCR primers and probe used in this study.

| Name | Sequence (5'–3') | Purpose or description | Amplicon size (bp) | Source |
|---|---|---|---|---|
| **Site-directed mutagenesis of genes of interest[a]** | | | | |
| pilA2-Up F | CCTCAGGAATCATCCGTAACC | Amplify the upstream region of *pilA2* (PD1926) | 839 | This study |
| pilA2-Up R | **GTCAGCAACACCTTCTTCACGAGGCAGAC**GATGAATCCTTAAATAGCGTTGGTAAG | | | |
| pilA2-Down F | **CATCAGAGATTTTGAGACACAACGTGGC**TTAACACCAGCAACAACACGATTC | Amplify the downstream region of *pilA2* (PD1926) | 909 | This study |
| pilA2-Down R | GGTGATGCCGACAAGATTGGCTG | | | |
| pilA3-Up F | CAGTAGCCCTATCCGTGAATGTGTC | Amplify the upstream region of *pilA3* (PD1077) | 823 | This study |
| pilA3-Up R | **GTCAGCAACACCTTCTTCACGAGGCAGAC**AAAAATTCCCCTAATCTTTGAAAGTG | | | |
| pilA3-Down F | **CATCAGAGATTTTGAGACACAACGTGGC**CCGAACGAGTAATGAGCGCCGATG | Amplify the downstream region of *pilA3* (PD1077) | 859 | This study |
| pilA3-Down R | CCATCACAGACCTATGCGATACTG | | | |
| pilB-Up F | GATAACAACGCAGGCCAAGGTG | Amplify the upstream region of *pilB* (PD1927) | 973 | This study |
| pilB-Up R | **GTCAGCAACACCTTCTTCACGAGGCAGAC**GAAAAGTTCTCTGGTTACTCTGC | | | |
| pilB-Down F | **CATCAGAGATTTTGAGACACAACGTGGC**GCATCAACAACACCTGTTAATGAC | Amplify the downstream region of *pilB* (PD1927) | 1,034 | This study |
| pilB-Down R | CCTTGGTGAATCAGGAGTTGG | | | |
| pilC-Up F | CAGTGCAGCCGGAAGGTCTCAG | Amplify the upstream region of *pilC* (PD1923) | 845 | This study |
| pilC-Up R | **GTCAGCAACACCTTCTTCACGAGGCAGAC**TGCTGTTCTCCCATCCACCGTC | | | |
| pilC-Down F | **CATCAGAGATTTTGAGACACAACGTGGC**AACGTTATGGCATTTCTTGATC | Amplify the downstream region of *pilC* (PD1923) | 835 | This study |
| pilC-Down R | CCATCCACCAACTGCCAGATAAG | | | |
| pilD-Up F | GTATGCGTACGATGGTCAATC | Amplify the upstream region of *pilD* (PD1922) | 958 | This study |
| pilD-Up R | **GTCAGCAACACCTTCTTCACGAGGCAGAC**AACGTTTTATCCAACGACAGAAG | | | |
| pilD-Down F | **CATCAGAGATTTTGAGACACAACGTGGC**GTGAGGTTGGAGTTGATGAGTGTC | Amplify the downstream region of *pilD* (PD1922) | 851 | This study |
| pilD-Down R | GCAACTCATACATCCATACAC | | | |
| pilE1-Up F | CTGTCTGGTGTATTTCCCTGGTC | Amplify the upstream region of *pilE1* (PD0024) | 874 | This study |
| pilE1-Up R | **GTCAGCAACACCTTCTTCACGAGGCAGAC**GGAAGTTACATCATTCACGAACATC | | | |
| pilE1-Down F | **CATCAGAGATTTTGAGACACAACGTGGC**GGTGATCTGATGTTTGGAGTGCTTG | Amplify the downstream region of *pilE1* (PD0024) | 952 | This study |
| pilE1-Down R | CAGGAGTCATCCGTCGTCTTTCG | | | |

| | | | | |
|---|---|---|---|---|
| pilE2-Up F | GCAATCTCTGGAAGTTCAACCTG | Amplify the upstream region of *pilE2* (PD1610) | 912 | This study |
| pilE2-Up R | **GTCAGCAACACCTTCTTCACGAGGCAGAC**ACGTTCCCACTTTAGGATCACCATC | | | |
| pilE2-Down F | **CATCAGAGATTTTGAGACACAACGTGGC**CGTGTTCCCGCATTGCTTTTAGTTG | Amplify the downstream region of *pilE2* (PD1610) | 893 | This study |
| pilE2-Down R | CTCATAGACGAACACGAGTAGG | | | |
| pilF-Up F | CAAGGTTTTAACCGCAATCTGAC | Amplify the upstream region of *pilF* (PD1623) | 855 | This study |
| pilF-Up R | **GTCAGCAACACCTTCTTCACGAGGCAGAC**GACTAAGCAGCCAGATAAAATC | | | |
| pilF-Down F | **CATCAGAGATTTTGAGACACAACGTGGC**TGTATTGTGATCAGTGATTTCG | Amplify the downstream region of *pilF* (PD1623) | 814 | This study |
| pilF-Down R | GTTCCCAGATTCTTGCACTTCCACC | | | |
| pilG-Up F | CAGATAGCGTTGCGCTATTGC | Amplify the upstream region of *pilG* (PD0845) | 869 | This study |
| pilG-Up R | **GTCAGCAACACCTTCTTCACGAGGCAGAC**AGCGCTCTGAATCTAAATACTGTG | | | |
| pilG-Down F | **CATCAGAGATTTTGAGACACAACGTGGC**CCTGACTGTTCATCTGATGCGTTTCC | Amplify the downstream region of *pilG* (PD0845) | 803 | This study |
| pilG-Down R | CAACTGCTGCGACAACACCTG | | | |
| pilH-Up F | GAAGTGATAGTTCGCGCGTTATG | Amplify the upstream region of *pilH* (PD1632) | 1,003 | This study |
| pilH-Up R | **GTCAGCAACACCTTCTTCACGAGGCAGAC**GTCTGCTTCAGCAGTTTAGTGTG | | | |
| pilH-Down F | **CATCAGAGATTTTGAGACACAACGTGGC**GGCACACCATAACGAGAAACCGGAC | Amplify the downstream region of *pilH* (PD1632) | 977 | This study |
| pilH-Down R | GTTATGTTGACTCCCTTCTCTG | | | |
| pilI-Up F | CTACCAGGGTGCACATAATGAAG | Amplify the upstream region of *pilI* (PD0846) | 845 | This study |
| pilI-Up R | **GTCAGCAACACCTTCTTCACGAGGCAGAC**CAGATGAACAGTCAGGTTAAACAG | | | |
| pilI-Down F | **CATCAGAGATTTTGAGACACAACGTGGC**TTGGCGGTCCCGTCTTGCATATTTAG | Amplify the downstream region of *pilI* (PD0846) | 977 | This study |
| pilI-Down R | GTTTATCACGTACCGAGCCAACC | | | |
| pilJ-Up F | GGTGTGAGGTGGTTACCGCTATTG | Amplify the upstream region of *pilJ* (PD0847) | 884 | This study |
| pilJ-Up R | **GTCAGCAACACCTTCTTCACGAGGCAGAC**TCAGGCAGCAGCCTGTCTAAATTCAG | | | |
| pilJ-Down F | **CATCAGAGATTTTGAGACACAACGTGGC**TTGAATGCTTCTCGGCTTGGAAAGG | Amplify the downstream region of *pilJ* (PD0847) | 928 | This study |
| pilJ-Down R | GGAAGCATCGACATGGAGCAATG | | | |

| | | | | |
|---|---|---|---|---|
| pilL-Up F | CGATAGCCGACGCGATTAACTGTG | Amplify the upstream region of *pilL* (PD0848) | 922 | This study |
| pilL-Up R | **GTCAGCAACACCTTCTTCACGAGGCAGAC**TCAAGCTGGCAATTTGAAGTTGGCTG | | | |
| pilL-Down F | **CATCAGAGATTTTGAGACACAACGTGGC**TAGCGTCCAAGGTAAGTTGGTTGC | Amplify the downstream region of *pilL* (PD0848) | 920 | This study |
| pilL-Down R | GATGCCCAGATTATCCCGAAGCAC | | | |
| pilM-Up F | CGTGCATCGGTATTGCTTTTGC | Amplify the upstream region of *pilM* (PD1695) | 1,016 | This study |
| pilM-Up R | **GTCAGCAACACCTTCTTCACGAGGCAGAC**GGGCACTTTTAAGACAGGAACATATC | | | |
| pilM-Down F | **CATCAGAGATTTTGAGACACAACGTGGC**ATGGCCAGAATTAACTTATTGCCCTG | Amplify the downstream region of *pilM* (PD1695) | 987 | This study |
| pilM-Down R | CTTACTAGGCAACTGCCGTAACATC | | | |
| pilN-Up F | GCTTCAAGGTGGAACACTATGCTG | Amplify the upstream region of *pilN* (PD1694) | 994 | This study |
| pilN-Up R | **GTCAGCAACACCTTCTTCACGAGGCAGAC**TCAGTCAAAACTCCTCAAAGCCAGAC | | | |
| pilN-Down F | **CATCAGAGATTTTGAGACACAACGTGGC**ATGAGTAAGAATTCGTTTAAATTGAG | Amplify the downstream region of *pilN* (PD1694) | 1,153 | This study |
| pilN-Down R | GTTCGACGCGATCTTCATTCACC | | | |
| pilO-Up F | CTATGCTGGCAGTGAGTACAAC | Amplify the upstream region of *pilO* (PD1693) | 920 | This study |
| pilO-Up R | **GTCAGCAACACCTTCTTCACGAGGCAGAC**TCAATTGTCCTCTGACAGCG | | | |
| pilO-Down F | **CATCAGAGATTTTGAGACACAACGTGGC**ATGAGTACAAAAACCCTCAAAAAAATAG | Amplify the downstream region of *pilO* (PD1693) | 912 | This study |
| pilO-Down R | GATCTGGCGTTGCAAGACAGTC | | | |
| pilP-Up F | GATAGCAGGGCTCTGCCGTATG | Amplify the upstream region of *pilP* (PD1692) | 853 | This study |
| pilP-Up R | **GTCAGCAACACCTTCTTCACGAGGCAGAC**TCATGGTCCATCCTTCTTGC | | | |
| pilP-Down F | **CATCAGAGATTTTGAGACACAACGTGGC**ATGATTGATTTAAGGGATAG | Amplify the downstream region of *pilP* (PD1692) | 822 | This study |
| pilP-Down R | CAAATTACTTTCTCATCACGGC | | | |
| pilQ-Up F | GAGGTGCTGAAGCAGATGTTAC | Amplify the upstream region of *pilQ* (PD1691) | 967 | This study |
| pilQ-Up R | **GTCAGCAACACCTTCTTCACGAGGCAGAC**TTAATCGTCGAACGAAAGCGC | | | |
| pilQ-Down F | **CATCAGAGATTTTGAGACACAACGTGGC**ATTTCGTCAATGCACTGCTTTATATCTC | Amplify the downstream region of *pilQ* (PD1691) | 897 | This study |
| pilQ-Down R | CAACTCGAAACGCTTGGCACAAC | | | |

| | | | | |
|---|---|---|---|---|
| pilR-Up F | CGTTGACTTTGTTGGCGCTTGG | Amplify the upstream region of *pilR* (PD1928) | 900 | This study |
| pilR-Up R | **GTCAGCAACACCTTCTTCACGAGGCAGAC**GAATGCCAAGATAACGCAGCAGAC | | | |
| pilR-Down F | **CATCAGAGATTTTGAGACACAACGTGGC**TGAGAGGAAGCAGGCAACATAGC | Amplify the downstream region of *pilR* (PD1928) | 937 | This study |
| pilR-Down R | GGTATCGACAAACTCGGTTACG | | | |
| pilS-Up F | CCATCCAGTGCGTACAAATCC | Amplify the upstream region of *pilS* (PD1929) | 924 | This study |
| pilS-Up R | **GTCAGCAACACCTTCTTCACGAGGCAGAC**GTCACGGTGTCCTAAGGCAGAAGAG | | | |
| pilS-Down F | **CATCAGAGATTTTGAGACACAACGTGGC**GTGCCCCGGTATTTCATTCACATTC | Amplify the downstream region of *pilS* (PD1929) | 881 | This study |
| pilS-Down R | GTATGCGCACCGGTGAAACTC | | | |
| pilT-Up F | ACATCATCTGGCACATCAG | Amplify the upstream region of *pilT* (PD1147) | 925 | This study |
| pilT-Up R | **GTCAGCAACACCTTCTTCACGAGGCAGAC**TTGCATAGACGACGAATGG | | | |
| pilT-Down F | **TCAGAGATTTTGAGACACAACGTGGC**TTACCGCACTATCACACATAAG | Amplify the downstream region of *pilT* (PD1147) | 903 | This study |
| pilT-Down R | GCTTGATTGGCGTTGTTG | | | |
| pilU-Up F | GATTCTGGTGACTGGGCCGACTG | Amplify the upstream region of *pilU* (PD1148) | 855 | This study |
| pilU-Up R | **GTCAGCAACACCTTCTTCACGAGGCAGAC**CGGGTGTTGCTCCTTTGCTGCTATG | | | |
| pilU-Down F | **CATCAGAGATTTTGAGACACAACGTGGC**ATCTATTATGAGGTTATATAATC | Amplify the downstream region of *pilU* (PD1148) | 879 | This study |
| pilU-Down R | GTCGATGGTCGCTATGAATTGC | | | |
| pilV1-Up F | GGAACGAAGGAGGTCTCAAAGG | Amplify the upstream region of *pilV1* (PD0020) | 858 | This study |
| pilV1-Up R | **GTCAGCAACACCTTCTTCACGAGGCAGAC**AACAGTGTTTTATTCGTCGTTAGG | | | |
| pilV1-Down F | **CATCAGAGATTTTGAGACACAACGTGGC**GTGAACCGTCGGTTTGCCCTGCAAC | Amplify the downstream region of *pilV1* (PD0020) | 885 | This study |
| pilV1-Down R | CAAGGCAACTCTAACCGCAATGAC | | | |
| pilV2-Up F | GCTGGCAACGTCAAGGAAGAAG | Amplify the upstream region of *pilV2* (PD1614) | 861 | This study |
| pilV2-Up R | **GTCAGCAACACCTTCTTCACGAGGCAGAC**TCATTGACACGCGCCCTTCTCCACC | | | |
| pilV2-Down F | **CATCAGAGATTTTGAGACACAACGTGGC**CGATGTATTCCTCACGCTTAGCGCGTC | Amplify the downstream region of *pilV2* (PD1614) | 911 | This study |
| pilV2-Down R | CAACGGGCACGATACAGTGATG | | | |
| pilW1-Up F | CAGTGCCAACGAGTTAGTTGCTTC | Amplify the upstream region of *pilW1* (PD0021) | 936 | This study |
| pilW1-Up R | **GTCAGCAACACCTTCTTCACGAGGCAGAC**TCACAGTTTACTCCTCACTTCAAAAATTTG | | | |

| | | | | |
|---|---|---|---|---|
| pilW1-Down F | **CATCAGAGATTTTGAGACACAACGTGGC**ATGAGGATGACTCACCTCGGGCCGTACAG | Amplify the downstream region of *pilW1* (PD0021) | 973 | This study |
| pilW1-Down R | CGTAATTGGCTTGAGGCCTTGGAG | | | |
| pilW2-Up F | GCTTATGGTCGTTGTTGCTGTCG | Amplify the upstream region of *pilW2* (PD1613) | 988 | This study |
| pilW2-Up R | **GTCAGCAACACCTTCTTCACGAGGCAGAC**CGTCATAACACACTAGTCAGATTG | | | |
| pilW2-Down F | **CATCAGAGATTTTGAGACACAACGTGGC**TGATGAAGTTACGGGCATTTGTCG | Amplify the downstream region of *pilW2* (PD1613) | 1,069 | This study |
| pilW2-Down R | CATACCCGTCATGCGACACACTAGC | | | |
| pilX1-Up F | GAAGTGAGGAGTAAACTGTGAACC | Amplify the upstream region of *pilX1* (PD0022) | 997 | This study |
| pilX1-Up R | **GTCAGCAACACCTTCTTCACGAGGCAGAC**TCATAAATTTCTGTTCCGTAAGCTG | | | |
| pilX1-Down F | **CATCAGAGATTTTGAGACACAACGTGGC**TTGGATTTCTTCAAGATATCGAAATG | Amplify the downstream region of *pilX1* (PD0022) | 931 | This study |
| pilX1-Down R | CACCATCAATCAAGTAGCTGTG | | | |
| pilX2-Up F | GTAGTGCGTCTGATTGGAATCC | Amplify the upstream region of *pilX2* (PD1612) | 843 | This study |
| pilX2-Up R | **GTCAGCAACACCTTCTTCACGAGGCAGAC**CATCAGTTCCCGTATAAGCGATTG | | | |
| pilX2-Down F | **CATCAGAGATTTTGAGACACAACGTGGC**GCAATGAAAAAAAACGTTCTGAAGTG | Amplify the downstream region of *pilX2* (PD1612) | 849 | This study |
| pilX2-Down R | CGTAACTCATCGGCAACAGAAC | | | |
| pilY1-1-Up F | CGATCATCACCGAATTGCGTGCAG | Amplify the upstream region of *pilY1-1* (PD0023) | 959 | This study |
| pilY1-1-Up R | **GTCAGCAACACCTTCTTCACGAGGCAGAC**TTCGATATCTTGAAGAAATCC | | | |
| pilY1-1-Down F | **CATCAGAGATTTTGAGACACAACGTGGC**CGATGTTCGTGAATGATGTAAC | Amplify the downstream region of *pilY1-1* (PD0023) | 864 | This study |
| pilY1-1-Down R | CTACGCAAACGTGTTGCCATACAG | | | |
| pilY1-2-Up F | GGAGTGGAGTCCTTGCAATTTC | Amplify the upstream region of *pilY1-2* (PD1611) | 988 | This study |
| pilY1-2-Up R | **GTCAGCAACACCTTCTTCACGAGGCAGAC**ATTGAGCGGCGGCACCACGTTGCTC | | | |
| pilY1-2-Down F | **CATCAGAGATTTTGAGACACAACGTGGC**TGGTGATCCTAAAGTGGGAACGTG | Amplify the downstream region of *pilY1-2* (PD1611) | 1,045 | This study |
| pilY1-2-Down R | CGATTCAATGTCCAGCTGCATCC | | | |
| pilY1-3-Up F | GGTGTATTAGGGATAGTGGAGTTC | Amplify the upstream region of *pilY1-3* (PD0502) | 918 | This study |
| pilY1-3-Up R | **GTCAGCAACACCTTCTTCACGAGGCAGAC**CAAATCTTGTTCCTTCATCCTTGCTC | | | |

| | | | | |
|---|---|---|---|---|
| pilY1-3-Down F | **CATCAGAGATTTTGAGACACAACGTGGC**TCCACAAAAAAAGAGATGGATAGAG | Amplify the downstream region of *pilY1-3* (PD0502) | 868 | This study |
| pilY1-3-Down R | GCTTCCACACCTCATTATTCTGTC | | | |
| pilZ-Up F | GAAGGATTAGGGCAACGTGCGGTTTC | Amplify the upstream region of *pilZ* (PD1497) | 883 | This study |
| pilZ-Up R | **GTCAGCAACACCTTCTTCACGAGGCAGAC**TACTCCTTACATTCTGCTCAGCATTC | | | |
| pilZ-Down F | **CATCAGAGATTTTGAGACACAACGTGGC**TTGCTGTTGTGCTTGATGTGTTGCTTG | Amplify the downstream region of *pilZ* (PD1497) | 1,028 | This study |
| pilZ-Down R | GAACGAAGATGCTCTAGCACTGC | | | |
| fimT1-Up F | GCTTGGTGCGTATTACTCAGGCCATC | Amplify the upstream region of *fimT1* (PD0019) | 909 | This study |
| fimT1-Up R | **GTCAGCAACACCTTCTTCACGAGGCAGAC**ATTTAGATACATATGCCACG | | | |
| fimT1-Down F | **CATCAGAGATTTTGAGACACAACGTGGC**AACACTGTTATGTCCATCGCAAGC | Amplify the downstream region of *fimT1* (PD0019) | 879 | This study |
| fimT1-Down R | CAGAATGCGGATTGCGTCTGTTCC | | | |
| fimT2-Up F | CATCGCTGCCGAATCGCCAAATC | Amplify the upstream region of *fimT2* (PD1615) | 880 | This study |
| fimT2-Up R | **GTCAGCAACACCTTCTTCACGAGGCAGAC**CTAAACGCTCCACAGCGTTAATGTC | | | |
| fimT2-Down F | **CATCAGAGATTTTGAGACACAACGTGGC**GTGTCAATGAAAAAGTGTTCCAATC | Amplify the downstream region of *fimT2* (PD1615) | 832 | This study |
| fimT2-Down R | CATCCGCATTGAGAAACAACGAACG | | | |
| fimT3-Up F | GTCTGACCGGTAAGCGGATCGAAC | Amplify the upstream region of *fimT3* (PD1735) | 884 | This study |
| fimT3-Up R | **GTCAGCAACACCTTCTTCACGAGGCAGAC**ACCGATAGTCTCGACAATGTTTAGG | | | |
| fimT3-Down F | **CATCAGAGATTTTGAGACACAACGTGGC**ACACCATCGTATTCACATGACATTTG | Amplify the downstream region of *fimT3* (PD1735) | 900 | This study |
| fimT3-Down R | GCATAAGGATTACGCGCTTTCACC | | | |
| chpB-Up F | GTCAGAGCACCTTTGCAGTTC | Amplify the upstream region of *chpB* (PD0849) | 845 | This study |
| chpB-Up R | **GTCAGCAACACCTTCTTCACGAGGCAGAC**CTTACCTTGGACGCTATCATTCTC | | | |
| chpB-Down F | **CATCAGAGATTTTGAGACACAACGTGGC**TATGTAAGGAAACTACCTTCAATG | Amplify the downstream region of *chpB* (PD0849) | 1,042 | This study |
| chpB-Down R | CAACTACGCTCCGATGAGGTAAAGC | | | |

| | | | | |
|---|---|---|---|---|
| chpC-Up F | CGAATTGATTGCTCGTCGTCAAGG | Amplify the upstream region of *chpC* (PD0850) | 984 | This study |
| chpC-Up R | **GTCAGCAACACCTTCTTCACGAGGCAGAC**TGAAGGTAGTTTCCTTACATATCAATG | | | |
| chpC-Down F | **CATCAGAGATTTTGAGACACAACGTGGC**CATGTGGTGGGATGAATGTGTTTTG | Amplify the downstream region of *chpC* (PD0850) | 857 | This study |
| chpC-Down R | CCAGATTGTCCAGATGGTAAGAG | | | |
| Km-F | GTCTGCCTCGTGAAG | Amplify the kanamycin resistance cassette | 1,202 | [21] |
| Km-R | AAGCCACGTTGTGT | | | |
| **Confirmation of site-directed mutagenesis of genes of interest** | | | | |
| pilA2-Fconf | CACTTTGATCGAGCTGATGATC | Amplify internal sequence of *pilA2* (PD1926) | 328 | This study |
| pilA2-Rconf | GTTGTTATCAGCGATGCGGGTCAAG | | | |
| pilA2Up-Fconf | CAGGTCACTCCATCTGCCTTACC | Amplify internal sequence of upstream region of *pilA2* (PD1926) and the kanamycin resistance cassette | 972 | This study |
| pilA3-Fconf | CAAGGCTTCACCCTGTTAGAGG | Amplify internal sequence of *pilA3* (PD1077) | 410 | This study |
| pilA3-Rconf | CTTGAGCCATGCAAACCTACATCG | | | |
| pilA3UP-Fconf | CGTCAATAACGCGAACAGCATC | Amplify internal sequence of upstream region of *pilA3* (PD1077) and the kanamycin resistance cassette | 1,119 | This study |
| pilB-Fconf | CTACTGGATGTGTCCGCATTCG | Amplify internal sequence of *pilB* (PD1927) | 493 | This study |
| pilB-Rconf | CAGCCTTCAGCAGACCATCAATCC | | | |
| pilBUp-Fconf | CCATCCATGCAGCGTTTCCATCTG | Amplify internal sequence of upstream region of *pilB* (PD1927) and the kanamycin resistance cassette | 1,130 | This study |
| pilC-Fconf | CATCACGCCTCTTGTGGTTAAG | Amplify internal sequence of *pilC* (PD1923) | 649 | This study |
| pilC-Rconf | CAACACCAACCTATCCAACAG | | | |
| pilCUp-Fconf | GTTGTACTCTTCCCATCAGCAATC | Amplify internal sequence of upstream region of *pilC* (PD1923) and the kanamycin resistance cassette | 1,096 | This study |
| pilD-Fconf | GGAAAACATTCCGGTGCTTAGC | Amplify internal sequence of *pilD* (PD1922) | 500 | This study |
| pilD-Rconf | CAAGCCAAACAGAGCCCAATACC | | | |
| pilDUp-Fconf | CCAACTTTATGCAGCGGTGGTG | Amplify internal sequence of upstream region of *pilD* (PD1922) and the kanamycin resistance cassette | 1,100 | This study |

| | | | | |
|---|---|---|---|---|
| pilE1-Fconf | GATGATTGTGGTGGTGATCGTG | Amplify internal sequence of *pilE1* (PD0024) | 305 | This study |
| pilE1-Rconf | GTGTGAGCCTGCCACAGTGATC | | | |
| pilE1Up-Fconf | CCAGGTACAGATTGTTGATGAG G | Amplify internal sequence of upstream region of *pilE1* (PD0024) and the kanamycin resistance cassette | 1,044 | This study |
| pilE2-Fconf | GTTGATGGTTGTGGTCGCTGTC | Amplify internal sequence of *pilE2* (PD1610) | 301 | This study |
| pilE2-Rconf | GCACTTGTCGTTTTGCTGAGTG C | | | |
| pilE2Up-Fconf | GGTATGTCGATTTGGTGGTACA G | Amplify internal sequence of upstream region of *pilE2* (PD1610) and the kanamycin resistance cassette | 1,024 | This study |
| pilF-Fconf | GTTTACAACGTGCGTGATGATG | Amplify internal sequence of *pilF* (PD1623) | 388 | This study |
| pilF-Rconf | CACAACCGCCGGAATTGGCTAA AG | | | |
| pilFUp-Fconf | CAAACTAATGCGGCAATTTGAC | Amplify internal sequence of upstream region of *pilF* (PD1623) and the kanamycin resistance cassette | 878 | This study |
| pilG-Fconf | GTCTTAGGGTGATGGTCATTG | Amplify internal sequence of *pilG* (PD0845) | 305 | This study |
| pilG-Rconf | CAGATATTGTTCGGAACCAACC AC | | | |
| pilGUp-Fconf | CAGCATTGACTGGAGGATCTTT AC | Amplify internal sequence of upstream region of *pilG* (PD0845) and the kanamycin resistance cassette | 1,042 | This study |
| pilH-Fconf | GCATTTTGATCGTCGAGGACTC | Amplify internal sequence of *pilH* (PD1632) | 313 | This study |
| pilH-Rconf | GAGAAAGGCTTGGTGATGTAGG | | | |
| pilHUp-Fconf | CTGGCTTTGGACATCTTCAGTG | Amplify internal sequence of upstream region of *pilH* (PD1632) and the kanamycin resistance cassette | 1,100 | This study |
| pilI-Fconf | GTGTTGGCTATCGTATTGGAAG | Amplify internal sequence of *pilI* (PD0846) | 252 | This study |
| pilI-Rconf | CGATAACCAGTGCTGCATTATC | | | |
| pilIUp-Fconf | GCGTCTGCTATTGCAGTACACA G | Amplify internal sequence of upstream region of *pilI* (PD0846) and the kanamycin resistance cassette | 1,021 | This study |
| pilJ-Fconf | GCTGATCGCTTCGTGAGTAATG TG | Amplify internal sequence of *pilJ* (PD0847) | 557 | This study |
| pilJ-Rconf | CCAACAATCGACACTACAGACA GC | | | |

| | | | | |
|---|---|---|---|---|
| pilJUp-Fconf | CCTGACTGTTCATCTGATGCGTTTC | Amplify internal sequence of upstream region of *pilJ* (PD0847) and the kanamycin resistance cassette | 1,072 | This study |
| pilL-Fconf | GCTTGATGAAGAGCAGGGTAATC | Amplify internal sequence of *pilL* (PD0848) | 464 | This study |
| pilL-Rconf | GGATTTCACGCAGTACACTGTCC | | | |
| pilLUp-Fconf | CGAGATTGCGGCTAGTATCGAAC | Amplify internal sequence of upstream region of *pilL* (PD0848) and the kanamycin resistance cassette | 1,236 | This study |
| pilM-Fconf | GCAGGATTTGGAAGCTCAGATAG | Amplify internal sequence of *pilM* (PD1695) | 408 | This study |
| pilM-Rconf | CCATAGCGATGCATTACTTCGTC | | | |
| pilMUp-Fconf | GTCAATGCCGCTGTGTTCGTAG | Amplify internal sequence of upstream region of *pilM* (PD1695) and the kanamycin resistance cassette | 1,049 | This study |
| pilN-Fconf | GGTCAGTGGTCAGAATGATCG | Amplify internal sequence of *pilN* (PD1694) | 439 | This study |
| pilN-Rconf | CTTATCTGGCGACTCTGGCTTGG | | | |
| pilNUp-Fconf | CAACAACACGGAGATGGTCCAAG | Amplify internal sequence of upstream region of *pilN* (PD1694) and the kanamycin resistance cassette | 1,188 | This study |
| pilO-Fconf | GTCATCATGGCTTGGTTCCTC | Amplify internal sequence of *pilO* (PD1693) | 377 | This study |
| pilO-Rconf | GGTAAAGAAGCCACACCACTGAC | | | |
| pilOUp-Fconf | CACATTGGTGGGTATCCTCGTTTC | Amplify internal sequence of upstream region of *pilO* (PD1693) and the kanamycin resistance cassette | 1,105 | This study |
| pilP-Fconf | GTCGGTTTGATTGTTCTCTTGG | Amplify internal sequence of *pilP* (PD1692) | 433 | This study |
| pilP-Rconf | GTTCGACGCGATCTTCATTCAC | | | |
| pilPUp-Fconf | GCAGCCATTAAAGCAGCAGTTG | Amplify internal sequence of upstream region of *pilP* (PD1692) and the kanamycin resistance cassette | 970 | This study |
| pilQ-Fconf | CGTATCGACGCTAAGCCTATGG | Amplify internal sequence of *pilQ* (PD1691) | 567 | This study |
| pilQ-Rconf | CATACCTTTGGCTTCTGTCAGTGC | | | |

| | | | | |
|---|---|---|---|---|
| pilQUp-Fconf | GAGAATCGTACTGCTGATGCTG | Amplify internal sequence of upstream region of *pilQ* (PD1691) and the kanamycin resistance cassette | 1,096 | This study |
| pilR-Fconf | CGTATCAGAACGGCAGCTAATCTG | Amplify internal sequence of *pilR* (PD1928) | 400 | This study |
| pilR-Rconf | CTTGACTACGCGCCACTTTAGC | | | |
| pilRUp-Fconf | CGATGAATACCGCCAGACATTG | Amplify internal sequence of upstream region of *pilR* (PD1928) and the kanamycin resistance cassette | 1,137 | This study |
| pilS-Fconf | CTTGTGGAATACATCTGGACTG | Amplify internal sequence of *pilS* (PD1929) | 642 | This study |
| pilS-Rconf | CCTTTGATCGCCTTTGTCAAG | | | |
| pilSUp-Fconf | CTGTTTCACTTCGTCGGTTGTC | Amplify internal sequence of upstream region of *pilS* (PD1929) and the kanamycin resistance cassette | 992 | This study |
| pilT-Fconf | GTGATGACATTGGACGAACTCG | Amplify internal sequence of *pilT* (PD1147) | 719 | This study |
| pilT-Rconf | CTTTATCCTTGGCGTATTCACG | | | |
| pilTUp-Fconf | CCTTCTTTACTATCTCCTGAGC | Amplify internal sequence of upstream region of *pilT* (PD1147) and the kanamycin resistance cassette | 951 | This study |
| pilU-Fconf | CTCAATGGCATCAATGATCAAC | Amplify internal sequence of *pilU* (PD1148) | 365 | This study |
| pilU-Rconf | GATTCAGCGACAGATCCATCAG | | | |
| pilUUp-Fconf | GATCATCTCGCAAATGTTGTTG | Amplify internal sequence of upstream region of *pilU* (PD1148) and the kanamycin resistance cassette | 940 | This study |
| pilV1-Fconf | CGTTAGTTTGATCGAAGTGCTG | Amplify internal sequence of *pilV1* (PD0020) | 323 | This study |
| pilV1-Rconf | CATCACAAGAGATCGAACCACAG | | | |
| pilV1Up-Fconf | GAACGAAGGAGGTCTCAAAGGAAAGC | Amplify internal sequence of upstream region of *pilV1* (PD0020) and the kanamycin resistance cassette | 1,353 | This study |
| pilV2-Fconf | GTGTTCCAATCATCGTTGCCATTG | Amplify internal sequence of *pilV2* (PD1614) | 381 | This study |
| pilV2-Rconf | CCATTCGTCGCAAGATTCACC | | | |
| pilV2Up-Fconf | CAGACAGTGTTTGCGCTGCACTG | Amplify internal sequence of upstream region of *pilV2* (PD1614) and the kanamycin resistance cassette | 1,111 | This study |

| | | | | |
|---|---|---|---|---|
| pilW1-Fconf | GAGATTCCGCCACAGAATGGTC | Amplify internal sequence of *pilW1* (PD0021) | 405 | This study |
| pilW1-Rconf | CATACCACTGCACTGCACGCAATTC | | | |
| pilW1Up-Fconf | CCAACACACGGATGGCTAACAAC | Amplify internal sequence of upstream region of *pilW1* (PD0021) and the kanamycin resistance cassette | 1,124 | This study |
| pilW2-Fconf | CGTCATGGACTGCTTACCGTCTTTC | Amplify internal sequence of *pilW2* (PD1613) | 408 | This study |
| pilW2-Rconf | GCAGTACCAGAACATCACTTCC | | | |
| pilW2Up-Fconf | CAGTTGTTCGTGTCAGTGAAGG | Amplify internal sequence of upstream region of *pilW2* (PD1613) and the kanamycin resistance cassette | 1,189 | This study |
| pilX1-Fconf | GCTAACCTCCGTGATCGCAGTTTG | Amplify internal sequence of *pilX1* (PD0022) | 409 | This study |
| pilX1-Rconf | CATTGCTAGACGGATCGCTGCTTCG | | | |
| pilX1Up-Fconf | CTCAAGGCGTCAGAGGAACAGAC | Amplify internal sequence of upstream region of *pilX1* (PD0022) and the kanamycin resistance cassette | 1,148 | This study |
| pilX2-Fconf | GTTGTGCTGGTGGTGTTGGTG | Amplify internal sequence of *pilX2* (PD1612) | 350 | This study |
| pilX2-Rconf | CTGAATACAGTGATACGGAAC | | | |
| pilX2Up-Fconf | GTTTGAACGAGCGCGGGATTCC | Amplify internal sequence of upstream region of *pilX2* (PD1612) and the kanamycin resistance cassette | 940 | This study |
| pilY1-1-Fconf | GCTAATCTGGCCGACACCTACAC | Amplify internal sequence of *pilY1-1* (PD0023) | 479 | This study |
| pilY1-1-Rconf | GAATCGTAAGGTTGGCCCTCTGTG | | | |
| pilY1-1Up-Fconf | CTGTTCTGCGCACGACGTTACTTC | Amplify internal sequence of upstream region of *pilY1-1* (PD0023) and the kanamycin resistance cassette | 1,042 | This study |
| pilY1-2-Fconf | CACATCGCGTATGGATGCTTTGC | Amplify internal sequence of *pilY1-2* (PD1611) | 541 | This study |
| pilY1-2-Rconf | CTGGTATGAGCCAGTCAACAGAC | | | |
| pilY1-2Up-Fconf | CAAGTTGCAACCATGCAGGAG | Amplify internal sequence of upstream region of *pilY1-2* (PD1611) and the kanamycin resistance cassette | 1,044 | This study |

| | | | | |
|---|---|---|---|---|
| pilY1-3-Fconf | CAGTGATGGATCGTGCTTATCC | Amplify internal sequence of *pilY1-3* (PD0502) | 552 | This study |
| pilY1-3-Rconf | GAGTAGTCGCCTTAGCATCCTCAG | | | |
| pilY1-3Up-Fconf | GCATCTCTGGATTGTCAGCATCG | Amplify internal sequence of upstream region of *pilY1-3* (PD0502) and the kanamycin resistance cassette | 1,187 | This study |
| pilZ-Fconf | CAGGGTATTCTGTCGCTCACC | Amplify internal sequence of *pilZ* (PD1497) | 326 | This study |
| pilZ-Rconf | GTGTGCGTTGGCTTATCTGAG | | | |
| pilZUp-Fconf | GCATTGCTGAAGACGCTTGAG | Amplify internal sequence of upstream region of *pilZ* (PD1497) and the kanamycin resistance cassette | 1,088 | This study |
| fimT1-Fconf | CAATTCTTGCGGGCCTTGCTTATC | Amplify internal sequence of *fimT1* (PD0019) | 396 | This study |
| fimT1-Rconf | GGTTGTTAGCCATCCGTGTGTTGG | | | |
| fimT1Up-Fconf | CAAAGGTGTTGATGACCCGATG | Amplify internal sequence of upstream region of *fimT1* (PD0019) and the kanamycin resistance cassette | 1,088 | This study |
| fimT2-Fconf | GCTTATGGTCGTTGTTGCTGTCG | Amplify internal sequence of *fimT2* (PD1615) | 367 | This study |
| fimT2-Rconf | GTTCGATGCACGCCGTAAGCCATGC | | | |
| fimT2Up-Fconf | GTGAAGAACGCAAGGATGCTTATTAC | Amplify internal sequence of upstream region of *fimT2* (PD1615) and the kanamycin resistance cassette | 1,014 | This study |
| fimT3-Fconf | CACGATGGCGCTAATCGCATTATTG | Amplify internal sequence of *fimT3* (PD1735) | 398 | This study |
| fimT3-Rconf | CATTGTTGCAGAAGCGGATGCTG | | | |
| fimT3Up-Fconf | CAACACATCAGAACGTGACAGC | Amplify internal sequence of upstream region of *fimT3* (PD1735) and the kanamycin resistance cassette | 1,090 | This study |
| chpB-Fconf | GATTGCTCGTCGTCAAGGTTG | Amplify internal sequence of *chpB* (PD0849) | 569 | This study |
| chpB-Rconf | GATCAACTGGCAAAGTGGAGACTC | | | |
| chpBUp-Fconf | CACTGGTGATGGTTGTGTGATTG | Amplify internal sequence of upstream region of *chpB* (PD0849) and the kanamycin resistance cassette | 1,034 | This study |

| | | | | |
|---|---|---|---|---|
| chpC-Fconf | CGAACGAGTGCTATTACCAAATGC | Amplify internal sequence of *chpC* (PD0850) | 357 | This study |
| chpC-Rconf | GCTTGAGTTTCACTAAGAAGTACAC | | | |
| chpCUp-Fconf | GCATTACCCGCTGGTTTCAAACATC | Amplify internal sequence of upstream region of *chpC* (PD0850) and the kanamycin resistance cassette | 1,020 | This study |
| Km-Rconf | GATTCAGTCGTCACTCATGGTG | Pairs with "Up-Fconf" primers to amplify internal sequence of upstream region of the gene of interest and the kanamycin resistance cassette | - | [93] |
| **Confirmation of pAX1-Cm recombination into the NS1 region of *X. fastidiosa* TemeculaL** | | | | |
| Cm-F | AATCAGCGACACTGAATACGG | Amplify the chloramphenicol resistance cassette | 1,119 | [21] |
| Cm-R | TCACTTATTCAGGCGTAGCAC | | | |
| **qPCR primers and probe used to quantify the *X. fastidiosa* population** | | | | |
| HL5 Forward | AAGGCAATAAACGCGCACTA | Quantify the *X. fastidiosa* population as genome equivalents | 221 | [84] |
| HL6 Reverse | GGTTTTGCTGACTGGCAACA | | | |
| HLp (probe) | 6FAM-TGGCAGGCAGCAACGATACGGCT-QSY | | | |
| **Cloning into pHIS-Parallel1[b]** | | | | |
| fimT1s-pHIS-F | CATG**CCATGG**ATAATGGTGAGCGCTTG (NcoI) | Amplify *fimT1s* to clone into pHIS-Parallel1 | 459 | This study |
| fimT1s-pHIS-R | CCG**CTCGAG**TTATTCGTCGTTAGG (XhoI) | | | |
| fimT2s-pHIS-F | CATG**CCATGG**ATCGCTCGAATCGAGTG (NcoI) | Amplify *fimT2s* to clone into pHIS-Parallel1 | 429 | This study |
| fimT2s-pHIS-R | CCG**CTCGAG**TCATTGACACGC (XhoI) | | | |
| fimT3s-pHIS-F | CATG**CCATGG**ATAAACAAACTCTGGAAC (NcoI) | Amplify *fimT3s* to clone into pHIS-Parallel1 | 519 | This study |
| fimT3s-pHIS-R | CCG**CTCGAG**TTATTGGAATGGTGTTTC (XhoI) | | | |
| **Amino acid exchanges in FimT3s (cloned into pHIS-Parallel1)** | | | | |
| FimT3_R160A R162A-F | GAGTATTAAGTATCCGTTGCGAGGCAATTGCGCCTAAATTGTTGATCACGATGA | Exchange the arginine amino acid residues at positions 160 and 162 of FimT3s cloned into pHIS-Parallel1 by alanine residues | 5,859 | This study |
| FimT3_R160A R162A-R | TCATCGTGATCAACAATTTAGGCGCAATTGCCTCGCAACGGATACTTAATACTC | | | |
| **Sanger sequencing** | | | | |
| T7 promoter - forward | TAATACGACTCACTATAGGG | Amplify the gene cloned downstream of the T7 promoter in pHIS-Parallel1 for Sanger sequencing | variable | Eurofins Genomics LLC |

[a]Nucleotides in bold indicate the 5' extended region of each primer that is homologous to the kanamycin resistance cassette sequence to allow fusion of chromosomal sequences to the marker by overlap-extension PCR.
[b]Nucleotides in bold indicate the restriction sites of the endonucleases (specified within parenthesis) used for cloning into pHIS-Parallel1.
